# Towards a comprehensive chemical and genetic tool library for rhamnogalacturonan-II oligosaccharides and exploitation

**DOI:** 10.64898/2026.03.13.711244

**Authors:** Didier Ndeh, Sergey Nepogodiev, Rodrigo Marcias-Garbette, Immacolata Venditto, Kalimullah Saighani, Anastasiia Kalachikova, Colin Ruprecht, Markus Blaukopf, Carmen Escudero-Martinez, Girma Dinsa, Abdelmadjid Atrih, Ian Lidbury, Nicolas Terrapon, Bernard Henrissat, Marie-Christine Ralet, Fabian Pfrengle

## Abstract

Rhamnogalacturonan-II (RG-II) is considered the most complex glycan in nature. It forms part of an intricate network of complex glycans in the plant cell wall where it plays a critical role in plant growth, development and defence. It has been identified as an important nutrient source for the human gut microbiota (HGM), a key modulator of human health and disease status. Increasing evidence also suggests that RG-II can modulate plant-microbe interactions. Given its importance and potential, detailed studies of RG-II’s structure-function relationships and metabolism are required to underpin future crop*-*improvement strategies and to harness its benefits for plant and human health. Progress in this field is however hampered by RG-II’s structural complexity and limited access to enabling tools, in particular chemically defined RG-II-derived oligosaccharide (CDRO) substructures. Achieving targeted, efficient, and scalable production of CDROs remains a significant challenge and is indeed one of the major reasons why RG-II and glycomic research in general, significantly lag behind genomic and proteomic research. Here, we have genetically engineered as well as screened a diverse set of genetic strains, including transposon (Tn) mutants of the prominent model human gut microbe *Bacteroides thetaiotaomicron* (*B. theta*) and its gut and plant-associated relatives for new CDRO-generating and/or RG-II-utilising strains. Several CDROs, some of which had never been produced before by any other means (including chemical synthesis), where generated and characterised by a combination of high-resolution mass spectrometry (MS), enzymatic profiling and 2D-NMR. In addition to expanding the CDRO toolbox, we identified key genetic strains that will serve as a base or platform for the production of an unprecedented amount of CDROs covering the complexity and diversity of chemical modifications in RG-II. CDROs were later exploited to gain new insights into the microbial metabolism of RG-II in the human gut, revealing key aspects of its chemical structure that drive or limit its metabolism in *B. theta*. Notably, we generated new evidence in support of an alternative operational paradigm for polysaccharide utilisation systems that are widespread in the Bacteroidota phylum. We confirmed the presence of pathways for the metabolism of RG-II and/or RG-II core sugars _D-_apiose (_D-_Api*f*), and 3-deoxy-_D-_manno-2-octulosonic acid (_D-_Kdo) in aerobic plant-associated microbes including fungi and *Flavobacterium spp.*, highlighting their potential to be exploited as cost-effective alternatives to *B. theta* for the generation of CDROs.

## Introduction

Plant cell wall glycans play critical roles in several physiological and pathophysiological processes in the plant, including immunity, cell signalling, growth and development, and maintenance of cell wall architecture^1,2^. They are also a major source of nutrients for the human gut microbiota which is known to impact human health and disease status^3^. Plant cell wall glycans include cellulose, hemicelluloses, arabinogalactans, proteoglycans, and pectins. The pectic network consists of three major structural units including homogalacturonan (HG), rhamnogalacturonan I (RG-I) and rhamnogalacturonan II (RG-II)^4^. RG-II is the most structurally complex polysaccharide in nature with a high number of diverse glycosidic linkages and rare and complex monosaccharides such as 3-deoxy-_D-_lyxo-2-heptulosaric acid (_D-_Dha), _L-_aceric acid (_L-_AceA*f*), 2-O-methy_L-L-_fucose (2-O-Me-_L-_Fuc*p*), 2-O-methy_L-_xylose (2-O-Me-_D-_Dxyl*p*), _D-_apiose (_D-_Api*f*), and 3-deoxy-_D-_manno-2-octulosonic acid (_D-_Kdo)^5^ (**Fig. 1A**). It exists as a large dimer whose monomeric units (∼4.5 kDa each) are held together by the micronutrient boron through diester bonds^6,7^. Each monomeric unit consists of a linear backbone of HG, containing α-(1-4)-linked _D-_galacturonic acid (_D-_GalA*p*) residues to which are attached several variable side chains A to F (**Fig. 1A**, <u>5 **1**</u>). Although largely conserved near the core in many plants, recent mass spectrometric (MS) evidence suggests that peripheral regions of RG-II exhibit more structural variability than previously thought^8–11^. Variations in the structure have not only been detected between plants but also between organs and different developmental stages within the same plant^8–10^. Variably methylated, acetylated or glycosylated forms of one or more side chains have been reported, modifications which significantly add to the complexity and diversity of variants of an already highly complex structure in different plants^5,8–10^

**Fig. 1:**
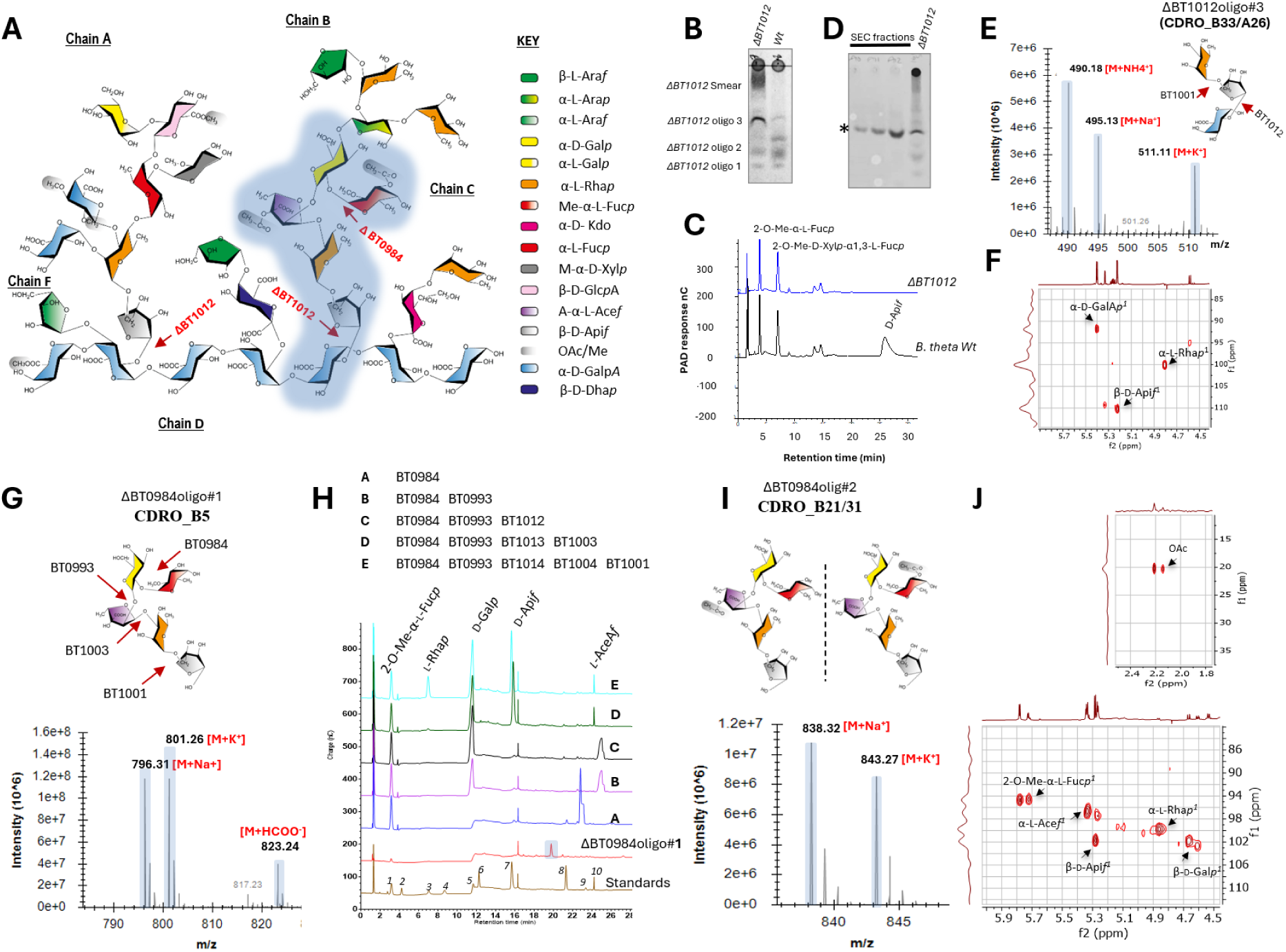
Production, purification and characterisation of chemically defined RG-II-derived oligosaccharides (CDROs) of ΔBT1012 and ΔBT0984. Data shows an in-depth characterisation of CDROs generated by the genetic strains ΔBT1012 and ΔBT0984, setting a stage for the production of diverse D-GalA*p*-linked CDROs a: Structure of apple RG-II showing glycosidic linkages targeted by BT1012 and BT0984 CAZymes. Highlighted region shows D-GalA*p*-linked CDRO targeted for production (also see Fig. 2) b: TLC comparison of the secretion product profile of ΔBT1012 and *B. theta* Wt after growth on RG-II as sole carbon source c: HPAEC-PAD analyses of samples in b. d: SEC and TLC analyses of purified ΔBT1012 sugars (from b.). Star shows position of a dominant and well purified sugar ΔBT1012 oligo #3 (CDRO_B33/A26) e: high resolution MS detection of CDRO_B33/A26 and its ionic adducts f: 2D NMR (^1^H,^13^C-HSQC) analyses of purified CDRO_B33/A26 g: MS analyses of ΔBT0984oligo#1 (CDRO_B5) showing various ionic adducts h: Enzymatic profiling of CDRO_B5 using RG-II specific enzymes. Standards: 2-O-Me-α-L-Fuc*p* (1), D-Fuc*p* (2), L-Rha*p* (3), L-Arap (4), D-Galp (5), D-Xylp (6), D-Apif (7), D-kdo (8), D-Gal*p* (9), L-AceA*f* (10) i: MS confirming the detection of acetylated variants of CDRO_B5. j: 2D NMR (^1^H,^13^C-HSQC) analyses of the structure of purified CDRO_B21/31 showing the detection of acetyl groups and other components.

Increasing evidence shows that RG-II is critical for plant growth and development, as mutations disrupting its structure or borate cross-linking cause developmental defects, lethality, and reduced crop yields^12–14^. Alterations to the structure of RG-II have been associated with a diminished capacity to withstand environmental stressors, such as high salinity and freezing temperatures^15,16^. RG-II components such apiose have been shown to be targeted by plant pathogens such as *Pectobacterium carotovorum, Pectobacterium atrosepticum* and *Rhizobium rhizogenes* which are responsible for significant crop losses in agriculture^17^. We and others have also recently showed that RG-II is an important nutrient source for the human gut microbiota (HGM)^5,18^, a key modulator of human health and disease status^3,19^. HGMs role in human health is reinforced by recent evidence highlighting strong links between its composition and important human diseases such as obesity, cancer, diabetes, autoimmune arthritis and inflammatory bowel disease^20–24^. We unravelled the pathway for the enzymatic deconstruction of RG-II in the model HGM species *B. theta*, identifying enzymes cleaving all but one of the 21 glycosidic linkages in this glycan^5^. These enzymes were found to be clustered in genetic loci termed polysaccharide utilisation loci (PULs) which are widespread in *B. theta*’s genome and known to encode cell envelo*p-*associated multiprotein systems that coordinate glycan capture, import and utilisation^25–29^.

Given RG-II’s importance in modulating plant growth, development, pathogen interactions, and potentially human health, detailed molecular studies of its metabolism are essential to underpin future cro*p-*improvement strategies and to harness its benefits for both plant and human health.

Progress in this field has been hindered by the structural complexity of RG-II and limited access to enabling tools, in particular chemically defined RG-II derived oligosaccharide (CDRO) substructures^30,31,32^. To date a majority of RG-II biosynthetic and degradative enzymes in plants remain unknown^7,33,34^. How the translocation or transport of RG-II and its components occurs in the plant and what proteins are involved also remains unclear. Our understanding of RG-II metabolism in the gut also remains limited as several elements of the RG-II transcriptome in *B. theta* remain uncharacterised. Additionally, how the microbe coordinates the extracellular capture, signalling and import of RG-II remains unclear. These setbacks limit our ability to unlock the full potential of RG-II. Extensive collections of CDROs representing the diversity of structural modifications in RG-II and easy access to CDRO-generating systems will be important for achieving these goals by facilitating the screening and identification of various proteins or enzymes involved in these processes. Critically, they will facilitate the development of more advance tools such as glycan arrays and RG-II specific monoclonal antibodies to track changes in plant cell wall. It is also worth noting that, as bioactive molecules, chemically defined oligosaccharides and their conjugates in general, have in the past been exploited in the pharmaceutical, food and biotech industries as drugs, drug delivery agents, prebiotics/nutraceuticals and as vaccine components^35–39^

As expected, RG-II’s role and potential has prompted several chemical synthetic efforts targeting CDROs, however, this has only been achieved for a very small number of side chain A and B-derived CDROs^35–39^. Notably, pure acetylated, unacetylated, methylated and homogalacturonan (HG) backbone-containing CDROs are rare^5,30^ and no CDROs to date are commercially available. These diverse chemical configurations or modifications in the CDROs are known to impact the recognition or specificities of RG-II several interacting proteins^5^. Indeed, in a recent study, the β-glucuronidase BT0996 that cleaves the terminal _D-_GlcA*p* residue in side chain A of RG-II was shown to contain a new CBM family (BT0996-C) that strangely shows specificity for the HG backbone which is several monosaccharide units away from _D-_GlcA*p*^45^. Refinement of BT0996-C specificity was however hampered by limited access to unique HG backbone-containing CDROs of side chain A oligos^45^ further highlighting the importance of the availability of a diverse and easily accessible repertoire of pure CDRO glycoforms. Given the key location of HG backbone (at the base or reducing ends of all RG-II side chains), it is highly likely that potential acceptor ligands for the majority of RG-II biosynthetic will contain terminal _D-_GalA*p* or HG-oligosaccharide units. In light of this, only recently and for the first time, a fully deprotected CDRO pentasacharide containing a HG-tetrasaccharide linked to single _D-_Kdo unit (derived from the structure of side chain C) was chemically produced using a post-glycosylation oxidation and stereoselective glycosylation with a _D-_Kdo fluoride donor^46^.

Chemical approaches have limitations in that they often require complex chemical reactions and expert knowledge of carbohydrate chemistry. Many of the resulting glycans also sometimes contain additional or unwanted chemical groups or unnatural modifications^47^ which can impact their interactions especially for highly specific enzymes. Chemical approaches also have issues of cost, safety and limited scale of production especially with very complex structures. Highlighting the importance of CDROs in plant research, Ruprecht *et al.* developed and demonstrated the use of a glycan array for the high-throughput screening of plant biosynthetic enzymes^48,49^. However, this array lacked CDROs due to the difficulty associated with their synthesis. Lastly, data from our detailed characterisation of RG-II, determined that critical components of the structure including a core rhamnose anomer (α-_L-_Rha) had been misannotated. Although, this led to the revision of the RG-II structure, it set us several years behind as some chemically synthesised CDROs carried the incorrect anomer^41,50^.

Our recent work identifying a range of RG-II specific enzymes has also greatly facilitated the *in-vitro* production of certain CDROs, in particular those detached from the backbone _D-_GalA*p/*HG unit of RG-II^5^. However, for such a large and highly complex glycan like RG-II, deploying tens of recombinant enzymes to derive specific CDROs is very laborious, time-consuming and resource-intensive. Buffer incompatibility issues due the requirement of complex enzymatic mixtures, frequent cross-contamination from some highly active enzymes leading to the loss of target CDROs, poor expression and *in-vitro* activity of some recombinant proteins and stability issues are often encountered. Under these circumstances, therefore, readily available engineered and customised systems for the scalable production of specific CDROs are far more desirable.

Recently we determined that *B. theta* deletion mutants defective in specific RG-II degradative pathways could be exploited to generate scalable quantities of specific CDROs. The success of this approach in generating scalable quantities of over 12 structurally diverse and complex CDROs including complex _D-_GalA*p/*HG-containing CDROs derived from side chain A suggests that it has huge potential as a major alternative to chemical and *in vitro* chemo-enzymatic synthesis routes^5^. Our estimations, based on the complexity and chemical modifications in RG-II show that more than 200 different or specific CDROs including acetylated, unacetylated and _D-_GalA*p/*HG-containing CDROs can be derived from RG-II, yet only a very small amount (<10%) have not been produced before or have readily available tools for their production^5,30^ (**Supplemental table 1**)

Here, we engineered new *B. theta* CDRO-generating strains and screened *B. theta* transposon libraries for more CDRO-generating strains. Other human gut microbes as well as plant-associated microbes were also investigated for their ability to metabolise RG-II and its constituent sugars and generate CDROs. We identify new CDRO-generating strains, successfully purified and characterised several CDROs and generated important data which set a stage for the production of an unprecedented amount of new CDROs covering the complexity and diversity of sugar modifications in RG-II. We further exploited various CDROs to gain new insights into the metabolism of RG-II in the prominent human gut microbe *B. theta*. Notably, we identified key drivers of RG-II uptake during its metabolism in *B. theta* and provide evidence supporting the existence of an alternative operational paradigm for polysaccharide utilisation systems (PUS) that are widespread in the phylum Bacteroidota^51–55^. Our work also allowed us to gain new insights into RG-II-microbe interactions out of the human gut environment, specifically plant-associated microbes, revealing that some of them in particular, *Flavobacterium spp.* possess just as much RG-II metabolic capacity as their gut counterparts and hence could potentially be exploited in a similar manner for the production of CDROs.

## Results

### Targeted production of diverse _D-_GalA*p-*linked CDROs of RG-II side chain B

#### B. theta ΔBT1012 as a starting platform

Our analyses based on genetic and biochemical data from these studies and previous suggest that more than 200 CDROs can be derived from RG-II by virtue of its intricate chemical complexity and extensive branching (**Fig. 1A, Supplemental table 1, Supplemental table 2, Supplemental figure 1**), yet the majority of these sugars are not accessible, due to lack of knowledge and challenging chemical procedures for their synthesis. Notably, none of the more than 100 _D-_GalA*p*/poly-_D-_GalA*p-*linked side chain B CDROs in the RG-II sequence space in **Supplemental table 1** (**CDRO_B26-B125, CDRO_D/B1-25, CDRO_M10-25**) have been produced or synthesised before by any other means including chemical synthesis. Here we show that a unique genetic strain ΔBT1012, defective in RG-II metabolism can be tailored to produce an unprecedented number of these CDROs derived from various side chains in apple RG-II. ΔBT1012 lacks the β-_D-_apiosidase BT1012 that cleaves the _D-_Api*f*-β1,2-_D-_GalA*p* glycosidic linkage connecting side chains A & B to the RG-II poly-_D-_GalA*p* backbone **(Fig. 1A)**, and hence by creating this strain, we anticipated that this critical linkage will be retained during RG-II metabolism, leading the generation of diverse _D-_Gal*p*A-linked CDROs. Thin layer chromatographic (TLC) analyses of ΔBT1012 products showed that the mutation caused metabolic defects in the strain, evident from the accumulation of several RG-II degradation products appearing as a smear of sugars (ΔBT1012smear) and distinct bands (ΔBT1012oligo#1, 2, 3) compared to the wild type *B. theta* strain (Wt) (**Fig. 1B**). High performance anion exchange chromatography with pulse amperometric detection (HPAEC-PAD) of the secreted products showed a complete loss of the sugar _D-_Api*f* in the ΔBT1012 strain compared to Wt (**Fig. 1C)**, a strong indication that the _D-_Api*f*-β1,2-_D-_GalA*p* linkage anchoring both side chains to the poly-GalA*p* backbone of RG-II had been retained. These data also reveal that BT1012 is the sole apiosidase in *B. theta,* cleaving both _D-_apiosyl linkages in side chains A & B. ΔBT1012 oligo#1 & 2, secreted by both mutant and Wt strains were identified as 2-O-Me-_L-_Fuc*p* and 2-O-Me-_D-_Xyl*p-*α1,3-_L-_Fuc*p* by comparison with secreted products from the Wt strain (**Fig. 1C**). The prominent and mutant specific sugar ΔBT1012oligo#3 (∼5mg or ∼5% yield of RG-II starting material)) was successfully resolved by size exclusion chromatography (SEC) (**Fig. 1D**) and analysed by mass spectrometry (MS) yielding mass peaks consistent with the detection of ionic adducts of the RG-II-derived trisaccharide fragment _L-_Rha*p-*α1,3-_D-_Api*f*-β1,2-_D-_GalA*p* (M+NH_4_^+^=490.18, M+K^+^=511.24 and M+Na^+^=511.11) **(CDRO_B33/A26), (Fig. 1E).** This result confirms the previous observation that BT1001 requires prior action of BT1012^5,56^. The identity of the trisaccharide was further confirmed by 2D NMR (^1^H,^13^C-HSQC) and using linkage specific enzymes BT1001 (α-_L-_rhamnosidase) and BT1012 (β-_D-_apiosidase) which together completely depolymerised the structure, releasing all three constituent monosaccharides; _L-_Rha*p*, _D-_Api*f* and the RG-II backbone sugar _D-_GalA*p* (**Fig. 1F, Supplemental fig. 1B**). Attempts to resolve or separate components of ΔBT1012smear by size exclusion chromatography (SEC) were unsuccessful, however, we gained some insight into their composition by acid hydrolysing and analysing them by HPAEC-PAD. The mixtures contained not only _L-_Rha*p*, _D-_Api*f* and _D-_GalA*p* sugars but also _L-_AceA*f* derived from side chain B in some fractions (also confirmed by NMR), (**Supplemental fig. 1C, D**). These findings identify ΔBT1012 as an effective strain for ready and scalable production of pure **CDRO_B33/A26**. Its ability to retain the Api*f*-β1,2-_D-_GalA*p* linkage in side chains A & B of RG-II, makes it uniquely suited as a platform to further engineer and generate a diverse array of highly specific Gal*p*A-linked CDROs.

#### Tailoring ΔBT1012 to produce specific CDROs linked to terminal _D-_GalAp

To demonstrate that ΔBT1012 can be tailored to produce more complex and specific _D-_GalA*p-*linked CDROs, we used it to generate and purify for the first time, the rare and complex side chain B-derived hexasaccharide **CDRO_B30** (2-O-Me-_L-_Fuc*p-*α1,2-_D-_Gal*p-*β1,3-_L-_AceA*f*-α1,3-_L-_Rha*p-*α1,3-_D-_Api*f*-β1,2-_D-_GalA*p*) and its acetylated variants **(Fig. 1A**, **Fig. 2, Supplemental table 1, Supplemental table 2,)**. Initially, a side chain B related mutant ΔBT0984 (lacking the 2-O-Me-_L-_fucucosidase) was created, to retain the 2-O-Me-_L-_Fuc*p-*α1,2-_D-_Galp glycosidic linkage during the metabolism of RG-II^5^ (**Fig. 1A**). Like ΔBT1012, this mutant accumulated several oligos, most of which were successfully resolved and analysed by MS (**Supplemental fig. 2A-H)**. The most abundant fraction ΔBT0984oligo#1 (∼5 mg or ∼5% yield of RG-II starting material) produced mass peaks consistent with the detection of ionic adducts of the unacetylated side chain B pentasaccharide 2-O-Me-_L-_Fuc*p-*α1,2-_D-_Gal*p-*β1,3-_L-_AceA*f*-α1,3-_L-_Rha*p-*α1,3-_D-_Api*f* (M+Na^+^=796.31, M+K^+^=801.26, and M+HCOO^-^=823.24) **(CDRO_B5) (Supplemental table 1, Fig. 1G, Supplemental fig. 2B).** As with **CDRO_B33/A26,** the structural composition of **CDRO_B5** was confirmed through enzymatic profiling with four RG-II-specific CAZymes (**Fig. 1H)**. Inclusion of BT1012 in the profiling mixture did not result in the production of _D-_Gal*p*A, confirming that **CDRO_B5** lacks the terminal/reducing end _D-_Gal*p*A **(Fig. 1H)**. Demonstrating its quality and scalability, we generated sufficient quantities of CDRO_B5 and used it to validate active site and kinetics of the BT0984 enzyme^57^. Analyses of other prominent ΔBT0984 products ΔBT0984oligo#2 and ΔBT0984oligo#3 showed incremental masses of 42.01 Da and 84.03 Da respectively (compared to ΔBT0984oligo#1), consistent with the detection and purification of single and double acetylated variants of ΔBT0984oligo#1 **(CDRO_B13, CDRO_B18, CDRO_B25) (Fig. 1I, Supplemental table 1**, **Supplemental table 2, Supplemental fig. 2C-E).** This agrees with previous reports of the existence of acetyl modifications on side chain B, on either 2-O-Me-_L-_Fucp, _L-_AceA*f* or on both sugars^7,58^. 2D NMR was also used to confirm anomeric H^1^ shifts for the various sugars and acetyl modifications of **CDRO_B5 (Fig. 1J),** but the precise locations of the acetyl groups are yet to be defined. ΔBT0984oligo#3 yielded mass peaks consistent with the detection of ionic adducts of **CDRO_B4**, further confirmed by 2D NMR data showing H^1^ shift for α-_L-_Ara*p* **(Supplemental fig. 2E).** ΔBT0984oligo#4 and #5 contained complex mixtures of **CDRO_B13/18, CDRO_B21/31** and **CDRO_B5 (supplemental fig. 2F-G).** Strangely, small amounts of other unrelated sugars were also resolved e.g. ΔBT0984oligo#6 which yielded peak masses at 1116.42 and 1158.43 m/z, suggesting the detection of ionised forms of acetylated **CDRO_B10/B15 (Supplemental fig. 2H)**. ΔBT0984, therefore, did not only enable us to readily generate and purify **CDRO_B5,** but also resolve rare, acetylated variants of **CDRO_B5** and other useful CDROs.

**Fig. 2:**
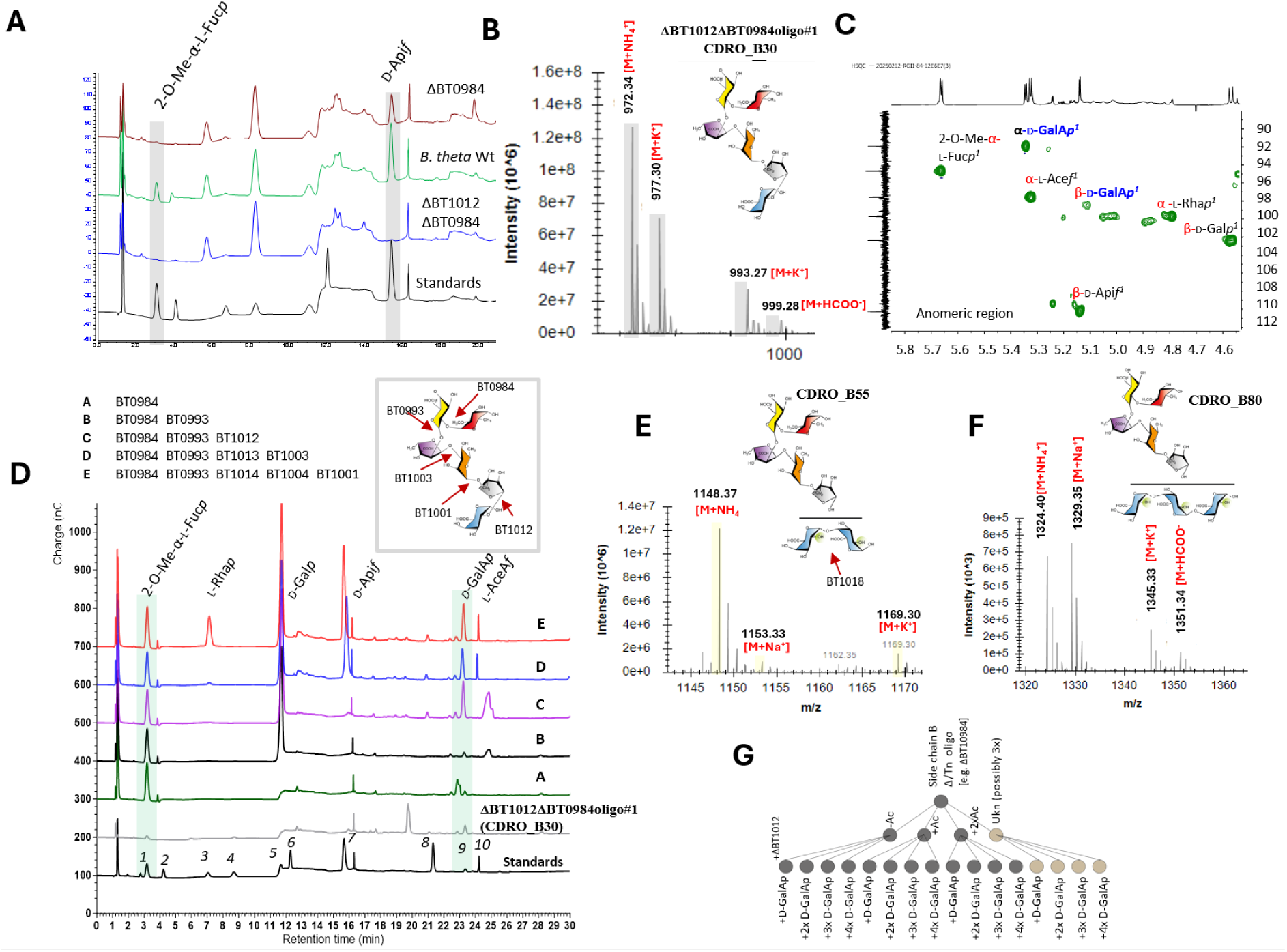
Production, purification and characterisation of side chain B CDROs linked to HG backbone components. **a:** HPAEC-PAD analyses of culture supernatants of ΔBT1012 and ΔBT0984 and a double knockout mutant of both (ΔBT1012ΔBT0984) showing changes in the amount of secreted 2-O-Me-L-Fuc*p* and D-Api*f*. **b:** MS analyses of CDRO_B30 showing various ionic adducts **c:** 2D NMR (^1^H,^13^C-HSQC) of CDRO_B30 **d:** Enzymatic profiling of CDRO_B30 using RG-II specific enzymes confirms detection of HG backbone-derived D-GalA*p*. Standards: 2-O-Me-α-L-Fuc*p* (1), D-Fuc*p* (2), L-Rha*p* (3), L-Arap (4), D-Galp (5), D-Xylp (6), D-Apif (7), D-kdo (8), D-Gal*p* (9), L- AceA*f* (10). **e:** MS analyses of CDRO_B55 showing various ionic adducts **f:** MS analyses of CDRO_B80 showing various ionic adducts **g:** Bayesian-like scheme showing the potential for an unprecedented expansion of side chain B-derived CDROs based on findings in the current study. When a single genetic strain e.g. ΔBT0984 is used for CDRO generation, there is the probability of obtaining unacetylated (-Ac), acetylated forms of the sugar which can be single (+Ac) or bi-acetylated (+2Ac). The existence of more yet-to-be-defined or unknown form (ukn also adds to these possibilities. In a double deletion strain like ΔBT1012ΔBT0984, the number of possibilities increases exponentially with the generation of CDROs with variable amounts of HG backbone components.

We thus reasoned, based on our findings, that combining ΔBT0984 and ΔBT1012 mutations in a single genetic strain (ΔBT0984ΔBT1012) will result in the generation of **CDRO_B30 (Fig. 2A-D),** especially considering the earlier evidence that ΔBT1012 enables retention of terminal/reducing end _D-_Gal*p*A, which is required to complete **CDRO_30** from **CDRO_B5**. As expected, ΔBT0984ΔBT1012 secreted neither 2-O-Me-_L-_Fuc*p* nor _D-_Api*f* (compared to the Wt. strain) **(Fig. 2A)** suggesting that the reducing and non-reducing end glycosidic linkages had been retained during RG-II metabolism, leading to the generation of **CDRO_B30** (∼5mg or ∼5% yield of starting RG-II material)(**Fig. 2A-D**). Several oligosaccharides generated by the strain were resolved **(Fig. 2, Supplemental Fig. 3A-F)**, with the most prominent sugar ΔBT0984ΔBT1012oligo#1 yielding major mass peaks at 972.34 and 993.2 m/z, corresponding to ammonium and potassium adducts of the target sugar **CDRO_B30 (Fig.2B, Supplemental Fig. 3A, B)**. 2D NMR analyses detected H^1^ anomeric signals for all predicted monosaccharides in **CDRO_B30** including both alpha (α) and beta (β) anomeric forms of its terminal _D-_GalA*p* (**Fig. 2C**). A combination of enzymatic profiling and HPAEC-PAD analyses further confirmed the predicted structure of **CDRO_B30** with _D-_GalA*p* detected after treatment of the sugar with the BT1012 (**Fig. 2D**). As with ΔBT0984 oligosaccharides, masses corresponding to acetylated versions of **CDRO_B30** were also detected in other resolved fractions e.g. ΔBT0984ΔBT1012oligo#2 (**Supplemental fig. 3B**). Sugars in ΔBT0984ΔBT1012oligo#3 and #4 fractions were identified by mass as the trisaccharide and disaccharide sugars _L-_Rha*p-*α1,3-_D-_Api*f*-β1,2-_D-_GalA*p* (**CDRO_B33/A26**) and 2-O-Me-_D-_Xyl_-_α1,3-_L-_Fuc*p* (**CDRO_A6**) respectively (**Fig. 1).** These data confirm the first report of the targeted production of the rare sugar **CDRO_B30** and the identification of a corresponding genetic strain ΔBT0984ΔBT1012 for this purpose. Critically, it highlights ΔBT1012 as a versatile platform that can be tailored to generate a diverse array of highly complex, yet very specific _D-_GalA*p-*containing side chain B-derived oligos - the logic being that, the ΔBT1012 mutation can now be combined with any other desired side chain B related mutation (as with ΔBT0984) to target and generate specific _D-_GalA*p-*containing CDROs of interest and possibly their acetylated variants e.g. those in the range **CDRO_B26-CDRO_B50** containing _D-_GalA*p* at their reducing ends (**Supplemental table 1**). Of course, this assumes that mutants lacking all other side chain B related enzymes will also generate specific CDROs, a question which we investigated and successfully confirmed in subsequent sections.

#### Potential for an unprecedented expansion of the CDRO library and new variants

We observed that the content of additional SEC fractions of the ΔBT0984ΔBT1012 strain (e.g. ΔBT0984ΔBT1012oligo#5, #6) yielded abundant ionic mass peaks at m/z 1148.3 and 1329.35 respectively, which were much higher than would be expected for mere ionised forms of **CDRO_B30 (Fig. 2E-F, Supplemental fig. 3E-F).** Interestingly, enzymatic profiling of the oligos using the **CDRO_B30** enzymatic cocktail yielded all constituent monosaccharides of **CDRO_B30** except _D-_GalA*p* (**Supplemental fig. 4A**), suggesting that presence of additional sugars at the base of the chain that are interfering with its release. We thus reasoned that these maybe additional **_D-_**GalA*p* units from an extended HG backbone. To verify this, an exo-polygalacturonase (GH28) enzyme BT1018 known to cleave between _D-_GalA*p-*α1,4-_D-_GalA*p* units of the HG backbone was included in the **CDRO_B30** enzyme cocktail, leading to a strong spike in _D-_GalA*p* as confirmed by HPAEC-PAD **(Supplemental Fig. 4B)**. The masses 1148.37, 1153.33, 1169.30 detected by MS for ΔBT0984ΔBT1012oligo#5 thus corresponded to ammonium, sodium and potassium adducts of the **CDRO_B30**-containing sugar **CDRO_B55 (**2-O-Me-_L-_Fuc*p-*α1,2-_D-_Gal*p-*β1,3-_L-_AceA*f*-α1,3-_L-_Rha*p-*α1,3-_D-_Api*f*-β1,2-_D-_[<u>_D-_GalA*p*</u>]_2_) which has two _D-_GalA*p* units at its reducing end **(Fig. 2E, Supplemental fig. 3E).** On the other hand, ionic masses detected for ΔBT0984ΔBT1012oligo#6 showed an increment of 176.03.1 Da (e.g. 1148.37 m/z to 1324.40 m/z) and 352.06 Da (e.g. 1148.37m/z to 1500.43 m/z) **(Fig. 2F, Supplemental fig. 3F)** consistent with the presence of one and two extra _D-_GalA*p* units. This implies that the octa- and nonasaccharide sugars **CDRO_B80 (**2-O-Me-_L-_Fuc*p-*α1,2-_D-_Gal*p-*β1,3-_L-_AceA*f*-α1,3-_L-_Rha*p-*α1,3-_D-_Api*f*-β1,2<u>[-_D-_GalA*p*]_3_</u>) and **CDRO_B105 (**2-O-Me-_L-_Fuc*p-*α1,2-_D-_Gal*p-*β1,3-_L-_ AceA*f*-α1,3-_L-_Rha*p-*α1,3-_D-_Api*f*-β1,2<u>[-_D-_GalA*p*]_4_</u>) containing three and four reducing end _D-_GalA*p* units respectively had been generated. Interestingly ionic masses of acetylated variants of **CDRO_B105 (CDRO_B113/B118)** were also detected **(Supplemental fig. 3F).** Informed by these findings, we revisited the ΔBT1012 data (**Fig. 1B**), speculating that the undefined ΔBT1012smear fractions likely contained mixtures of CDRO_A26/B33 oligos with variably extended _D-_GalA*p* units in their HG-backbone. This was indeed confirmed by MS with the data showing ionic masses consistent with the detection of **CDRO_B58**, **CDRO_B83** and **CDRO_B108** containing from two to four _D-_Gal*p*A units at their reducing ends (**Supplemental fig. 4C)**. These new findings lay the groundwork for an unprecedented expansion and diversification of the CDRO library with at least seventy-five more side chain B-derived structures or species containing variable lengths of the backbone _D-_GalA*p* units, ranging from **CDRO_B51** to **CDRO_B125 (Supplemental table 1)**. Along similar lines, we also obtained new data suggesting that there are likely more variants (at least 7) of side chain B than previously thought. This is based on HPAEC-PAD data from the analyses of mixtures of purified oligosaccharides of the side chain B related mutant ΔBT0986^5^ which strangely yielded over seven sugar peaks instead of maximum 4 peaks, as expected, based on the number of possible acetyl variants of its main product **CDRO_B3: CDRO_B11, CDRO_B16 and CDRO_B23** (**Supplemental fig. 5A, Supplemental table 1**). Interestingly, all seven products were sensitive to the BT0986 rhamnosidase (**Supplemental fig. 5A)** confirming the presence shared features including the _L-_Rha*p-*α1,2-_L-_Ara*p* linkage targeted by the enzyme. MS analyses also showed peaks of the main expected product **CDRO_B3** alongside those of several yet-to-be-defined products (**Supplemental fig. 5B**). Factoring in our ability to produce a) unacetylated and acetylated side chain B CDROs b) CDROs with variable amounts of reducing end _D-_GalA*p* units c) the existence of yet to be defined variants of side chain B (**Fig. 2G**), it becomes evident that our current approach of using genetic strains has considerable potential to expand the growing CDRO library.

### Screening a *B. theta* genome-wide transposon (Tn) mutant library for new CDRO-generating strains

Here, we aimed to identify new CDRO-generating strains by screening an ordered and barcoded *B. theta* genome-wide Tn mutant library (**Supplemental table 2)**^59^. More importantly, we sought to find out if all other side chain B-related mutants (like ΔBT0984), generate specific CDROs, an outcome that will greatly facilitate efforts to further exploit them to generate new CDRO glycoforms (such as those in the range **CDRO_B26-125** to **CDRO_B/D1-25)**. Of the ∼3277 *B. theta* transposon mutants in the library, 54 contained gene disruptions mapping to RG-II PUL elements, 35 mapping to RG-II-sensitive genes within the PULs^18^, and 16 mapping to genes of previously confirmed RG-II-degrading CAZymes^5,56^ **(Fig. 3A, Supplemental fig. 6)**. Tn mutants for some enzyme including some side-chain B related genes BT0985 (CE20 acetylesterase) and BT0983 (GH2_6 _L-_arabinopyranosidase) were not present. Counter-selectable allelic exchange was used to create KO mutants for some missing Tn strains, although ΔBT0983 failed despite several attempts. Initial screens showed that Tn mutants for various RG-II CAZyme genes in RGII PUL 1 yielded CDROs which were later characterised, while none of those in RGII PUL 2&3 did, highlighting a less critical role of the latter in the enzymatic degradation of RG-II **(Fig. 3B**, **Fig. 4**, **Fig. 5).** Considering the dual activity of BT0996 (side chain A-degrading glucuronidase and side chain B-degrading arabinofuranosidase), its transposon mutant BT0996:Tn was expected to generate both side chain A and B CDROs, however, mainly masses corresponding to side chain A-derived CDROs (**CDRO_A2, CDRO_A_M3**) were detected by MS in partially purified/desalted samples (**Fig. 3B, Supplemental fig.7**), suggesting that the Tn insertion might have only affected the terminal GH2 region of the gene. A complete knockout mutant ΔBT0996 lacking the entire gene was thus created, and its products resolved (fractions ΔBT0996oligo#1-4) and later characterised (**Fig. 8B, supplemental fig. 8A-D**). ΔBT0996oligo#1 yielded a very high molecular weight sugar (m/z=1489.5) of unknown identity while ΔBT0996oligo#2 yielded ionic adducts of acetylated and unacetylated **CDRO_B1** [M+NH4^+^=1352.5] and **CDRO_B9/B14** [M+NH4^+^=1394.92] alongside peaks of several other unidentified sugars (**Supplemental fig. 8B**). Sugars in both fractions were sensitive to BT0996 which released _L-_Ara from the CDROs (**Supplemental fig. 8E**). Strangely the most abundant and best resolved CDROs which eluted in ΔBT0996oligo#3 and ΔBT0996oligo#4 had ionic masses corresponding to complete side chain A CDROs **CDRO_A27/CDRO_A31** [M+NH_4_^+^=1486.45] and **CDRO_A20** [M+NH_4_^+^=1477.4] respectively. The former two are methylated variants of **CDRO_A20.** In agreement with this, treatment of ΔBT0996oligo#4 with BT1010 (α-L-galactosidase) led to the release of _D-_Gal*p* (**Supplemental fig. 8E**). Other side chain B-related mutants BT1019:Tn BT0986:Tn, BT0993:Tn, BT1003:Tn and BT1001:Tn generated largely anticipated and abundant products **CDRO_B2, CDRO_B3, CDRO_B6, CDRO_B7 and CDRO_A7/B8** respectively, with acetylated and unacetylated variants detected in many instances **(Fig. 3D, Supplemental table 1, Supplemental table 2, Supplemental fig. 9-12),** however, masses corresponding to unexpected CDROs were also detected with BT1019:Tn products e.g. **CDRO_B1** (M+NH_4_^+^=1352.5), a CDRO of ΔBT0996 above (**Supplemental fig. 9**). Indeed, we also determined that **CDRO_B1** and **CDRO_B2** resolved better with BT1019:Tn as there were fewer contaminating MS peaks in a SEC fraction of BT1019:Tn (BT1019:Tnoligo#2) (**Supplemental fig. 9**) compared to ΔBT0996oligo#2 (**Supplemental fig. 8B**) which both contain **CDRO_B1**. As with ΔBT0996, mass peaks for several sugars of unknown identity were also detected (**Supplemental fig. 9B-G**). ΔBT0985 also generated a rather interesting picture with the most abundant CDRO purified being **CDRO_B4** [M+Na^+^=933.30], which contains 2-O-Me-L-Fuc*p* and _L-_Ara*p* at its non-reducing end. This was further confirmed by superposition of its 2D NMR spectra with that of a very close structure lacking only _L-_Ara*p* such as **ΔBT0984oligo#5 (Fig. 1H, Supplemental Fig. 11)**. Acetyl groups were also detected in the NMR data suggesting the presence of acetylated **CDRO_B4** species. These data add new information about the hierarchy of side chain B degradation in *B. theta,* being that BT0985’s esterase activity precedes the activities of both the BT0983 α-_L-_arabinopyranosidase and the BT0984 methylfucosidase. BT1001:Tn generated also previously shown to be produced by ΔBT1001. The disaccharide sugar was cleaved by BT1001 rhamnosidase Rha and Api (**Supplemental fig. 12E**). Some side chain A-related mutants also generated unexpected results, for example, BT1010:Tn supernatants contained several species including a dominant CDRO **CDRO_A1** (lacking reducing end _D-_GalA*p*) and the previously reported **CDRO_A20** (<u>66/</u>) (**Supplemental fig. 13A-C)**. **CDRO_A1** and **CDRO_A20** were successfully purified, however other detected CDROs such as **CDRO_M2, CDRO_A8/CDRO_A12** and yet-to-be-identified CDROs co-purified in a complex mixture (**Supplemental Fig. 13D**). BT0997:Tn generated diverse methylated forms of **CDRO_A3**, but the most abundant species were unmethylated (**Supplemental fig. 14**). ΔBT1017 strain yields **CDRO_A-F64** and **CDRO_A-F66** as previously shown^5,56^, however, a closer inspection identified several less abundant mass peaks of previously unreported species terminating in 6-O-Me-_D-_GlcA*p* at the non-reducing ends from **CDRO_M26** to **CDRO_M30 (Supplemental fig. 15)**. A mass peak corresponding to an adduct of **CDRO_A26/CDRO_B33** was detected in BT1011:Tn supernatants (**Supplemental fig. 16A**), however, the identities of several sugars corresponding to major peaks detected in BT1011:Tn and BT1020:Tn supernatants were not obvious or easily predicted by mass analyses (**Supplemental fig. 16**). A ΔBT1020 strain however generated a large and complex CDRO, which was significantly characterised in later sections. Lastly, about twenty (20) other mutants screened including those for genes in RG-II PUL3 (e.g. BT3662 to BT3665, and BT3670 to BT3672), did not secrete or accumulate unique CDROs compared to *B. theta* Wt (**Fig. 3A**, **Supplemental table 2**). In summary, we have generated and identified several new *B. theta* CDRO-generating strains as well as determined the genetic impact of diverse RG-II-inducible genes/metabolic enzymes for the first time, an aspect that remained understudied, despite extensive genetic investigations of other carbohydrate metabolic pathways in *B. theta*^60^. Together with previous work^5^, we have now identified strains producing all major detached side chain A and B CDROs **(CDRO_B1-B8)** (**Fig. 3B**) including in some cases acetylated/methylated variants (**Fig. 3B, Supplemental fig. 3-16, Supplemental table 2**), thus paving the way for an unprecedented expansion of the growing CDRO library using the ΔBT1012 platform.

**Fig. 3:**
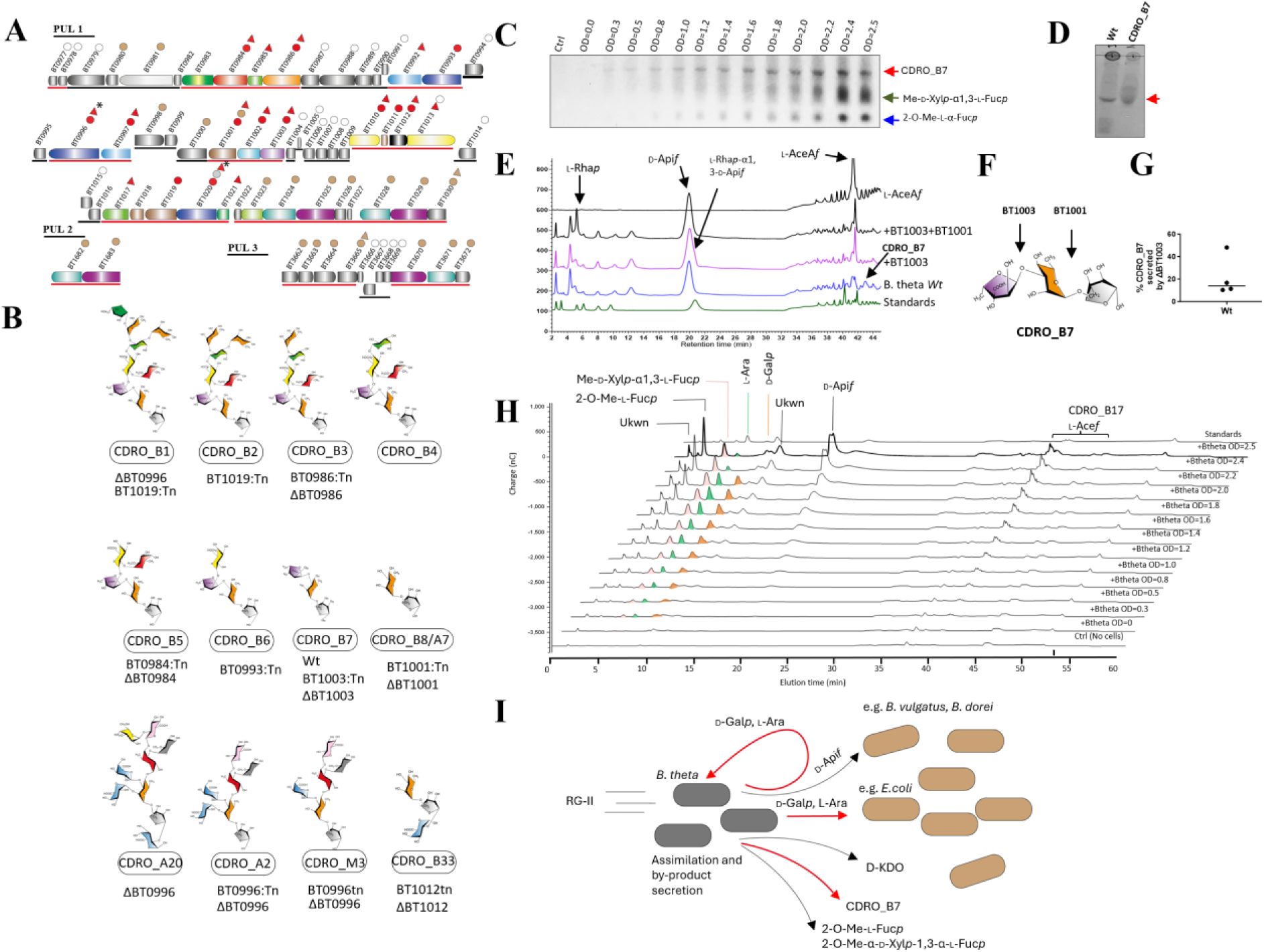
Screening a *B. theta* Tn library for CDRO-generating strains and new insights into RG-II metabolism. **a:** RG-II PUL structure and summary of all genes for which transposon (circles) or knockout/deletion mutants (triangles) were obtained or generated. Genetic loci induced by RG-II are underlined red, while those not induced are underlined black according to previous study^18^. Red and gray filled circles and triangles highlight genes whose mutant strains generated unique CDROs or secretion product profiles compared to a *B. theta* Wt strain. Brown filled circles and triangles highlight genes whose mutant strains did not generate unique CDROS compared to *B. theta* Wt. Gray filled circles are for mutants that generated CDROs that are yet-to-be characterised. Empty circles highlight genes whose mutant strains were not tested largely because they appeared in loci not induced by RG-II or have not been biochemically shown to be implicated in RG-II metabolism. **b:** Examples of prominent CDROs generated and purified from a range of Tn and Δ mutants characterised in this study. All major side chain B sugars were generated and corresponding strains for the production identified **c:** Time course and TLC analyses of *B. theta* Wt secretion products during growth on RG-II at different growth OD’s, showing newly identified product CDRO_B7 (L-AceA*f*-α1,3-L-Rha*p*-α1,3-D-Api*f*)(red arrow) and previously identified products 2-O-Me-D-Xyl*p*-α1,3-L-Fuc*p* and 2-O-Me-L-Fuc*p* (green and blue arrows respectively)^5^ **d:** Comparison of *B. theta* Wt supernatant and CDRO_B7 trisaccharide purified from ΔBT1003 mutant **e:** Treatment of secreted products of *B. theta* Wt supernatant with CDRO_B7-specific enzymes shows release of constituent sugars L-Rha*p,* L-AceA*f and* D-Api*f* **f:** Cleavage sites of CDRO_B7-specific enzymes on the structure **g:** Comparison of secreted CDRO_B7 quantities between *B. theta* Wt and the ΔBT1003. **h:** Time course and HPAEC-PAD analyses of *B. theta* Wt secretion products during growth on RG-II over different OD’s showing newly identified products D-Gal*p* and L-Ara alongside previously identified products Me-D-Xyl*p*-α1,3-L-Fuc*p* and 2-O-Me-L-Fuc*p*^5^. **i:** Summary of diverse secreted RG-II-derived products of *B. theta* Wt and potential cross-feeding pathways based on known metabolic capacities of interacting microbes in previous reports^5,63,64^. Newly identified secreted products and pathways are shown with red arrows.

**Fig. 4:**
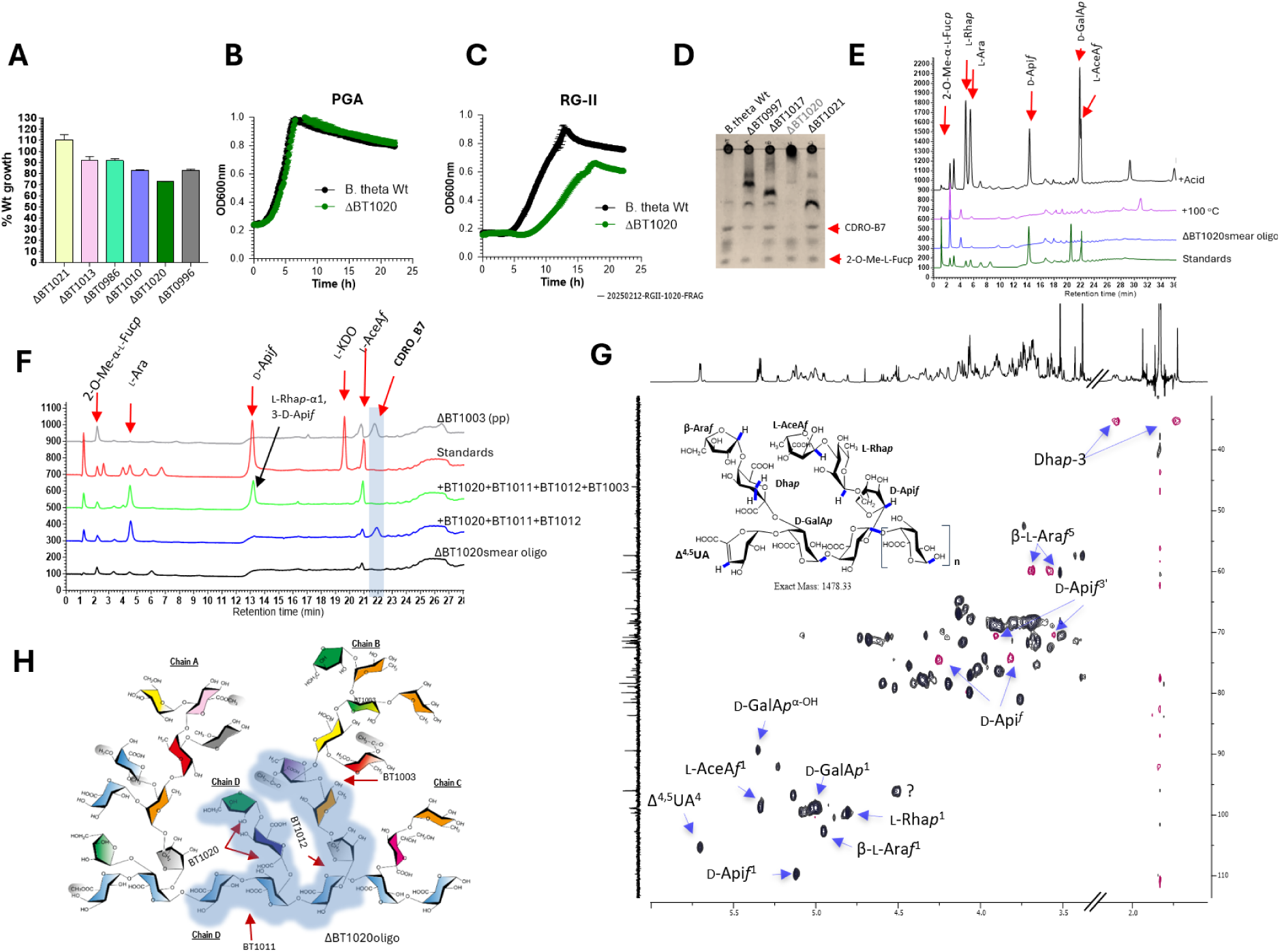
Impact of various side chains on RG-II metabolism in *B. theta* and characterisation of a large and complex CDRO generated by ΔBT1020 strain. **a:** Comparison of growth defects for various side chain related mutants relative to *B. theta* Wt. **b, c:** Growth of ΔBT1020 on polygalacturonic acid (PGA) and RG-II respectively**, d:** TLC analyses showing absence of CDRO_B7 in ΔBT1020 secreted products during RG-II metabolism compared to other mutants and *B. theta* Wt **e:** HPAEC-PAD analyses of ΔBT1020smear oligo after heat and acid treatment **f:** HPAEC-PAD analyses following enzymatic degradation of ΔBT1020smear oligo. Evidence suggests ΔBT1020smear oligo contains side chain D and components of side chain B including CDRO_B7 **g:** NMR analyses of ΔBT1020smear oligo **h:** Structural context of ΔBT1020smear oligo (highlighted region) within whole RG-II.

**Fig. 5:**
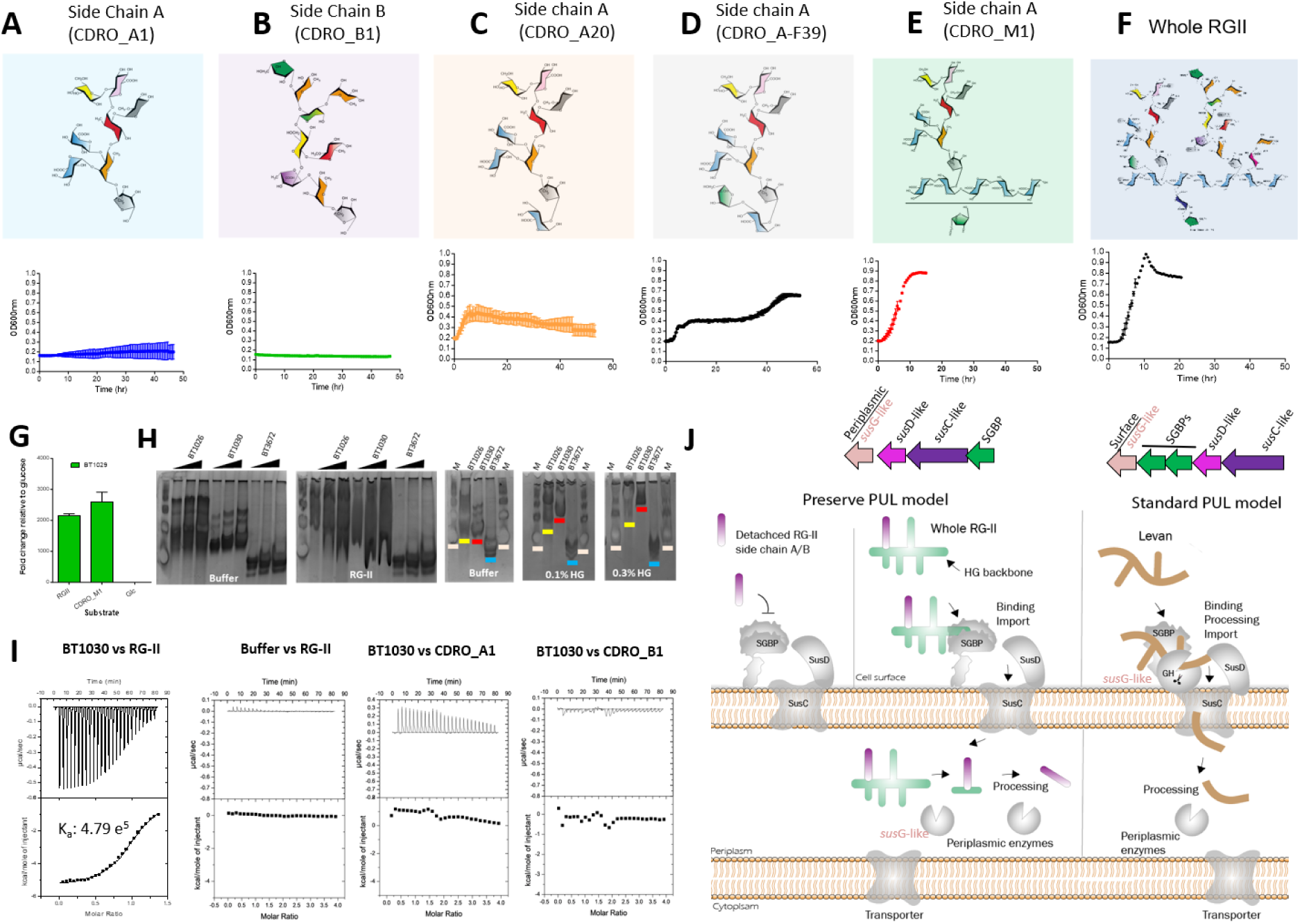
Understanding molecular basis of extracellular RG-II acquisition and import in the human gut microbe *B. theta.* **a-f:** Growth of *B. theta* on diverse side chain A and B-derived CDROs containing or lacking HG backbone components (more growth curves in **Supplemental fig. 21A**). Cells were cultured in 1 %w/v of each substrate. Panels above growth curves show the structure of the most abundant variants present in the purified side chain fractions that were used. Other variants in the fractions included methylated and acetylated forms of the sugars^8^ (Fig. 1i) and side chain A structures lacking the α1-2-linked D-GalA*p*^56^ for CDRO_A20 and CDRO_A-F39 **g:** Induction of an RG-II transporter *sus*CD-like transporter proteins by **CDRO_M1** (more in **Supplemental fig. 21**). **h:** Native affinity gel electrophoresis (NAGE) for the identification of RG-II-binding proteins within various RG-II inducible PULs in *B. theta* (Fig. 3A). Substrates tested included RG-II and HG (homogalacturonan). Control substrate was water and control protein was bovine serum albumin (BSA). **i:** Representative isothermal calorimetry titrations (ITC) for various RG-II binding or non-binding proteins titrated against RG-II. The top panel in each graph shows the raw ITC heats while the bottom panel represents integrated peak areas fitted using MicroCal Origin software. **j:** Comparison of the standard and ‘‘preserve PUL model’’. In the standard model, complex glycans (such as starch^28,53^ glycosaminoglycans^80^ or fructans like levan^81,82^) on the outside of the cell are processed into small oligosaccharides by a surface endo-acting *sus*G-like enzyme^53^ which can be imported through the *sus*-like apparatus into the cell periplasm. In the ‘‘preserve model’’, minimal or no surface degradation of the substrate occurs on the surface of the cell, and the entire glycan is imported. This is also supported by the fact that the main endo-acting (*sus*G-like) enzyme is periplasmic and co-existence of side chain and backbone components in an intact structure is essential for efficient RG-II capture, import and utilisation.

### Exploiting CDROs and genetic tools to better understand RG-II metabolism in *B. theta*

#### New secreted products of RG-II metabolism

CDROs can serve as crucial standards for RG-II metabolic studies. Analysing growth media from time course experiments with Wt *B. theta*, we detected previously unreported secretion products of RG-II metabolism including a sugar of unknown identity (CDROx) which exhibited a similar accumulation pattern to the previously identified secreted sugars 2-O-Me-α-_D-_Xyl*p-*1,3-α-_L-_Fuc*p* and 2-O-Me-_L-_Fuc*p*^5^ **(Fig. 3C)**. Interestingly, CDROx had a similar retention time to the trisaccharide sugar **CDRO_B7** (_L-_AceA*f*-α1,3-_L-_Rha*p-*α1,3-_D-_Api*f* - product of ΔBT1003) by TLC (**Fig. 3D**), an unusual observation given that *Wt B. theta* possesses all enzymes for the complete degradation of **CDRO_B7**. To verify this, spent supernatants from *B. theta* Wt cultures were treated with **CDRO_B7**-specific enzymes leading to the complete degradation of CDROx and the appearance of its constituent monosaccharides _L-_Rhap and _L-_AceA*f* and _D-_Api*f* by TLC and HPAEC-PAD **(Fig. 3E, F, Supplemental fig. 17). CDROx** was thus identified as **CDRO_B7.** Comparison of the total amount of secreted **CDRO_B7** between *B. theta Wt* and ΔBT1003 revealed that between 10% - 50% of total **CDRO_B7** generated in *B. theta Wt* is secreted during normal growth on RG-II **(Fig. 3G, Supplemental fig. 17A-C)** implying that a significant amount is normally assimilated. Two more new secretion products identified as _L-_Ara and _D-_Gal*p* were also confirmed by comparison with monosaccharide standards (**Fig. 3H**). Interestingly, unlike **CDRO_B7** and 2-O-Me-_L-_Fuc*p,* which increased steadily during growth, these sugars peaked in the medium at about OD_600nm_ 1.8-2.2 and then decreased drastically thereafter in the stationary phase, probably due to reabsorption by cells as nutrients become limited under these circumstances **(Fig. 3H)**. Considering that *B. theta* Wt also has known pathways for the utilisation of **_L-_**Ara and **_D-_**Gal*p*^61,62^ (**Supplemental fig. 18**), there is a strong probability that these sugars, including **CDRO_B7** are detected in culture because their extracellular secretion is occurring at a rate faster than their intracellular utilisation. This is different from the cases of 2-O-Me-_L_-Fuc*p* and 2-O-Me-α-_D-_Xyl*p-*1,3-α-_L-_Fuc*p* for which there are no known pathways for the metabolism in *B. theta Wt.* It is also possible that they are not priority nutrients for the host given that various RG-II monosaccharides for example are metabolised at different rates (**Supplemental fig. 18**) and there will be a plethora of these and other RG-II breaKdown products that will become available to the microbe during RG-II metabolism. It is noteworthy that secreted products of *B. theta* like _L_-Ara, _D_-Gal*p*, _D_-Api*f* etc have been shown to be metabolised by other gut microbes^5,63^ including enteric human pathogens like enterohaemorrhagic *Escherichia coli* (EHEC) known to metabolise _D_-Gal*p* and _L_-Ara^63,64^. Indeed it is reported that _L_-Ara can trigger the expression of the EHEC type 3 secretion system which is important for infection^63^. Pathways for other secreted *B. theta* products 2-O-Me-α-_D-_Xyl*p-*1,3-α-_L-_Fuc*p*, _D_-Kdo and 2-O-Me-_L_-Fuc*p* in other gut microbes have not been confirmed. Whether **CDRO_B7** can be taken up other microbes is also yet to be determined.

#### Understanding steric constraints imposed by RG-II side chains and their energetic contribution

RG-II side chains significantly contribute to its resistance to microbial degradation but how individual chains contribute to this process is not well-explored. The availability of various knockout *B. theta* mutants did not only enable us to probe this but also gain an understanding of the structural and molecular basis. Considering the size and complexity of side chains A and B, it is logical to assume that these two chains will pose the highest steric constraints on RG-II degradative enzymes. However, comparing the maximum growth levels for deletion strains lacking enzymes that initiate the breaKdown of various sidechains in RG-II including ΔBT1010, ΔBT1021, ΔBT1013, ΔBT1020, ΔBT0986 and ΔBT0996 (**Supplemental fig. 6**), we made the rather unexpected observation that ΔBT1020, which lacks the enzyme that removes one of the smaller side chains (side-chain D) had the strongest growth defect based on maximum attainable growth ODs (represented as a % Wt growth) **(Fig. 4A).** The nature and impact of the defect was also evident from time course OD_600nm_ measurements of ΔBT1020 vs *B. theta* Wt growths **(Fig. 4B, C).** To better understand these effects, ΔBT1020 secretion products were investigated. On TLC plates, the major product appears as a smear (ΔBT1020smear oligo) close to the origin (**Fig. 4D**). Notably, **CDRO_B7** was not detected in its product profile although clearly visible in the *B. theta* Wt. This suggests that side chain D likely sterically impacts the enzymatic degradation of side chain B which carries **CDRO_B7** and that its prior removal is required for complete degradation of side chain B. It also implies that ΔBT1020 secretion products are more complex than the previously proposed sugar _L-_Araf-sugar β1,5-_D-_Dha-β1,3-[_D-_GalA*p*]n^5^. To confirm this, ΔBT1020smear oligo was purified, heat treated, acid hydrolysed and analysed by HPAEC-PAD. Heat treatment (100°C, <15 min) of the sugar alone led to the release of side chain D, evident by its degradation to _L**-**_Ara after treatment with BT1020 enzyme **(Supplemental fig. 19)**. Acid hydrolysis led the release of several RG-II monosaccharides including _L-_Fuc*p*, _L-_Ara, _L-_Rha*p*, _D-_GalA*p*, _L-_AceA*f*, _D-_Api*f* (**Fig. 4E**) consistent with the presence of side chain D and B components in the analysed sugar. After a series of enzymatic screens with diverse RG-II degrading enzymes, it was also determined that ΔBT1020smear oligo is also susceptible to the unsaturated glycoside hydrolase BT1011 and apiosidase enzyme BT1012 after initial treatment with BT1020 (**Fig. 4F**). Indeed, a combination of these three enzymes led to the release of **CDRO_B7** from ΔBT1020smear oligo (**Fig. 4F**). Production of CDRO**_B7** was further confirmed by treatment with BT1003 which depleted the CDRO_B7 peak and released _L-_Ace*f* and _L-_Rha*p-*α1,3-_D-_Api*f* sugars (**Fig. 4F**). The requirement for BT1011 not only confirmed the presence of RG-II backbone elements but also unsaturated _D-_GalA*p* sugars (ΔGalA*p*). In line with this NMR analyses confirmed the detection of a chemical shift at ∼5.7 ppm, consistent with the detection of the H^4^ atom of an unsaturated uronate (UA^4,5^) sugar (**Fig. 4G**), a typical shift reported following pectolyase activity^65^. Anomeric H^1^, OH and other chemical shifts for constituent monosaccharides in **CDRO_B7** and side chain D (β-_L-_Ara*f* and β-_D-_Dha) were also detected further confirming the presence of both chains in ΔBT1020smear oligo (**Fig. 4G**). ΔBT1020smear oligo is therefore a much larger and complex sugar than previously thought^5^, extending from terminal β-_L-_Ara*f* in side chain D through the HG backbone and terminating in _L-_Ace*f* on side chain B (**CDRO-D/B7**:_L-_AceA*f*-α1,3-_L-_Rha*p-*α1,3-_D-_Api*f*-β1,2-(_L-_Ara*f*-β1,5-_D-_Dha-β1,3-)-[UA-_D-_GalA*p-*α1,4-[_D-_GalA*p-*α1,4-_D-_GalA*p*]n) (**Fig. 4G, H**). This also explains the unexpectedly strong growth defect observed with ΔBT1020 mutant compared to other side chain related mutants. As with ΔBT1012, this revelation sets a stage for the exploitation of ΔBT1020 as a platform for the further expansion and diversification of the CDRO library i.e. by combining ΔBT1020 with various side chain B-related mutations, it will now be possible to generate much larger and complex CDROs containing components of both side chain D and B such those in the range **CDRO_D/B1-25** (**Supplemental table 1**). ΔBT0996 and ΔBT1010 show noticeable defects, although relatively low considering the size of the affected chains. This implies that both chains are easily released from the backbone by lyase and apiosidase enzymes, minimising their steric effects on enzymes required to degrade the rest of the structure. In conclusion, *B. theta* genetic tools and CDROs have not only helped us to better understand the steric impact of various side chains on RG-II metabolism but also provided new information that will enable us to further expand the growing CDRO library.

#### Alternative operational paradigm for PUL_-_mediated glycan uptake in Bacteroidota

Although intracellular enzymatic pathways for RG-II degradation in *B. theta* are well characterized, the mechanisms underlying its extracellular capture and transport across the cell envelop remain unclear. In the standard PUL paradigm, a complex glycan substrate is initially captured on the outside of the cell through binding interactions with a PUL_-_encoded proteins including surface glycan binding proteins (SGPBs) and endo-acting enzymes that bind and partially process the substrate into small oligosaccharide structures respectively for import into the cell periplasm^27,28,66–68^. Genes for the endo-acting RG-II pectolyase BT1023 (PL1_2) and the putative lyase BT0980 are present in the RG-II PUL, however, BT1023 is periplasmic^5^ while the latter is not induced during growth on RG-II^18^ (**Fig. 3A**). Among RG-II PUL-encoded enzymes displaying a strong lipidation signal^69^ (and hence a possible outer membrane/extracellular localisation), is the acetylesterase BT0985 (CE20) which we investigated, comparing it with a previously confirmed extracellularly localised protein BT1030^5^ (**Supplemental fig. 20A, B**). Cellular localisation assays, using a combination of ultracentrifugation and western blotting showed enrichment of the protein in soluble cellular fractions as opposed to BT1030 detected in membrane fractions, suggesting BT0985 is either periplasmic or cytoplasmic (S**upplemental fig. 20A-C**). Aerobic cell-surface activity assays were also performed to probe endolytic cell-surface degradation of RG-II by incubating RG-II with whole cells of *B. theta Wt,* followed by analysis of the substrate using acrylamide gel electrophoresis. However, no detectable endolytic degradation of the substrate was observed (**Supplemental fig. 20D**). With no detectable surface enzymatic activity against RG-II that will conform to the standard PUL paradigm and the accumulation of large complex glycans like the ΔBT1020smear oligo in some mutant strains, this raises the prospect of an alternative PUL paradigm, whereby the entire substrate is captured and imported intact or as a whole.

To further investigate this, a series of growth experiments with *B. theta* Wt against a variety of complex and simple CDROs (generated using our genetic strains or acid hydrolysis^70^) were performed. The range of CDROs tested primarily consisted of side chain A/B–derived variants featuring different chain lengths of the HG backbone or none e.g. **CDRO_A1, CDRO_B1, CDRO_A20, CDRO_A-F39, CDRO_AF66, CDRO_B7, CDRO_M1** as well as whole RG-II. (**Fig. 5A-F**, **Supplemental fig. 21, Supplemental table 1**). *B. theta* generally failed to grow to detectable levels (over 48 h period) when cultured with CDROs lacking the HG backbone (**Fig. 5A, B**). However, growth was gradually observed with increasing complexity of the glycan and the presence of the HG backbone (**Fig. 5A-F**). To eliminate possibility that the significant growth observed with **CDRO_M1** could have solely been due to the presence of the backbone HG or poly-_D-_GalA*p* units (and hence independent of the RG-II machinery), qPCR analyses were performed on cells cultured with **CDRO_M1** showing induction of various RG-II PUL transporter or *sus*C proteins BT1025, BT1029, BT1683, BT3670 (**Fig. 5G**, **Supplemental fig. 21**). Taken together, the results suggest that an intact HG backbone is important for RG-II utilisation and that side chain units on their own are less suitable for microbial uptake. Given that the HG backbone does not induce the RG-II utilisation machinery^18^, we hypothesised that its main role is likely to enable the import of whole RG-II into the cell. To test this, native affinity gel electrophoresis (NAGE) and isothermal titration calorimetry (ITC) experiments were performed with three genes of unknown function but also putative RG-II SGBPs BT1030, BT1026, and BT3672 due to their proximity to the canonical *sus*CD_-_like pairs (**Fig. 3A**). Notably, BT1026 is located upstream of the *sus*C gene, unlike all *sus*E-positioned SGBPs characterised to date. BT1030 and BT1026 (but not BT3672) showed retarded migration on NAGE gels supplemented with RG-II and HG and more so with increasing concentration of HG from 0.1 to 0.3% (**Fig. 5H**). The retardation effect was more profound with BT1030 compared to BT1026, with ITC data showing that it binds with at least a two-fold higher affinity (association constant (K_a_)=4.8x10^5^) than BT1026 (K_a_=1.9 x10^5^) (**Fig. 5I, Supplemental fig. 22**). Binding to purified side chain A/B CDROs was also assessed by ITC, but no saturation or significant binding was observed (**Fig. 5I**). BT1030 and BT1026 successfully crystallised for X-ray structural studies, however, electron density maps in several regions of the structure were not sufficient for confident assignments. Consistent with their binding preferences, AlphaFold structural data showed that both BT1030 and BT1026 adopt β-helix/solenoid fold (similar to that of the novel CBM89 domain in CapCBM89-GH10 xylanase^71^) and strongly align with each other (∼73.2%, rmsd 1.11Ǻ) but not with BT3672 which shows a completely different fold (**Supplemental fig. 23A, B**). The β-sheet fold of BT1030 also shows three-dimensional homology to a predicted HG-binding protein BT4112 from a PUL in *B. theta* that metabolises HG^18,54^ (∼72%, rmsd 2.9Ǻ) (**Supplemental fig. 24**). Other structurally similar homologues of BT1030 were searched on PDBefold, with the top hit being a capsule-specific depolymerase (PDB:6tku) from Klebsiella phage^72^. There was no conservation of active site residues, consistent with BT1030 being a specialised binding protein (**Supplemental fig. 24B**). Strangely, the *sus*D_-_like protein BT1028 showed no binding to either HG or RG-II *in-vitro* by NAGE (**Supplemental fig. 25A, B**). Lastly knockout mutants of BT1030 (ΔBT1030) and its functional *sus*CD_-_like pair BT1028-29 ((ΔBT1028-29) of the RG-II PUL showed defective growth on RG-II, highlighting the importance of their transport functions during RG-II metabolism (**Supplemental fig. 25C**). Taken together, these results reveal new insights into the mechanisms governing extracellular RG-II capture and import in *B. theta*. Detached RG-II side chains on their own are weakly metabolised due to poor interaction and uptake by surface SGBPs and the rest of the RG-II surface apparatus. The side chains however must be imported since the HG backbone on its own is incapable of inducing the RG-II PUL. Efficient extracellular RG-II capture and uptake in *B. theta* is therefore dependent on the co-existence of both the backbone and side chain components in an intact or unprocessed structure. This allows SGBPs to bind the HG backbone, and in the process pull the rest of the structure towards the remainder of the *sus*CD transport apparatus for import. Therefore, unlike the prototypical PUL model or paradigm (**Fig. 5J**), which works by processing and simplifying glycans at the cell surface into smaller oligosaccharides for subsequent uptake, the RG-II PUL has rather evolved to preserve the structural integrity and complexity of the substrate during uptake, highlighting an alternative operational or mechanistic paradigm for PULs which we have termed ‘’the preserve paradigm or model’’ (**Fig. 5J**). In this model, different components of the glycan, each playing independent, yet integral roles, are preserved extracellularly to facilitate the efficient capture, import and utilisation of the whole glycan. The periplasmic localisation of the critical RG-II endo-lyase BT1023 and the BT1012 apiosidase capable of releasing RG-II side chains support the proposed model, as this limits extracellular endolytic degradation of RG-II. Our data are also in agreement with a previous study tracking the uptake of whole RG-II with fluorescent glycan conjugates^73^, where the authors show that BTΔRGII mutant lacking the entire RG-II PUL1 but not the orphan *sus*CD-like pair BT1682/1683 of RG-II PUL2 (**Fig. 3A**) still exhibited intracellular RG-II staining, suggesting these two non-enzymatic proteins alone are capable of importing whole RG-II.

### New CDRO-generating systems and alternative RG-II metabolic pathways

Thus far, evidence suggests that *B. theta* is a highly efficient and tailorable system for the production of diverse CDROs. *B. theta* is also an anaerobe and encodes the only microbial RG-II metabolic system that has been functionally characterised to date^5^. Identifying new or alternative pathways in other microbes and environments will not only deepen our understanding of RG-II-microbe interactions beyond the human gut but also open up new engineering opportunities for CDRO generation. In this light we performed an extensive bioinformatic and biochemical investigation of potential RG-II metabolic pathways in other microbes including plant and environmental microbes. Analysing thousands of genomes (22053 genomes) from the CAZy database^74^ (**Figure 6A, Supplemental table 3,**) we observed that ∼130 species from Bacteroidia, Chitinophaga, Cytophagia and Sphingobacteria (all Bacteroidota) have a complete RG-II degradome based on the presence of RG-II-degrading gene families, and another ∼130 lack one or two (sub) families, including one non-Bacteroidota species *Verrucomicrobia sp*. More diversity was observed in ∼170 species containing at least half of the degradome in bacteria and Eukaryota phyla including Ascomycete and

**Fig. 6:**
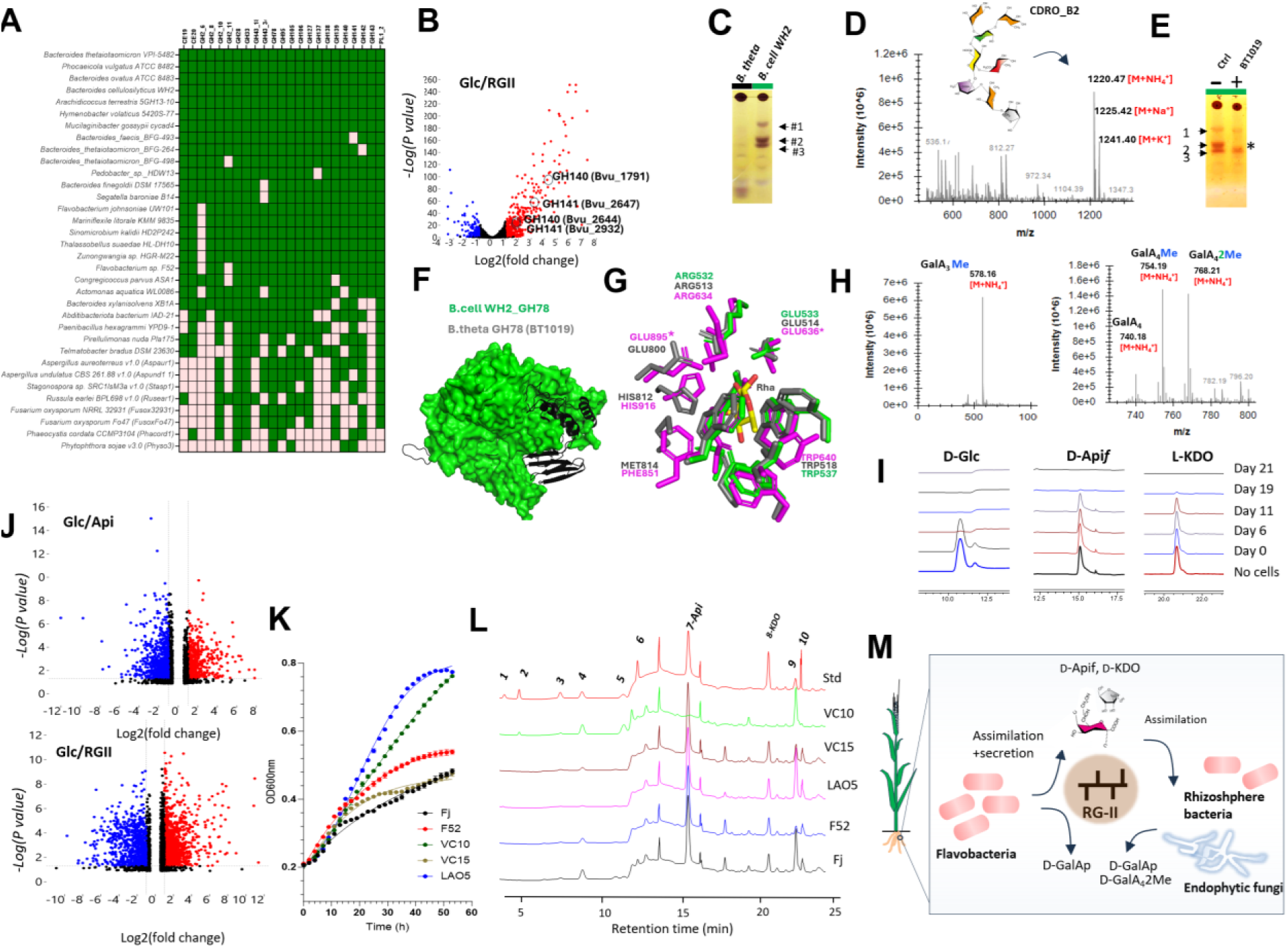
Alternative microbial RG-II metabolic pathways in gut and environmental microbes and new insights. **a:** Heatmap from screening of thousands of microbial genomes from diverse environments from the CAZy database and identification of RG-II-degrading CAZymes (full data in **Supplemental table 3**) **b:** Volcano plot showing differential expression of *P. vulgatus* genes following growth on RG-II as sole carbon source. Red and blue dots are for genes that are differentially expressed (log2(fold change) ≥1or ≤-1) to statistically significant levels (p>0.05) as opposed to black dots. Redundant gene families GH140 and GH141 are shown in black circles. **c:** Evidence for alternative RG-II degradative strategies in another prominent human gut microbe *B. cell*WH2. TLC comparison of *B. theta* Wt and *B. cell*WH2-secreted products shows the generation of three unique CDROs in *B. cell*WH2 **d:** MS analyses of *B. cell*WH2 secreted products in culture media shows several mass peaks including prominent peaks consistent with the detection of the side chain B sugar **CDRO_B2 e:** Treatment of *B. cell*WH2 secreted products with BT1019 α-L-rhamnosidase enzyme and TLC analyses identifies the band corresponding to **CDRO_B2 f: 3D**-structural comparison of *B. cell* WH2_GH78 and *B. theta* GH78 (BT1019) proteins showing missing protein folds in *B. cell* WH2_GH78 (shown in black/gray) **g: C**omparison of the active sites of structures in **f** with that of a well-characterised GH78 CAZyme (SaRha78A) from *S. avermitis*^76^) showing the absence of key catalytic residues (starred) in *B. cell* WH2_GH78 **h:** MS analyses of products generated by root associated *Fusarium spp*. shows detection of methylated oligogalacturonides following growth and degradation of RG-II (**Supplemental fig. 32b, c**)) **i:** Time course and HPAEC-PAD analyses of RG-II core sugars D-Api*f* and D-Kdo alongside D-Glc*p* (Glc) as they are metabolised over time by *F. oxysporum* **j:** Volcano plot of proteomic data after growth of *F. oxysporum* on RG-II and D-Api*f*. Red and blue dots are for genes that are differentially abundant (log2(fold change) ≥1or ≤- 1) to statistically significant levels (p>0.05) as opposed to black dots **k:** Growth curves of *Flavobacteria* spp. on RG-II showing variation in RG-II metabolising capabilities **l**: HPAEC-PAD analyses of spent culture supernatants in **(k)** shows detection of a diverse array of sugars including L-Ara, D-GalA*p,* D-Api*f*, D-Kdo etc, the latter two of which are metabolised by *F. oxysporum* in (**i**). **m:** Deduced model of potential RG-II cross-feeding networks in the soil based on findings in this study. Flavobacteria *spp*. are highly efficient degraders of RG-II compared to fungi which take longer to degrade and metabolise RG-II. Secreted products of Flavobacteria *spp*. metabolism of RG-II such as D-Api*f*, D-Kdo can be metabolised by both endophytic and rhizosphere soil bacteria (Fig. 6i**, Supplemental fig. 32f).**

Basidiomycetes fungi. Amongst the strains were commensal endophytic as well as economically important fungal pathogens (**Fig. 6A, Supplemental table 3,**). Several RDE families were widespread in both bacteria and fungi, but there was notable absence of glycoside hydrolase families GH143, GH138, GH137 and carbohydrate esterase families CE19 and CE20 in many fungal and oomycete strains. GH140s were widespread in soil, endophytic bacteria, oomycete species and fungi but rarely in *Fusarium* species (**Fig. 6A, Supplemental table 3**). Drawing from our experience with *B. theta*, these and other select strains lacking a few RG-II CAZymes and possessing glycan secretion capacity could potentially be exploited as CDRO-generating system. High gene family redundancy was also evident in many species and although this could potentially enhance RG-II metabolic efficiency and competition, it may represent a significant obstacle for the purposes of CDRO generation, which is highly reliant on bespoke or non-redundant enzymatic activities. Multiple copies of RG-II CAZymes were detected in close relatives of *B. theta* like *B. cellulosilyticus* WH2*, Phocaeicola vulgatus 8482,*

*B. xylanisolvens* XB1A*, B. ovatus* 8483 (**Supplemental table 3**). In *P. vulgatus 8482*, transcriptomic studies showed that redundant gene copies of GH140, GH141, GH95 etc., alongside tens of CAZymes and other proteins that are differentially expressed during growth on RG-II (**Fig. 6B, Supplemental fig. 26, 27, Supplemental table 4**) suggesting alternative pathways. These studies also allowed us to correct and reassemble PUL predictions for this microbe in the PULDB database^74,75^ by identifying several co-regulated loci (**Supplemental fig. 27C)**. These include a large region spanning BVU_01767-BVU_1816 (50 genes; 17 CAZymes, 4 SusCD), BVU_2910-BVU_2925 (16 genes; 3 CAZymes and 3 SusCD), BVU_0214-BVU_2019 (not predicted as a PUL with no CAZymes or SusCD pairs), BVU_0299-BVU_0300 (SusCD pair), BVU_1983- BVU_1985 (two CAZymes separated by one gene), and the CAZyme cluster BVU_2642-BVU_2648 (**Supplemental fig. 27C)**.

Surprisingly, despite possessing all RG-II-degrading CAZyme families (**Fig. 6A**), the wild_-_type strain of *B. cellulosilyticus* WH2 showed defective growth on RG-II, generating at least three unique oligos BcellWH2oligo#1-3 based on TLC analyses and comparison with *B. theta* Wt secreted products (**Fig. 6C, Supplemental fig. 28A, B**). Mass spectrometry analyses of spent culture supernatants showed detection of a range of masses, the most abundant being mass peaks for the ionic adducts of the side chain B/rhamnose-containing octasaccharide **CDRO_B2** (M+NH_4_^+^ [1220.47], M+Na^+^ [1225.42], and M+K^+^ [1241.40]) (**Fig. 6D**). When the culture was treated with the **CDRO_B2-sepcific** enzyme BT1019 (GH78/α-rhamnosidase), only BcellWH2oligo#2 was degraded, leading to the release of _L-_Rha and confirming its identity as **CDRO_B2** (**Fig. 6E, Supplemental fig. 28B**). In line with this, the amount of secreted _L-_AceA*f*, (unique component of side chain B and **CDRO_B2 (Fig. 1A))** significantly decreased by about ∼6-fold compared to levels in *B. theta* (**Supplemental fig. 28B, C)**. The production of **CDRO_B2** by *B. cell*WH2 and its sensitivity to the *B. theta*-derived GH78 enzyme indicated possible defects in either the native *B. cell*WH2 *GH78* homologue or other parts of the genomic locus carrying the gene. Genomic, sequence and structural comparisons targeting homologues of *B. theta* GH78 in three closely related strains *B. cell*WH2*, B. finegoldii* 17565*, B. ovatus* ATC8483 and *B. cell* DSM 14838 (closest relative of *B. cell*WH2) showed strong synteny or conservation of gene order around GH78 homologues, protein sequence and structure (**Supplemental fig. 29A, B, Supplemental fig. 30A**), except for the *B. cell*WH2 GH78 gene, which surprisingly, has lost significant parts of its gene and protein structure (**Fig. 6F, G**). These include regions spanning several α-helices and β-strands (β41, α14, α16, α17, α18, β42, β44, β45, β46, β47, β48, β49) some carrying critical catalytic acid residues based on comparison with the well characterised structural homologue SaRha78A *from S. avermitis*^76^ and correspond to the introduction of a stop codon at the gene C-terminus (**Supplemental fig. 30A**). Additional tests also showed that none of the compared strains accumulated **CDRO_B2** during growth on RG-II, consistent with the possession of intact and active GH78s (**Supplemental fig. 30B**). **CDRO_B2,** 2-O-Me-_D-_Xyl*p-*α1,3-_L-_Fucp, _D-_Kdo and other yet-to-be-defined oligos were successfully purified by SEC as shown by HPAEC-PAD analysis (**Supplemental fig. 31A, B**), making *B. cellWH2* a natural and readily available system for their scalable production and recovery. *B. cell*WH2 is also genetically tractable^77,78^ and hence represents a valuable alternative to *B. theta* for CDRO production.

The metabolism of RG-II and key component sugars _D-_Api*f* and _D-_Kdo was also evident in some plant/soil associated fungal and Flavobacteria strains. Endophytic fungal isolates from the microbiota of the healthy Barley plant including *Fusarium spp* slowly degraded RG-II after several days (>7 days), evident by an altered migration on acrylamide gels and the release of mono, di-methylated, unmethylated oligogalacturonides and _D-_GalA*p* into the culture medium (**Fig. 6H**, **supplemental fig 32A-C**). An isolate, identified by 16s sequencing as *Fusarium oxysporum* also metabolised _D-_Api*f* and _D-_Kdo (**Fig. 6I**), completely depleting them from culture between 7-19 days. Utilisation of _D-_Api*f* was also detected in rhizosphere microbes from *Nicotiana benthamiana* (**Supplemental fig. 32D-F**), revealing it as an important nutrient source for both endophytic and rhizosphere plant-associated microbes. Comparative proteomic analyses and peptide mapping (against the proteome of a reference strain *F. oxysporum* Fo47) identified several differentially expressed proteins when cultured on RG-II and _D-_Api*f* (**Fig. 6J, Supplemental fig. 33, Supplemental table 5, 6**) including predicted CAZymes associated with RG-II metabolism (e.g. GH28, GH78, GH43, GH105, etc.,), kinases, oxidases, dehydrogenases, transporters as well as uncharacterised or hypothetical proteins. Interestingly, no previously described bacterial _D-_Api*f* metabolic enzymes^17^ (e.g. _D-_apionate oxidoisomerase, 3-oxo-isoapionate-4-phosphate decarboxylase, and 3-oxo-isoapionate transcarboxylase/hydrolase etc.,) were identified, suggesting possible alternative metabolic pathways for rare sugar catabolism in *Fusarium spp* (**Supplemental table 6)**.

Lastly, we show that five aerobic soil Flavobacteria strains including *Flavobacterium* VC-15, *Flavobacterium johnsoniae UW101*, *Flavobacterium* F52, *Flavobacterium LAO5* and *Flavobacterium* VC10 (AS3 29) can metabolise RG-II, the most efficient being the latter two (based on growth rate within 48 h) (**Fig. 6K and L)**. Interestingly, most of the *Flavobacterium* strains did not metabolise _D-_Api*f* and _D-_Kdo (unlike *F. oxysporum)*, evident by the secretion of these and other yet-to-be-defined sugars in the growth medium (**Fig. 6L**). As with *Bacteroides*, this sugar secretion capability establishes them as strong candidates for CDRO generation through genomic engineering. Genomic annotation of CAZymes in *Flavobacterium* VC10 and *Flavobacterium F52*, confirmed the presence of all known RG-II-degrading gene families (**Fig. 6A, Supplemental table 7),** in line with their robust RG-II-metabolising capacities. Comparison of the RG-II PUL structures across other sequenced Flavobacteria showed some key differences with *B. theta*. Notably, all genes are located on the same strand, without interruption, starting an AraC-like regulator (e.g. Fjoh_4106) likely compensating the absence of the hybrid two-component system (HTCS). Other noticed absences include the GH2_6, GH2_11, GH105 and GH141. In most strains, the first is totally absent while the latter could be found only 10 genes away (e.g. Fjoh_4116) adjacent to related CAZymes (GH28 and PL10). GH2_11 and GH105 could be found in a more distant loci (e.g. Fjoh_4221 to Fjoh_4236) with related enzymes (CE20, GH28, GH106) and more surprisingly a GH117, mostly reported as α-L-anhydrogalactosidase so far, opening questions about plant-specific RG-II variations or possible novel activities in GH117. Additionally, the repeated presence in these PULs of CAZy families not previously associated with RG-II metabolism, such as CE8, CE12, GH2_1 and GH29, candidate β-galactosidase and α-_L_-fucosidase (**Supplemental Fig. 34).**

Given novelty and frequency of the CE8 family, it was further investigated, starting with the CE8 gene Fjoh_4096 from *F. johnsoniae UW101* (**Supplemental fig. 35**). We show that, recombinantly expressed Fjoh_4096, targets the RG-II backbone, as it could de-esterify purified di-methyl tetra-galacturonic acid (GalA_4_2Me) and methyl-trigalacturonic acid (GalA_3_Me) (**Supplemental figure 35A-G**). As further proof of its activity and importance, the RG-II backbone-degrading polygalacturonase BT1018^5^ efficiently processed the resulting de-esterified products (**GalA_4_**) to _D-_GalA*p* after pretreatment with Fjoh_4096 (**Supplemental fig. 35C, D, E**). It was also active against methylated apple pectin, promoting its gelation in the presence of Ca^2+^ ions^79^ (**Supplemental fig. 36F**). It did not process di-methylated **CDRO_M26** containing side chain A components, likely due to steric constraints from associated side chain A (**Supplemental fig. 36G**). Considering the abundance of *Flavobacteria* in soil and plant root systems and their efficiency in metabolising whole RG-II (compared to fungi), we speculate that they are likely to be primary or keystone degraders RG-II in these environments. Targeting of _D-_Api*f* and _D-_Kdo (both metabolic by-products of *Flavobacteria)* by endophytic and rhizoplane microbes further solidifies this notion and highlights a new mechanism for bacterial_-_fungal cross-feeding within plant and soil microbiomes.

To conclude, these studies highlight new and alternative metabolic pathways for RG-II and its component sugars in other microbial systems, some from different environments other than the human gut. Deeper biochemical studies of these pathways, especially in aerobic and genetically tractable systems like *Flavobacteria* will not only shed new light into the mechanism underpinning this process and the impact on plant-microbe interactions but offer new and likely cheaper engineering opportunities for the generation of new CDROs and their variants.

## Materials and methods

### B. theta strains and cultures

Wild type *B. theta* VPI-5482 (*B. theta* Wt), Bacteroides and genetic *B. theta strains* were cultured anaerobically in rich and minimal medium as described previously <u>3/</u> <u>66/</u>. Briefly, cells were cultured in a Whitley A25 Workstation (Don Whitley) at 37 °C in rich medium such as Brain Haeart Infusion (BHI) or tryptone yeast extract glucose (TYG) medium or minimal medium (MM) containing apple RG-II (0.5-1% w/v). RG-II was purified as described in Ndeh *et al*^5^. Small volume cultures (∼50 ul to 200 ul) were performed on 96/384 well plates and monitored using a BioTek Epoch® Microplate Spectrophotometer (Agilent technologies) while large volume cultures ∼5 ml were prepared in glass test tubes and monitored using a Biochrom WPA cell density meter (Cambridge, UK). *B. theta* single and double deletion mutants were created by counter selectable allelic exchange as described previously^83^ with the following modifications; a) gene cloning was performed using In-infusion® cloning kit (Takara) into pExchange plasmid b) conjugation of *tdk*− and *E.coli* cells S17-1 λ pir strains were performed at OD_600nm_ of 0.5. *B. theta* genome-wide Tn library was resourced from the authors of Arjes *et al*.^59^

### RG-II purification

RG-II from wine and apple were purified as described previously^5,84^. For *N. benthamiana* RG-II, 5 g of plant leaves were freeze-dried and blended using a Kenwood BL350 blender. Further tissue disruption was performed using a tissue Lyser II (Qiagen). Ground tissues were washed 5 times with 100 mL of ethanol every 10 min for 1 h and the precipitates filtered each time using a silicate filter until it goes brownish white. It was then transferred to a glass beaker and washed with 100 ml 80:20 methanol/chloroform for 30 min and filtered again. The resulting precipitate was washed in 100 ml absolute acetone for 30 min and later dried in a 37 °C oven overnight to obtain alcohol insoluble residue (AIR). An amount of AIR (250 mg) was dissolved in 10 mL of 50 mM NaOAc buffer (pH 5.0) containing 4 mg of pectolyase enzyme *Aspergillus japonicus* and incubated at 37 °C while shaking overnight. Next day, the digested AIR was centrifuged at 4000 x g for ∼10 min and the supernatants recovered and dialyzed in water. The resulting pectic extract was resolved by SEC in 50 mM NaOAc pH 5.0 on a Sephdadex G75 resin and RG-II fractions collected

### Production and purification of CDROs

The protocol was as described in Ndeh *et al.*^5^ with modifications. In summary, *B. theta* genetic strains were cultured anaerobically (Whitley A35 Workstation; Don Whitley, UK) in minimal medium containing 1-2% apple, wine or *N. benthamiana* RG-II overnight for 48 h at 37 °C. CDROs secreted in cell cultures were recovered by centrifugation at 17000 x g for 10 min and filtering through a 1.2 μm filter (VWR).

For large-scale high-resolution purification, 5 mL of culture supernatants containing CDROs were resolved on a Bio-Gel P2 gel extra fine resin (Biorad cat. #1504118) loaded on a Econo-Column® chromatography column (2.5 × 120 cm) equilibrated with 50 mM acetic acid. The flow rate was set at 0.2 ml/min and 2 ml fractions were collected and analysed by TLC.

For small scale purification or desalting, chromatography was performed using a 1.5 × 10 cm Econo-Column (Bio-Rad) packed with 1 mL of Bio-Gel P2 resin. The column was connected via standard tubing to a peristaltic pump (Pharmacia Biotech) for controlled flow. Prior to use, the bed was equilibrated with ultrapure water. Aliquots of 50 µL of CDRO-containing supernatant were applied directly to the gel and the stopcock was briefly opened to allow the sample to enter the matrix by gravity flow. The column was then reconnected to the peristaltic pump (flow rate 1 drop (∼50 µL) /15 seconds) and fractions collected sequentially into a 96-well plate, typically up to row D (∼48 wells). Fractions were monitored by TLC.

The protocol for the production and purification of acid hydrolysed CDROs was as described in Buffetto et al.^70^ Abundant CDROs produced by acid hydrolysis included CDRO_A1, CDRO_B1 and CDRO_M1 which were present in mixtures with their methylated or acetylated variants.

### Production and purification of GalA_4_2Me

One gram of apple pectin (70-75% esterification) (Sigma Cat. 76282) was dissolved in 50 mL NaH_2_PO_4_ buffer pH 7.0 followed by addition of 500 ul of 5 mg/mL pectolyase enzyme (Sigma Cat. P3026). The sample was incubated for 2 d at 37 °C. Digested pectin was freeze-dried and resuspended in 4 ml H_2_0 and 2 mL resolved by size exclusion on a Bio-Gel P-2 extra fine resin (Biorad). Resolved fractions were analysed by TLC, MS and NMR as described above in earlier sections.

#### Mass spectrometry and NMR

CDROs were purified by SEC using a Biorad P2 resin in 0.3w/v % acetic acid. Resolved fractions were collected and analysed by TLC and later freeze-dried and dissolved in LC-grade distilled water. Samples were analysed on an orbitrap fusion mass spectrometer (Thermo Scientific) and Excalibur was used for data acquisition. CDRO samples were diluted 1 in 10 or 1 in 5 in 30% acetonitrile and 5 mM ammonium formate. 10 µl of sample were loaded into the nanoflow probe tips and placed in a nano-sprayflex source (PN ES072). The tip holder was mounted onto the nanoESI capillary holder and positioned to within 30-40 mm of the MS. The MS was operated in positive mode with a spray voltage of 1200-1500kv, CDRO samples are introduced to the MS via static infusion from glass nanoflow capillaries. The ion transfer temperature was set to 310oC. the MS scans were detected in the orbitrap with a resolution of 120000, mass range normal, RF lens 60%, normalized AGC target 100%, maximum injection time 100ms and microscans was set to 1 and quadrupole isolation was on. For 2D-NMR, CDROs (∼10 mg) were dissolved in D_2_O (∼600ul) and spectra were recorded using Bruker Avance NEO 600 MHz NMR spectrometer equipped with TCI CryoProbe. Data were processed using Topsin® or Mnova software.

#### HPAEC-PAD

Samples were analysed using either a DX-500 or ICS6000 HPAEC-PAD system fitted with a Dionex^TM^ CarboPac^TM^ PA1 anion-exchange columns ((ThermoFisher). The run (flow rate 0.4 - 1.2 ml/min) consisted of an initial isocratic phase of separation with 20 mM NaOH followed by a second phase consisting of a gradient (0–100%) of increasing concentration of 500 mM NaOAc in 20 mM NaOH. Resolved and eluted samples were constantly monitored with a fitted electrochemical pulsed amperometric detector (PAD). HPAEC data was analysed using Chromeleon version 7.3.2 (Thermofisher) or GraphPad Prism software version 10 (Prism)

#### TLC

The protocol is as described in Ndeh *et al.*^27^. In summary, for each reaction sample, 2-6 ul (2 µl × n [number of times 2 ul is spotted and dried]) was spotted onto silica gel 60 TLC plates (Merck) and resolved for at least 1 h in TLC running buffer (butanol/acetic acid/water (2:1:1, v/v/v)). Plates were stained or developed with either orcinol sulfuric acid reagent (sulfuric acid/ethanol/water in the ratio 3:70:20 v/v/v and 0.5% orcinol) or diphenylamine–aniline–phosphoric acid (DPA) reagent^85^. Sugar bands were visible after heating of dried plates at ∼70 °C for up to 10 min

### Production of recombinant proteins

Recombinant plasmids for various RG-II degrading enzymes were obtained from Nzytech. Gene expression was achieved in *Escherichia coli* strains TUNER as described in Ndeh *et al*^5,27^. In summary, transformed *E. coli* cells were cultured in LB to exponential phase (∼ OD 0.6-0.8) and induced with 1 mM isopropyl β-d-galactopyranoside (IPTG). Induced cultures were further grown overnight at 16 °C and harvested the next day by centrifugation at 4000 x g for 10 mins. Cell were later resuspended in buffer containing 20 mm Tris/HCl, pH 8.0, plus 100 mm NaCl (Talon) and disrupted by addition of Bugbuster protein extraction reagent (Merck) for 20 mins at room temperature. Released recombinant proteins were purified from the supernantant cell-free extract by immobilized metal affinity chromatography (IMAC) using Talon™ gel resins (Clontech) a described in Ndeh *et al*^67^. For CE8 enzymes, initial cloning was performed using infusion (Takara Bio) protocol and the following primers; CE8F: gtcgcggatccgaattcgagctcgataacaaattgacattaaccgttgctca CE8R: ccgcaagcttgtcgacggagctcttatttggccggattccatccattc

#### Substrate binding studies (NAGE and ITC)

NAGE was performed as described previously^86^. Substrates tested including HG and RG-II were added to a final concentration of 0.5% (w/v) into the 10% w/v separation gels and allowed to solidify. A control gel contained H_2_0 instead of the polysaccharides. Protein samples including a negative control such as bovine serum albumin were applied to wells (0.75mm thick) of the solidified gel and run at room temperature at 150 V for at least 1 h using a Biorad miniprotean system (Biorad). Gels were stained with Coomassie blue. ITC was performed by using a V*P-*ITC calorimeter (MicroCal). Titrations were carried out in 50 mM Na-Hepes buffer, pH 7.5, 200 mM NaCl at 25 °C. The reaction cell contained protein at 50–100 µM, while the syringe contained ligand (between 2–50 mM). The stoichiometry of binding (n), the association constant K_a_, and the binding enthalpy ΔH were evaluated by using MicroCal Origin 7.0 software. The standard Gibbs energy change ΔG and the standard entropy change ΔS were calculated using the thermodynamic equation −RTlnKA = ΔG = ΔH – TΔS where R is the gas constant and T represents the absolute temperature.

#### qPCR

Cells were cultured in RG-II or _D_-Glc*p* followed by RNA purification using the RNeasy Kit (Qiagen). Purified RNA was immediately frozen or converted to cDNA using the QuantiTect Reverse Transcription Kit (Qiagen).Generated cDNA was used alongside respective *sus*C primers for qpcr previously reported^5^ in a LightCycler® 96 real_-_time qPCR machine (Roche). Data was extracted and analysed using GraphPad Prism software version 10 (Prism)

#### Protein cellular localization assay

Recombinant BT0985 and BT1030 proteins were purified and used to generate polyclonal antibodies in rabbit (Eurogentec). *B. theta* cells were cultured overnight in 5 mL minimal media and 1% (w/v) RGII to stationary phase (OD∼ 2.0) and cells harvested by centrifugation at 17 000 x g for 1 min. Pelleted cells were lysed by sonication and various cellular fractions resolved as described previously by ultracentrifugation as described in Kotarski *et al*^87^. Resolved cellular fractions were analysed by SDS-PAGE and western blotting using 1/5000 dilution primary rabbit polyclonal antibodies (Eurogentec) generated against BT0985 and BT1030 and secondary goat anti-rabbit antibodies (Santa Cruz Biotechnology) as described in Ndeh *et al.*^27^. Fractions were analysed by Western blotting using a 1/10,000 dilution of primary rabbit anti-FLAG® antibodies (F7425, Sigma) followed by a 1/5000 dilution of secondary donkey anti-rabbit IgG conjugated to HRP (Santa Cruz biotechnology, USA).

#### Whole cell assays

*B. theta* Wt cells were cultured anaerobically in BHI medium overnight, next day, cells were transferred to minimal medium containing 1% RG-II and cultured overnight. Cells in 300 ul culture volume were harvested by centrifugation at 17000 x g for 1 min, resuspended and washed with 1 mL PBS by centrifuging. Washed cells were incubated at 37 °C with 1% RG-II dissolved in PBS. Controls included cells plus PBS and RG-II plus PBS. After 6h and 12 h of incubation, supernatants were collected and volumes containing 10 ug RG-II analysed by acrylamide gel electrophoresis as described previously^88^

#### Proteomic analyses

*F. oxysporum* cells were cultured in fungal minimal medium (0.2g/L MgSO_4_.7H_2_0, 0.4g/L KH_2_PO_4_, 0.2g/L KCl, 1g/L NH_4_NO_3_, 0.01g/L FeSO_4_, 0.01g/L ZnSO_4_ and 0.01g/L MnCl_2_) containing _D-_Glc, _D-_Apif, or apple RG-II. Cells were allowed to grow for up to 7 days and harvested by centrifugation at 17 000 g in 2 mL Eppendorf tubes. Cells pellets were frozen at -80 until proteomic analyses. *S-Trap Processing of Samples*: Frozen cell pellets were lysed in 150 µl of 5% SDS and 50 mM TEAB, unsoluble material representing cell wall was discarded by centrifugation and protein concentrations were determined using a micro-BCA assay. Aliquots containing 100 µg of protein were processed using the S-Trap mini protocol (Protifi) as recommended by the manufacturer, with minor modifications. Disulfide bonds were reduced with 20 mM DTT for 10 min at 95 °C, followed by alkylation with 40 mM IAA for 30 min in the dark. After loading the samples onto the S-Trap mini spin columns, the trapped proteins were washed five times with 500 µl of S-Trap binding buffer. Proteins were digested with trypsin (Pierce, Thermo Fisher) at a 1:40 enzyme-to-protein ratio in a two-step digestion: first overnight at 37 °C in 160 µl of 50 mM TEAB, and then for an additional 6 h at the same temperature. Peptides were eluted from the S-Trap mini columns by adding 160 µl of 50 mM TEAB, followed by 160 µl of 0.2% aqueous formic acid, and finally 160 µl of 50% acetonitrile containing 0.2% formic acid, with centrifugation at 1,000 × g for 1 min after each addition. Tryptic peptides were dried in a SpeedVac and stored at −20 °C.

*LC-MS:*Peptides (equivalent of 1.2 µg) were injected onto a nanoscale C18 reverse-phase chromatography system (UltiMate 3000 RSLC nano, Thermo Scientific) and electrosprayed into an Orbitrap Exploris 480 Mass Spectrometer (Thermo Fisher). For liquid chromatography the following buffers were used: buffer A (0.1% formic acid in Milli-Q water (v/v)) and buffer B (80% acetonitrile and 0.1% formic acid in Milli-Q water (v/v). Samples were loaded at 10 μL/min onto a trap column (100 μm × 2 cm, PepMap nanoViper C18 column, 5 μm, 100 Å, Thermo Scientific) equilibrated in 0.1% trifluoroacetic acid (TFA). The trap column was washed for 3 min at the same flow rate with 0.1% TFA then switched in-line with a Thermo Scientific, resolving C18 column (75 μm × 50 cm, PepMap RSLC C18 column, 2 μm, 100 Å). Peptides were eluted from the column at a constant flow rate of 300 nl/min with a linear gradient from 3% buffer B to 6% buffer B in 5 min, then from 6% buffer B to 35% buffer B in 115 min, and finally from 35% buffer B to 80% buffer B within 7 min. The column was then washed with 80% buffer B for 4 min. Two blanks were run between each sample to reduce carry-over. The column was kept at a constant temperature of 50oC.

#### Data acquisition and analyses

The data was acquired using an easy spray source operated in positive mode with spray voltage at 2.20 kV, and the ion transfer tube temperature at 250oC. The MS was operated in DIA mode. A scan cycle comprised a full MS scan (m/z range from 350-1650), with RF lens at 40%, AGC target set to custom, normalised AGC target at 300%, maximum injection time mode set to custom, maximum injection time at 20 ms, microscan set to 1 and source fragmentation disabled. MS survey scan was followed by MS/MS DIA scan events using the following parameters: multiplex ions set to false, collision energy mode set to stepped, collision energy type set to normalized, HCD collision energies set to 25.5, 27 and 30%, orbitrap resolution 30000, first mass 200, RF lens 40%, AGC target set to custom, normalized AGC target 3000%, microscan set to 1 and maximum injection time 55 ms. Data for both MS scan and MS/MS DIA scan events were acquired in profile mode. Raw mass spectrometry data was processed using Spectronaut (Biognosys) version 20.3.251119092449 with the DirectDIA option selected. Briefly, these parameters were used for the processing: cleavage rules were set to Trypsin/P, maximum peptide length 52 amino acids, minimum peptide length 7 amino acids, maximum missed cleavages 2. Carbamidomethylation of cysteine was set as a fixed modification, with the following variable modifications selected: oxidation and dioxydation of methionine, deamidation of asparagine and glutamine, acetylation of the protein N-terminus and Gln->pyro-Glu. The FDR threshold for both precursor and protein were set to 1%. Profiling and imputation were disabled. Quant 2.0 was selected. DirectDIA data were searched against uniprotkb_proteome of *F. oxysporum*. Data was was analysed using spectronaught proteomics software.

#### Transcriptomics and qPCR

For RNAseq studies, cells were cultured in RG-II or Glc followed by RNA purification using the RNeasy Kit (Qiagen). Purified RNA was treated with DNAse to eliminate genomic DNA and the final sample (28 ng/ul) later sent for sequencing at the Wellcome Trust Centre for Human Genetics (University of Oxford). For qPCR, RNA was converted to cDNA using the QuantiTect Reverse Transcription Kit (Qiagen).Generated cDNA was used alongside respective SusC primers (b1025F: gtgcctgatcaagtaaatggatat, b1025R: caaaatcatactctaatgaaccgtt, c1683F: tgaagaaagattttacttggagct c1683R: ccgaacaaagcagcagtttcctt, c3670F: gtttcagtggaataggtaatacag, c3670R: gcattgggattttctctcatac, qBT_1029bF: gagtgctaaatatccaaacttaag, qBT_1029bR:ctataagcaactgtcaaattacgt) for qpcr as previously reported <u>66/</u> using a LightCycler® 96 rea_L-_time qPCR machine (Roche). Data was extracted analysed using GraphPad Prism software version 10 (Prism).

### Pectin gelation assay

The protocol is as described in Slavov et al., 2009. In summary, citrus pectin (1%) was dissolved in 5 mL of 3 mM CaCl_2_ in 50 mM sodium phosphate buffer pH 7.0. The mixture (500 uL) was treated 20 uL CE8 enzyme and incubated at 37 °C for ∼15 min in 1 mL cuvettes. Gelation was visualised by adding 500 ul of ½ dilution SDS PAGE sample buffer.

#### Isolation of plant root fungi and taxonomic identification

Barley seeds were surface-sterilised by serial washes in 70% ethanol for 30 s, followed by 5% sodium hypochlorite for 15 min. After sterilisation, seeds were germinated in 0.5 % water agar. Individual seedlings were transplanted into pots filled with agricultural soil (Quarryfield^89^). Plants were grown in a glasshouse at 18/14 °C (day/night) under a minimum light intensity of 150 µmol m⁻² s⁻¹. Each plant received 40 mL of distilled water every two days. Plants were maintained under these conditions for five weeks, until they reached the stem elongation developmental stage.

After five weeks, plants were carefully uprooted. Roots were separated from shoots and washed thoroughly to remove adhering soil. Root material was then surface-sterilised following a previously established protocol^90^. Briefly, ten root segments of 0.5 cm each were excised and placed onto potato dextrose agar (PDA) supplemented with 20 µg/mL chloramphenicol to inhibit bacterial growth. Plates were incubated at room temperature in the dark and inspected daily. Once macroscopically visible fungal growth appeared, a small portion of mycelium was transferred to a fresh PDA plate to obtain a pure fungal isolate.

A sample of fungal mycelium was collected using a sterile inoculation loop and suspended in 10 µL of distilled water. The suspension was boiled for 10 min at 96 °C in a PCRmax Alpha Cycler Thermal Cycler to lyse the cells. The ITS PCR reaction mixture consisted of 4 µL 5× KAPA HiFi Buffer, 1 µL of 10 mg/µL bovine serum albumin (BSA), 0.6 µL of 10 mM KAPA dNTPs, 0.6 µL of ITS-1 forward primer (cttggtcatttagaggaagtaa), 0.6 µL of ITS-1 reverse primer (gctgcgttcttcatcgatgc), 0.25 µL KAPA HiFi polymerase, 8.5 µL distilled water, and 5 µL of the boiled fungal DNA sample. PCR was performed under the following conditions: 94 °C for 5 min; 35 cycles of 98 °C for 30 s, 50 °C for 30 s, and 72 °C for 1 min; followed by a final extension at 72 °C for 10 min. PCR products were visualised via gel electrophoresis, purified using ExoProStar 1-step enzyme according to the manufacturer’s instructions, and Sanger sequenced. Resulting sequences were trimmed and analysed using BLAST searches against both the NCBI and UNITE databases^91,92^ with default parameters. These searches returned the closest matching taxa for each ITS sequence, enabling confident identification of each isolate to the genus level.

## Data availability

Data in this study are available in figures, tables and supplementary information.

## Acknowledgments

We would like to thank acknowledge Arthur Rogowski (Neogen/Megazyme Ireland) and Thierry Docco (INRAE) who contributed to the purification of apple RG-II material. We are thankful to Professors Breeanna Urbanowicz and Malcom O’Neil from the Complex Carbohydrate Research Center (CCRC), Georgia, USA, for providing wine RG-II material. We thank Kerwyn C. Huang and Anthony L. Shiver for access to *B. theta* barcoded Tn libraries and Professors Claire Halpin, Professor Mike Ferguson and Professor Nicola Stanley-Wall at the University of Dundee, UK, for the kind donation of vital reagents and access to analytical lab equipment. We also thank Professor Eric Martens (University of Michigan Medical School, USA) for access to various gut *Bacteroides* strains. This work was supported by a European Union’s Horizon 2021 Marie Skłodowska-Curie grant (agreement No 101068733), a University of Dundee strategic support fund, a Royal Society (RS) University Research Fellowship grant (URF\R1\221864), and a RS research grant (RG\R2\232406) to D.A.N.

## Author Contributions Statement

**D.A.N** produced various *B. theta* knockout genetic strains, **S.A.N and M.B,** performed NMR sample processing and data acquisition, **D.A.N**, **S.A.N, C.R, M.B, and F.P** performed NMR data analyses. **D.A.N**, **R.M.G** and **A. K** carried out the purification and enzymatic profiling or characterization of CDROs by HPAEC-PAD and TLC. **M.C.R** produced and purified acid hydrolyzed CDROs. **D.A.N** and **I.V** performed ITC and NAGE experiments. **D.A.N, R.M.G K.S**, **A.K and G.F.D** expressed and purified various recombinant proteins, **C.R** and **A.A** performed mass spectrometry sample processing and data acquisition. **D.A.N** performed analyses of proteomic and transcriptomic data. **D.A.N** and **F.P** performed analyses of mass spectrometry data, **C.M.E** performed the isolation and confirmation of Barley endophytic fungal strains, **I.L** performed growth cultures of *Flavobacteria* strains. **N. T** and **B. H** performed bioinformatic analyses of RG-II CAZymes, **D.A.N** conceptualized the work and wrote first draft of the manuscript. **All** authors contributed to editing and revision of the manuscript

## Competing Interests Statement

The authors declare no competing interests

## Notes

### Competing Interest Statement

The authors have declared no competing interest.

