## Supplemental figures for "Towards a comprehensive chemical and genetic tool library for rhamnogalacturonan-II oligosaccharides and exploitation"

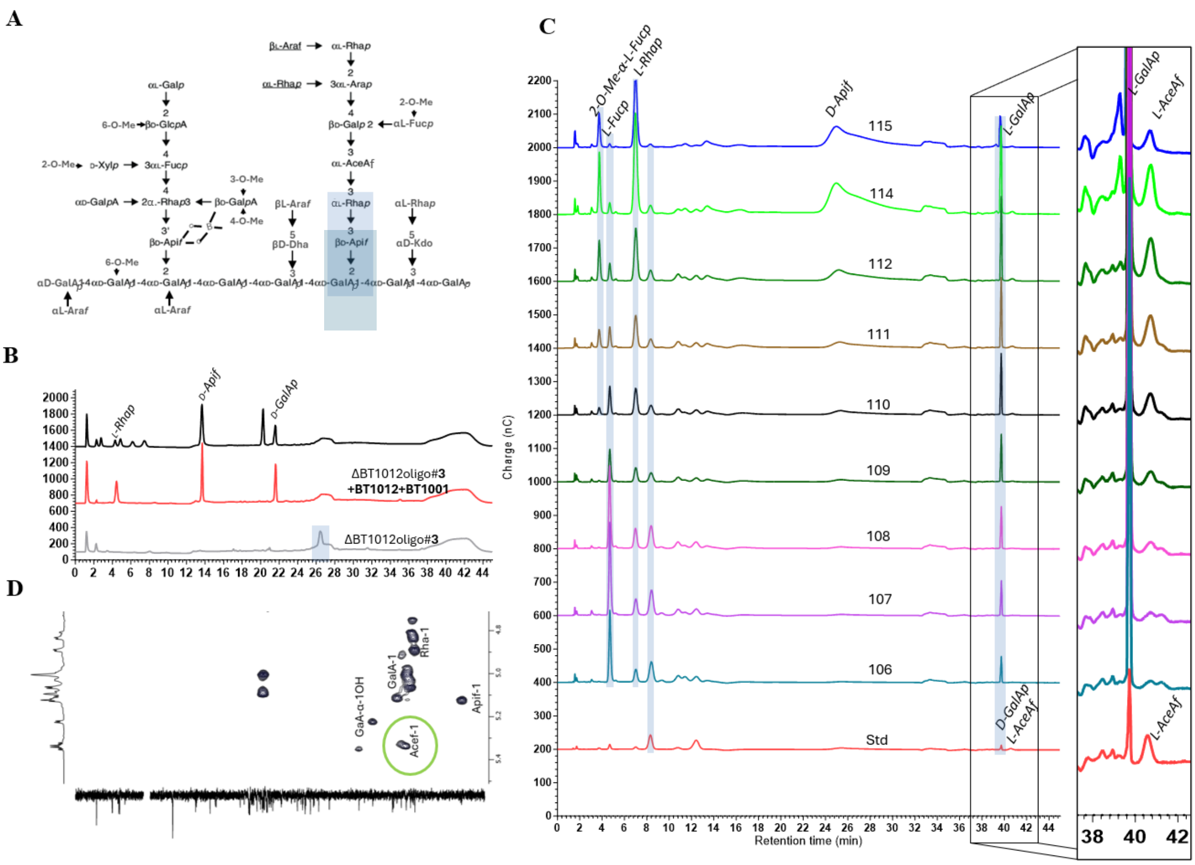

**Supplemental figure 1: Characterisation of secreted products of ΔBT1012 strain.** **A**: Structure of RG-II showing the context of ΔBT1012 oligo#3 **B**: Enzymatic profiling of ΔBT1012 oligo#3 using RG-II specific enzymes **C:** HPAEC-PAD showing the detection of L-AceA*f* in various purified fractions of ΔBT1012 products. **D:** NMR of ΔBT1012 smear products shows detection of $H^1$ signals of α-L-Ace*f* (Acef-1)

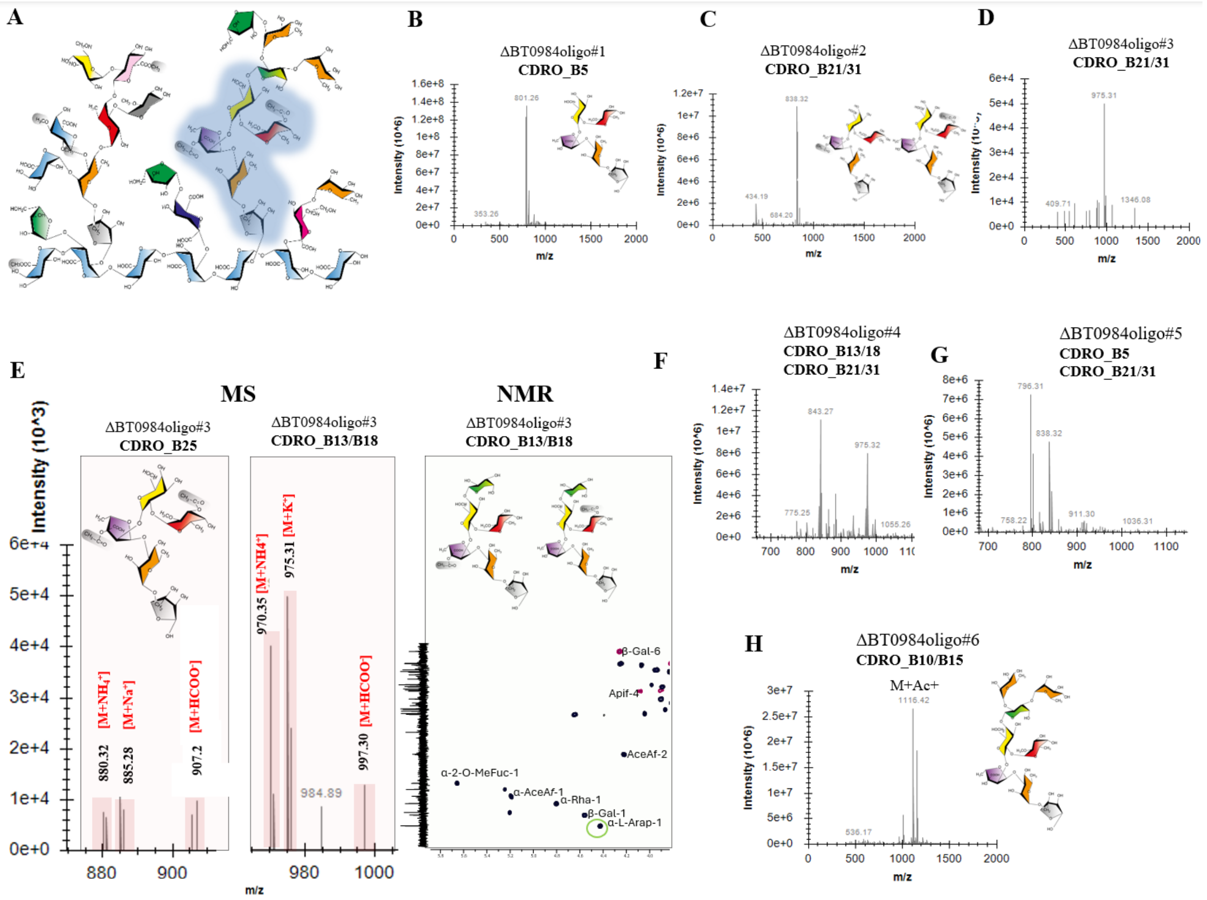


**Supplemental figure 2: Characterisation of secreted products of ΔBT0984 strain.** **A:** structure of RG-II showing the context of ΔBT0984 oligo#1 **B-H:** MS and NMR analyses of various other oligos generated by ΔBT0984

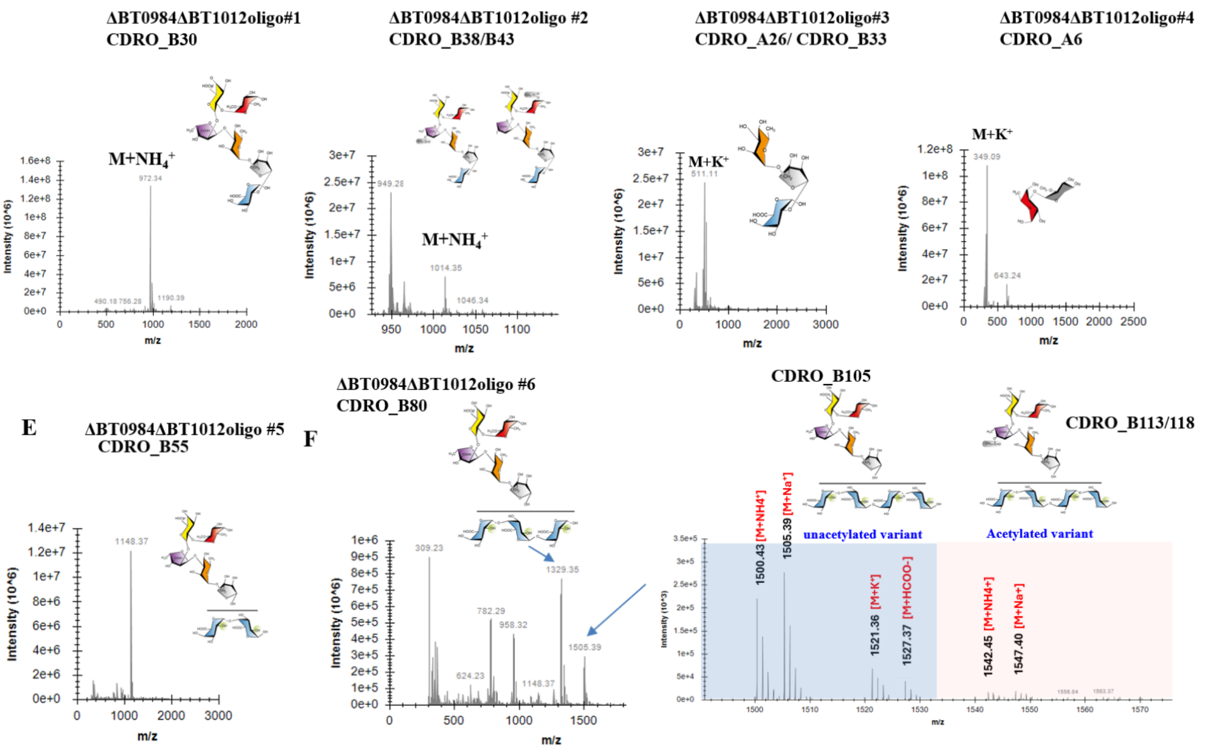


**Supplemental figure 3: MS analyses and identification of secreted products of ΔBT0984ΔBT1012 strain.** Data shows detection of masses consistent with the detection of RG-II-derived oligosaccharides with HG backbone components

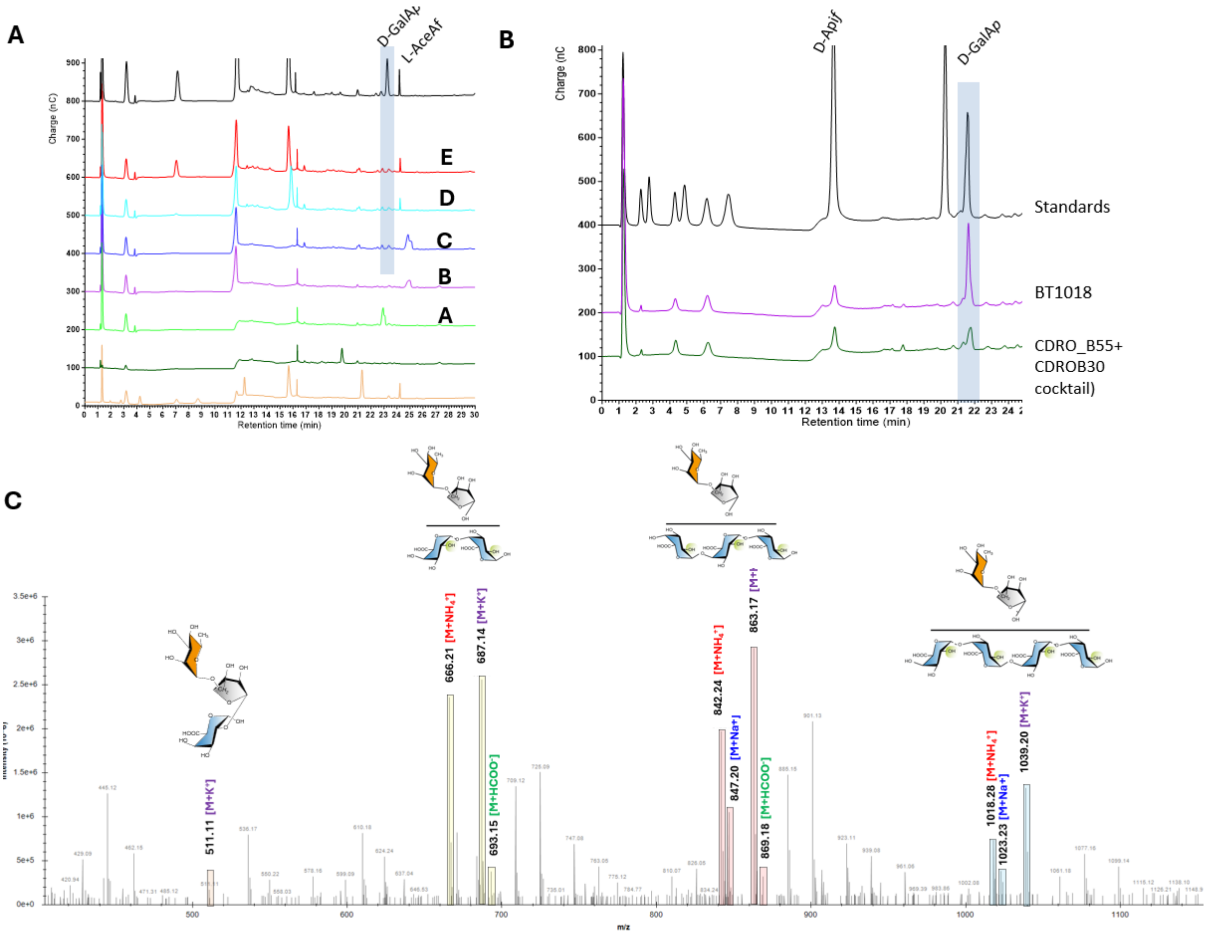


**Supplemental figure 4: Identification HG backbone-containing oligosaccharides generated by ΔBT0984ΔBT1012 and ΔBT1012 strains.** **A**: Enzymatic profiling of ΔBT0984ΔBT1012oligo#5 shows absence of D-GalAp**. B:** Release of D-GalAp from ΔBT0984ΔBT1012oligo#5 **C**: MS analyses of ΔBT1012 oligos shows detection of HG backbone-containing oligosaccharides

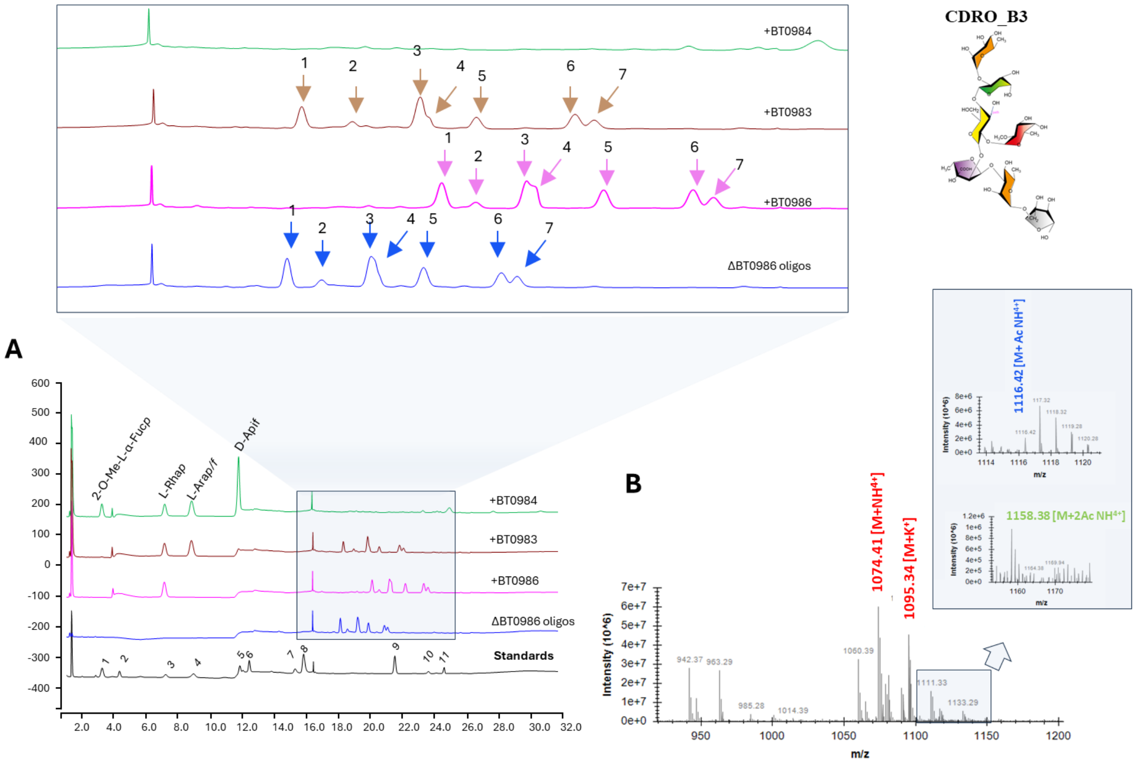


**Supplemental figure 5: HPAEC-PAD and MS detection of the range of oligosaccharides generated by ΔBT0986.** Partially purified oligosaccharides were treated with various side chain B-degrading enzymes. About 7 distinct HPAEC-PAD peaks from the mixture of partially purified ΔBT0986 oligos were found to be sensitive to BT0986 and BT0983 enzymes.

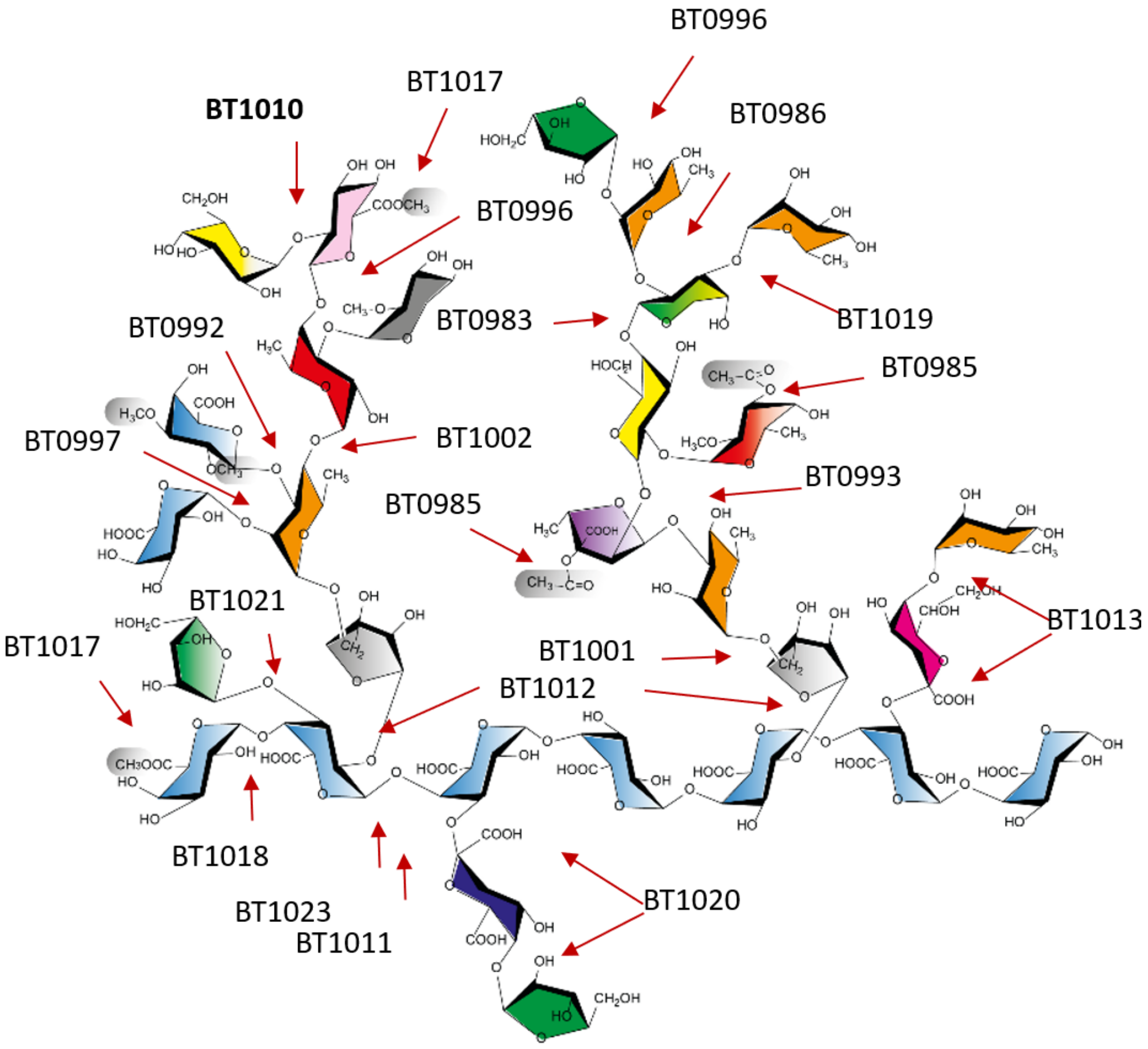


**Supplemental figure 6: Structure of apple RG-II showing components units and cleavage sites of various RG-II-degrading enzymes based on previous studies Ndeh *et al.,* 2017 [1] and Duan *et al.,* 2020 [2]**

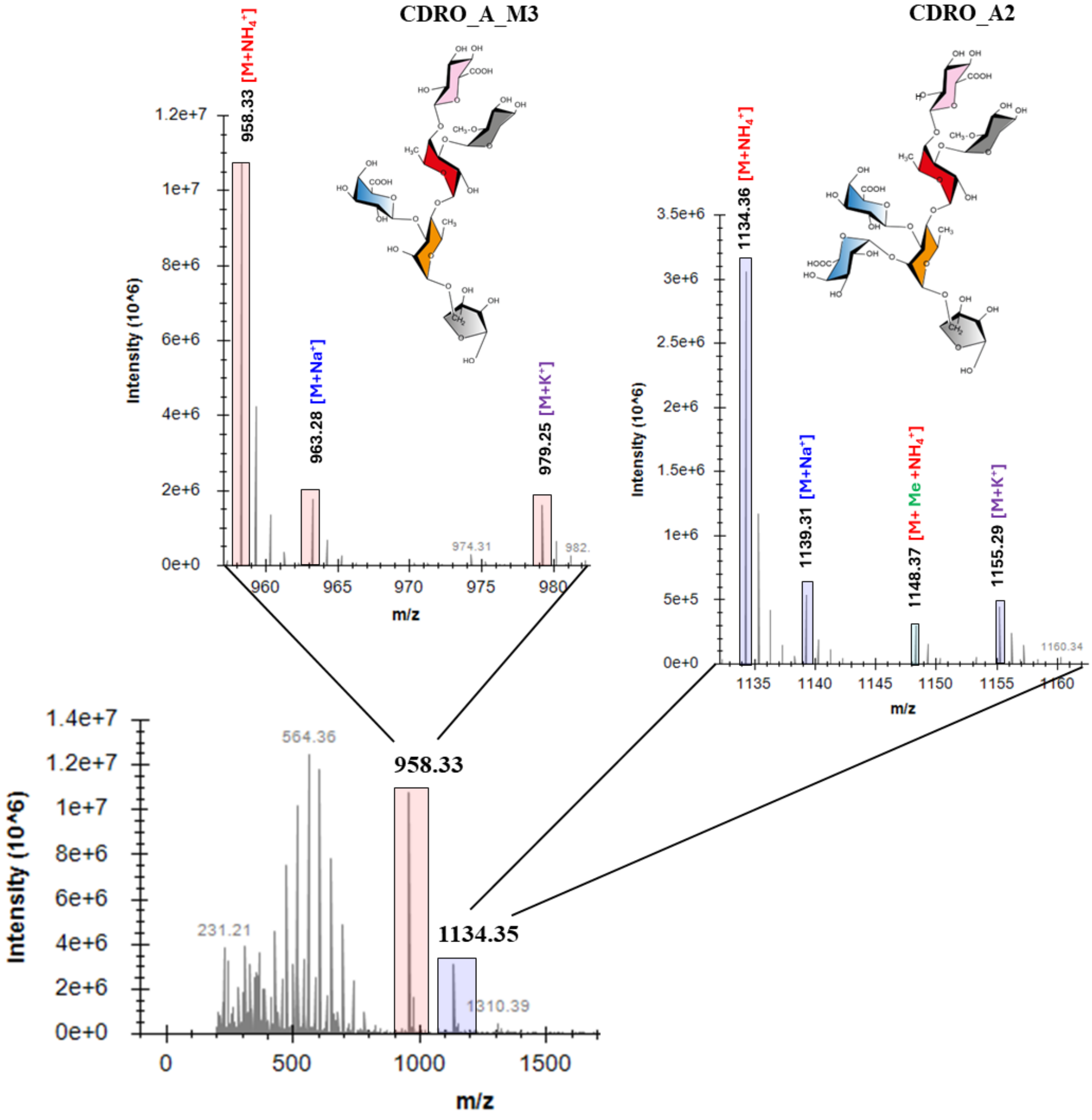


**Supplemental figure 7: MS analyse of major oligosaccharides produced by BT0996:Tn**. Growth supernatants were desalted, diluted with LC-MS grade water and later analysed by MS.

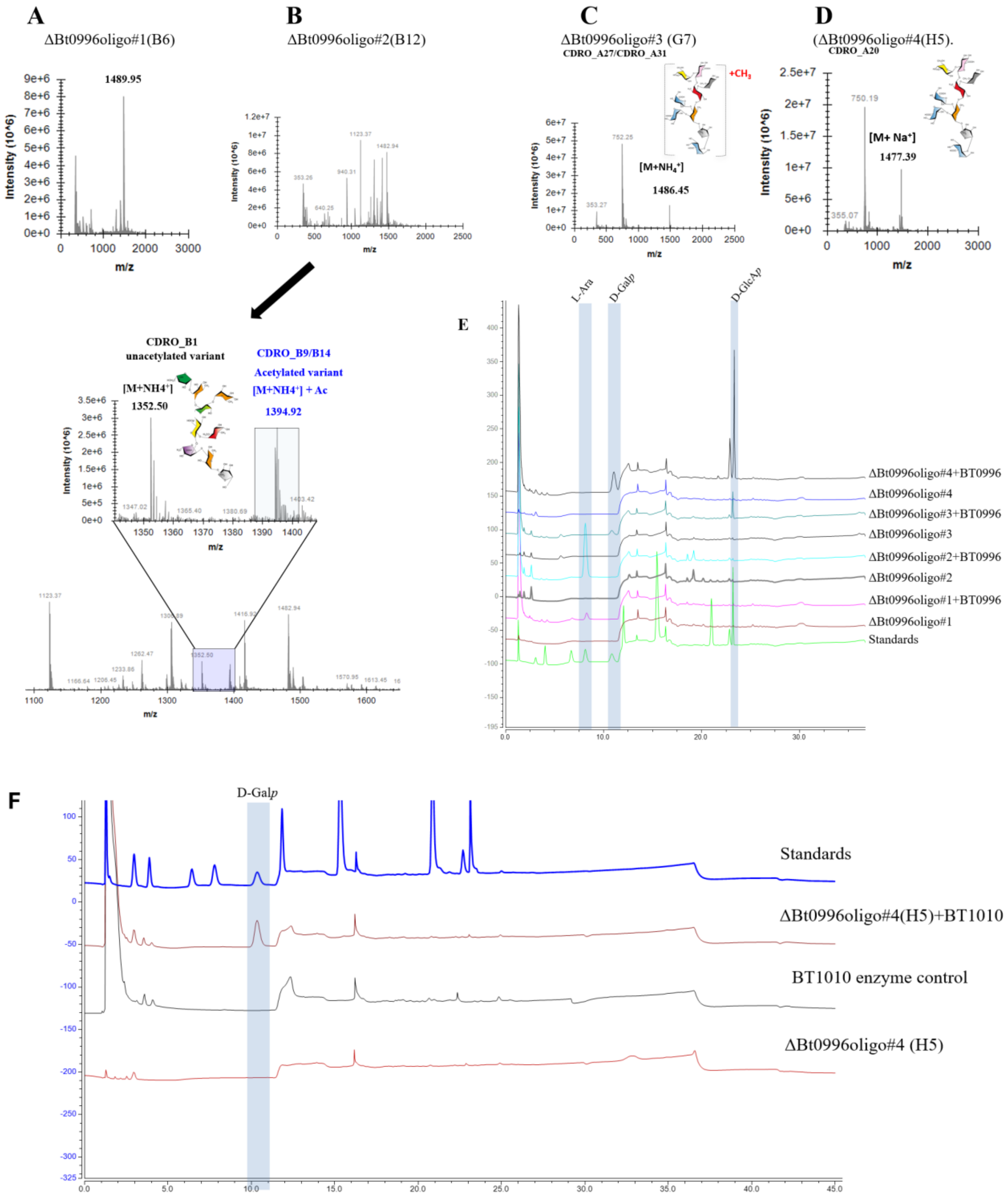


**Supplemental figure 8: MS analyses of major oligosaccharides produced by ΔBT0996**. The contents of culture medium after growth of the mutant on RG-II were resolved by SEC and fractions analysed by MS. **A-D:** MS of major fractions collected from SEC, **E:** Treatment of various SEC fractions with BT0996 recombinant enzyme showing release of L-Ara, D-GlcA*p* and D-Gal*p*. **F:** ΔBT0996oligo#3 and #4 both yielded peaks for D-Gal*p* in panel E, likely due to contamination with BT1010, hence purified BT1010 alone was used to treat ΔBt0996oligo#4 showing release of D-Gal*p*, providing further confirmation that the oligo is derived from side chain A

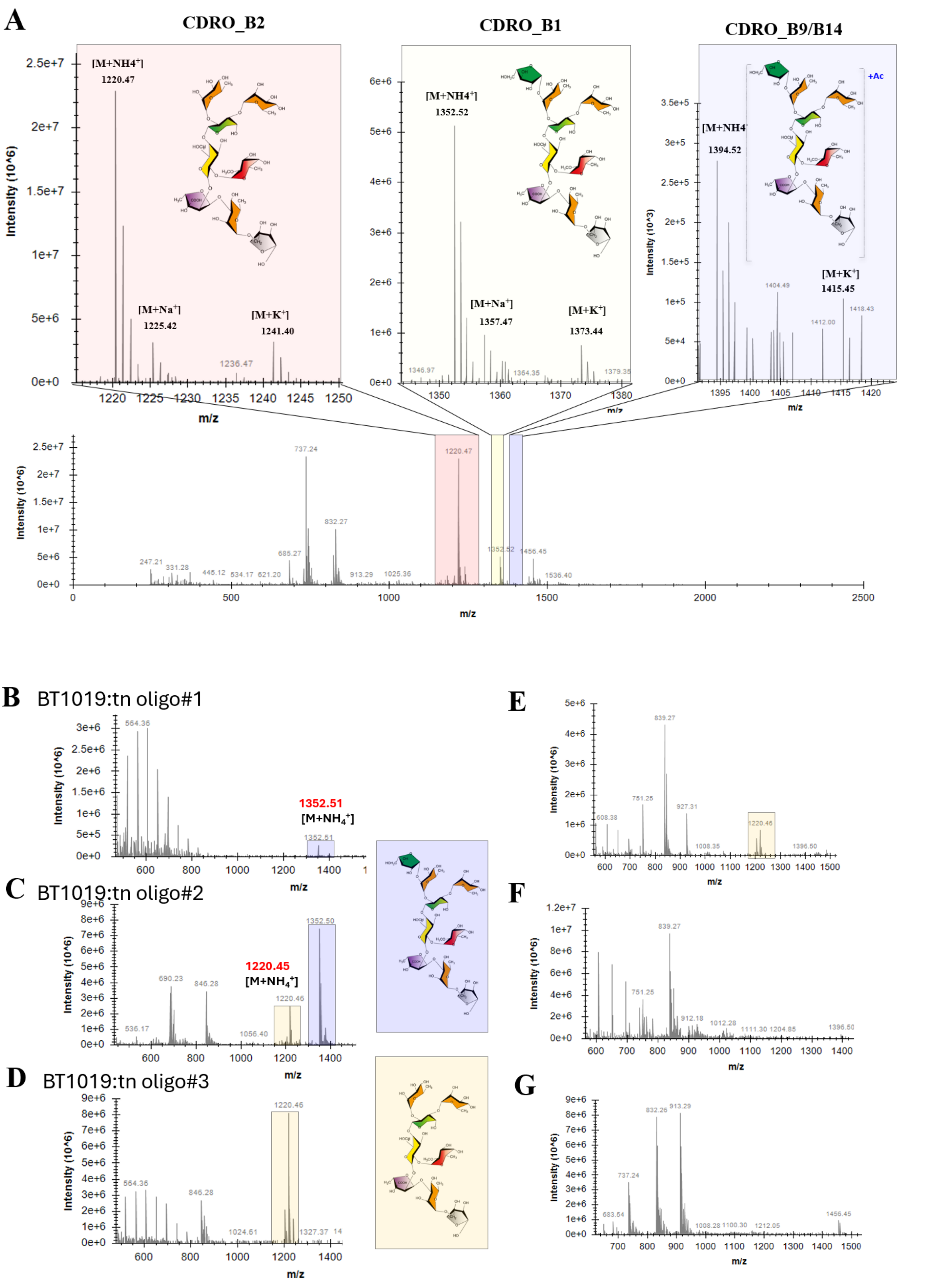


**Supplemental figure 9: MS analyses of major oligosaccharides generated by BT1019: Tn. A:** Cells were cultured in the presence of 1% apple RG-II. Growth supernatants were desalted, diluted with LC-MS grade water and later analysed by MS. **B-G:** MS analyses of full SEC fractions. Note that the RG-II source used here was *N. benthamiana*.

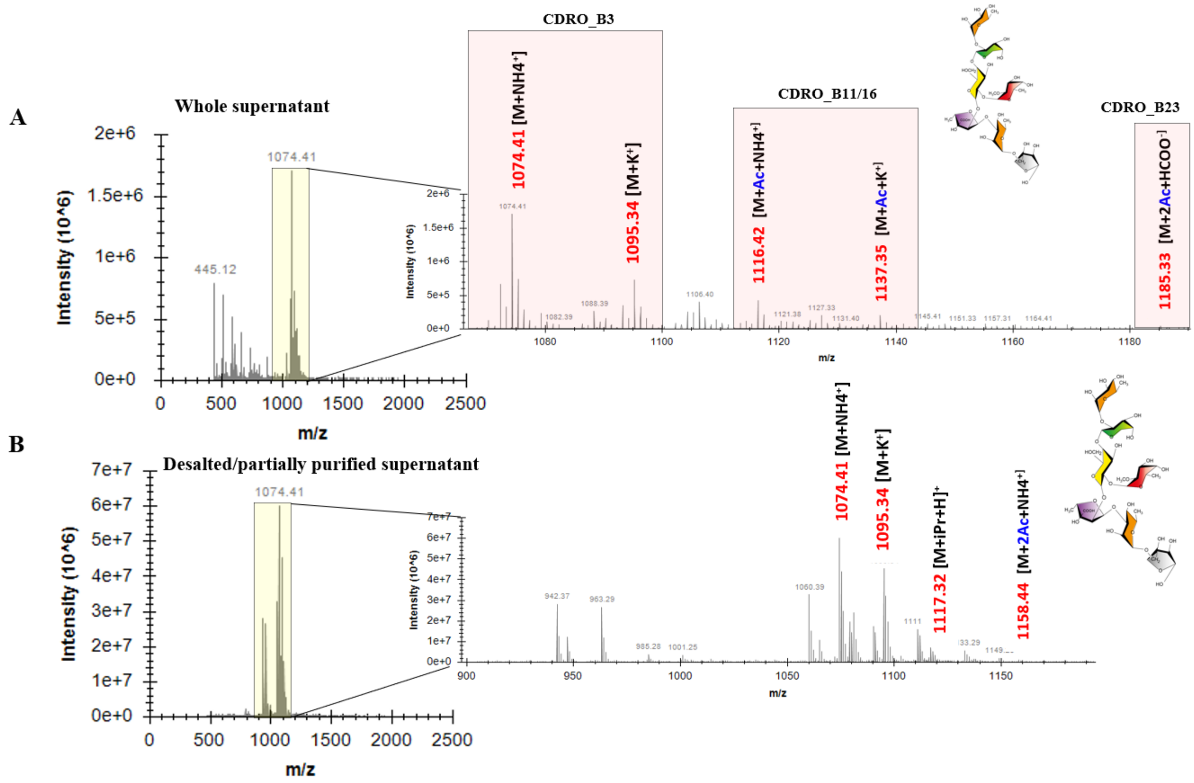


**Supplemental figure 10: MS analyses of major oligosaccharides produced by BT0986:Tn**. Growth supernatants were desalted, diluted with LC-MS grade water and later analysed by MS.

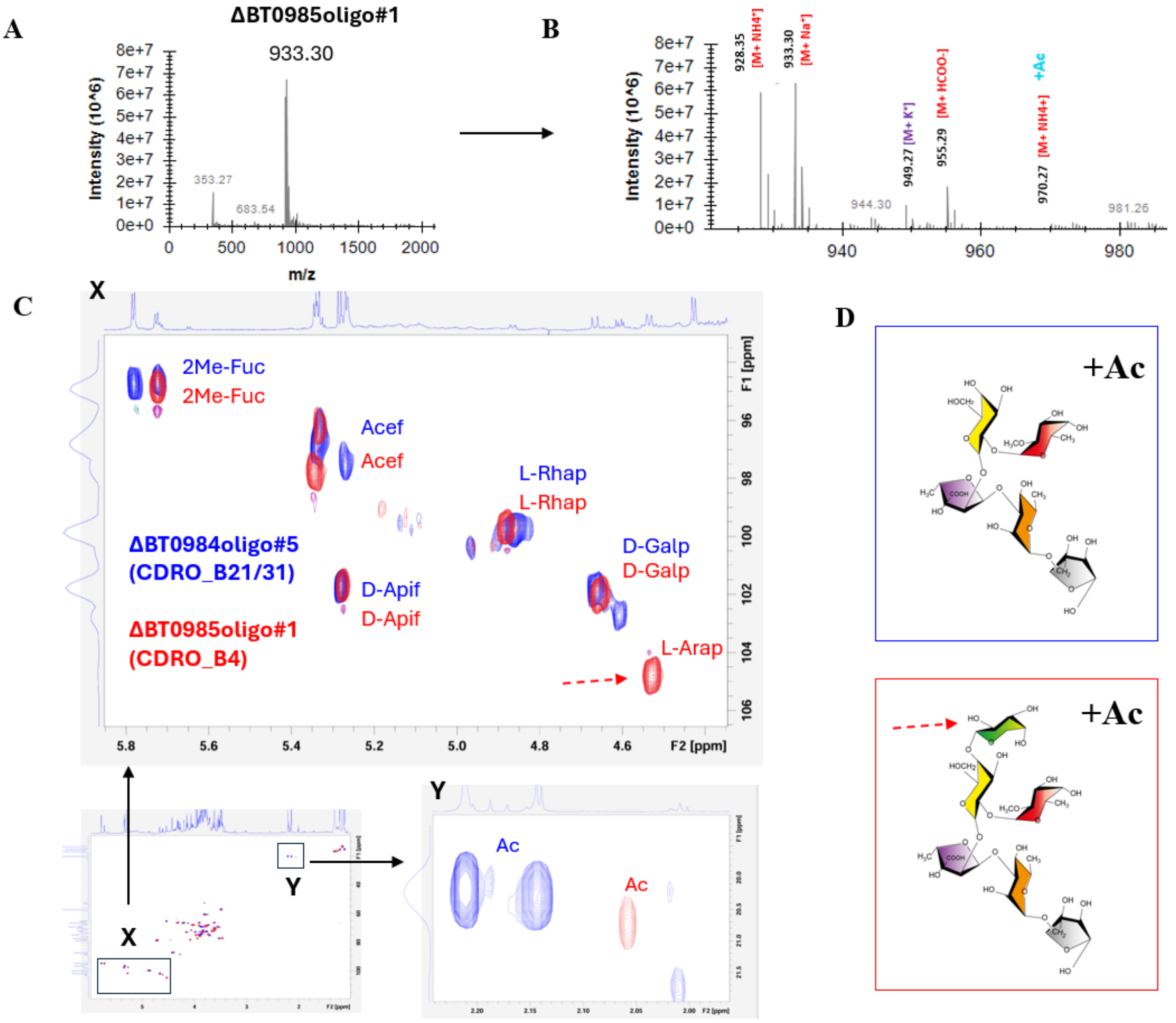


**Supplemental fig. 11: MS and NMR analyses of CDRO generated by ΔBT0985 strain**. **A**: MS analyses of showing detection of a major peak corresponding to **CDRO_B4**. **B**: Zoom-in of (A) showing various ionic adducts or other forms detected **C:** Overlay and comparison of NMR spectra of previously characterised **ΔBT0984oligo#2** (CDRO_B21/31) (blue) and **ΔBT0985oligo#1** (CDRO_B4) (red) showing the presence of L-Ara*p* in the latter. Acetyl groups were also detected in both analysed samples.

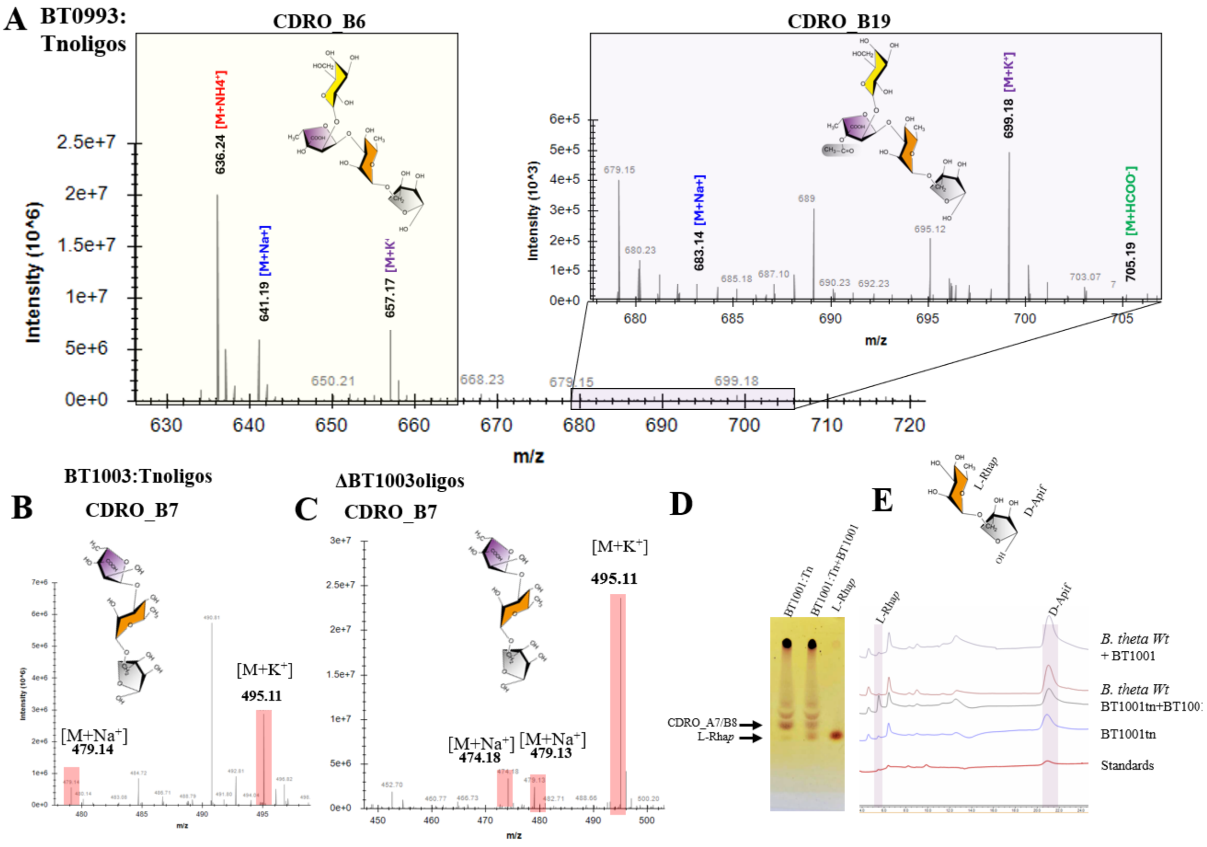


**Supplemental fig. 12: MS, TLC and HPAEC-PAD analyses of BT0993:Tn, BT1003:Tn, and BT1001:Tn oligos. A:** MS detection of **CDRO_B6** and **CDRO_B19** in desalted/partially purified **BT0993:Tn** supernatants **B**: MS detection of **CDRO_B7** in desalted/partially purified **BT1003:Tn** supernatants. **C**: MS detection of **CDRO_B7** in desalted/partially purified **ΔBT1003** supernatants **D**: TLC analyses of BT1001:Tn supernatants showing digestion of BT1001:Tn oligo (CDRO_A7/B8) to L-Rha*p*. **E**: HPAEC-PAD analyses of BT1001:Tn supernatants showing production of L-Rha*p* after digestion with BT1001.

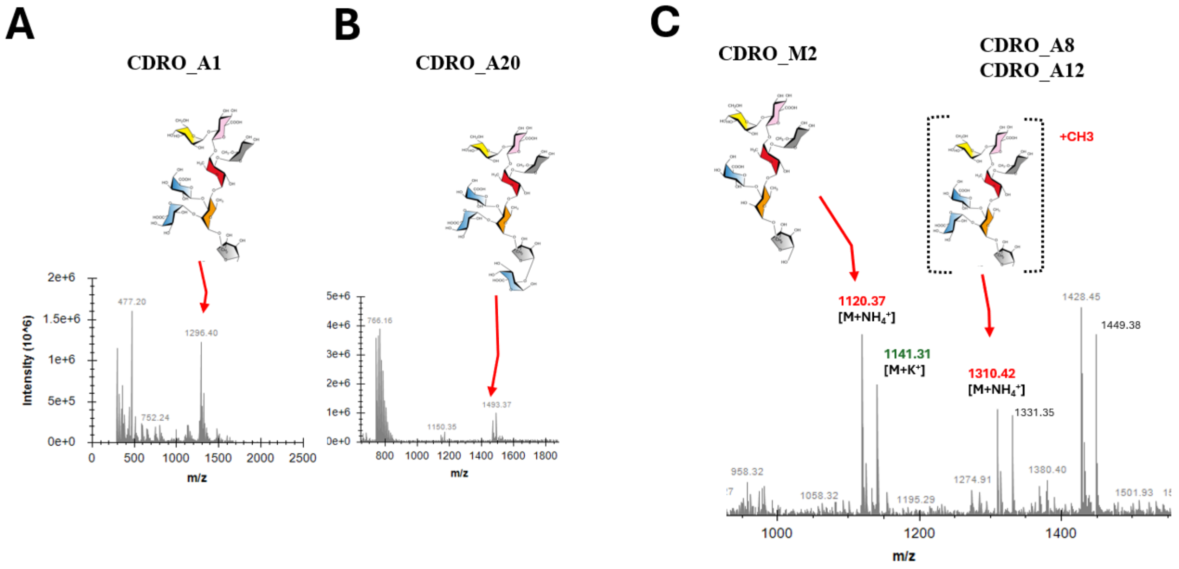


**Supplemental fig. 13: MS detection and analyses of BT1010:tn oligosaccharide products (A-C).** Growth supernatants were desalted, diluted with LC-MS grade water and later analysed by MS.

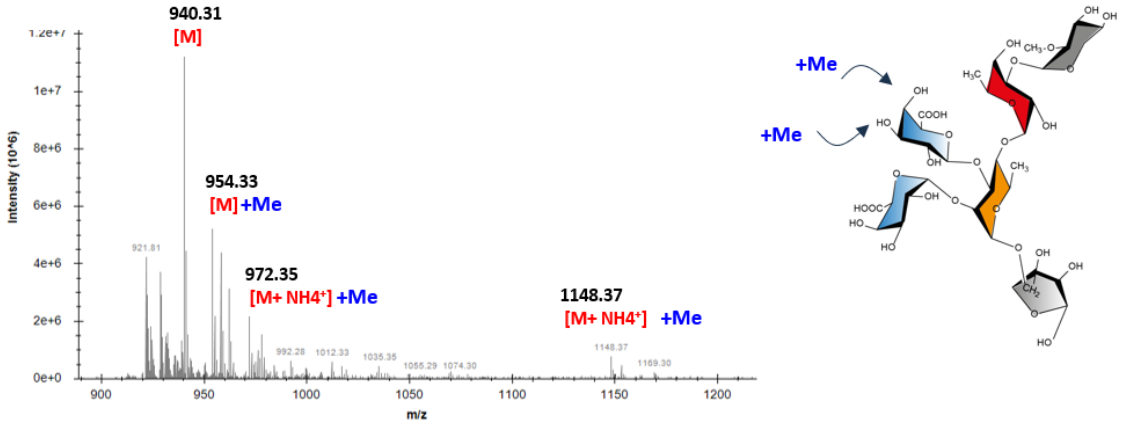


**Supplemental fig. 14: MS detection and analyses of BT0997:tn oligosaccharide products.** Growth supernatants were desalted, diluted with LC-MS grade water and later analysed by MS.

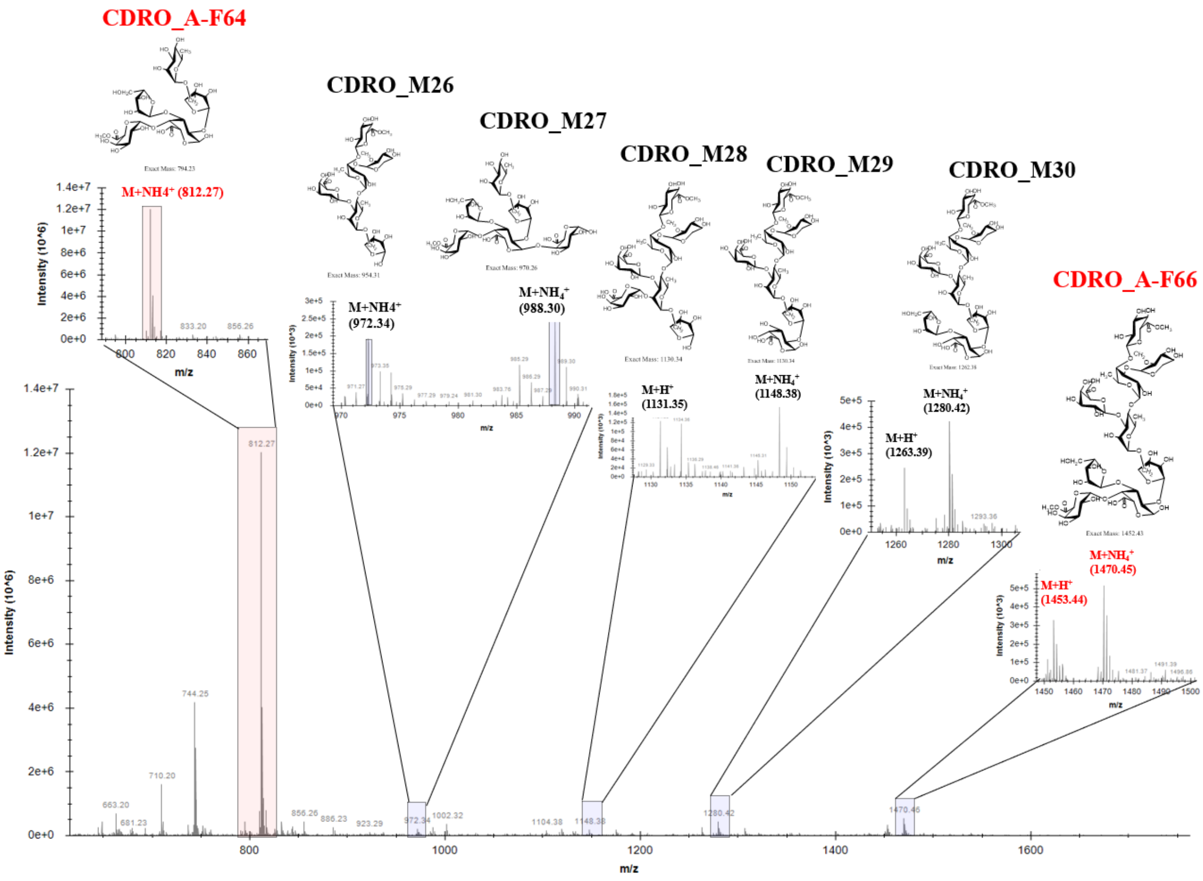


**Supplemental fig. 15: MS detection and analyses of ΔBT1017 oligosaccharide products.** Growth supernatants were desalted, diluted with LC-MS grade water and later analysed by MS. Oligos highlighted in red were previously purified and reported[2]. All oligos in black were newly identified in this study from further analyses.

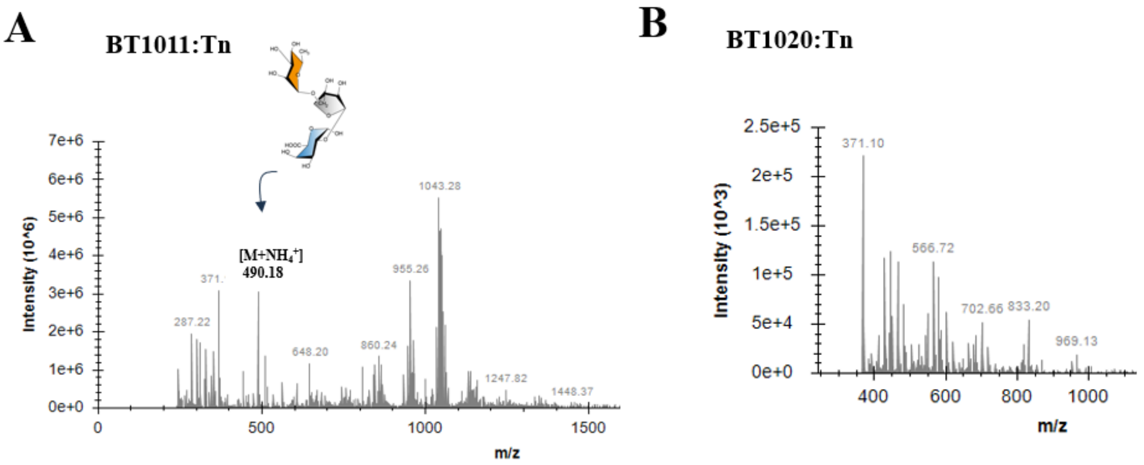


**Supplemental fig. 16: MS detection oligosaccharide products generated by BT1011:Tn and BT1020:Tn.** Growth supernatants were desalted, diluted with LC-MS grade water and analysed by MS.

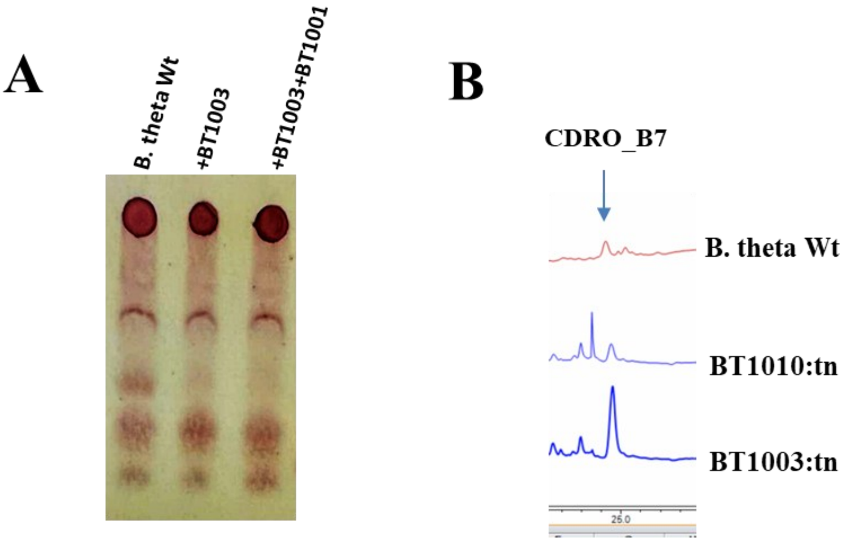


**Supplemental fig. 17: Confirming the secretion of CDRO_B7 by *B. theta* Wt. A:** Sensitivity of *B. theta* Wt -secreted CDRO_B7 to BT1003 and BT1001 enzymes confirms its identity – also see HPAEC detection in **Fig. 3E**. **B:** Comparing amounts secreted by *B. theta* Wt and BT1003:tn

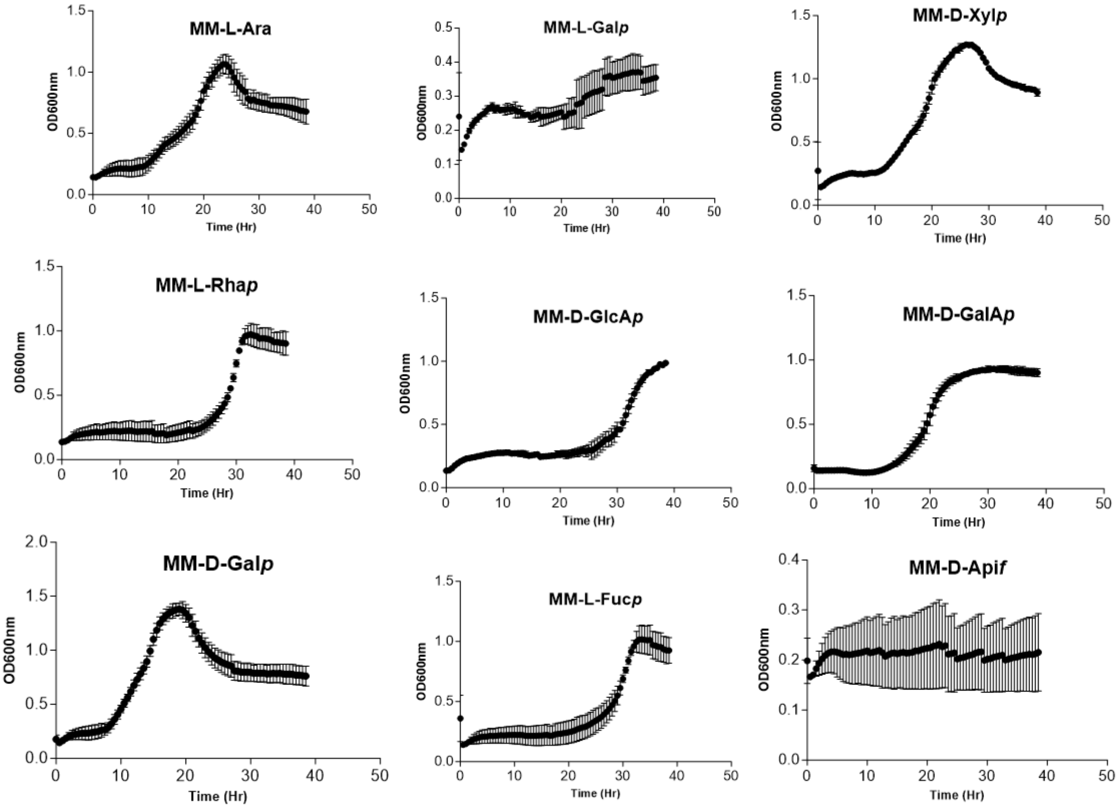


**Supplemental fig. 18: Growth of *B. theta* Wt on various RG-II-derived monosaccharides.** *B. theta Wt* cells were cultured in all cases on 1% substrate in minimal medium as sole carbon sources

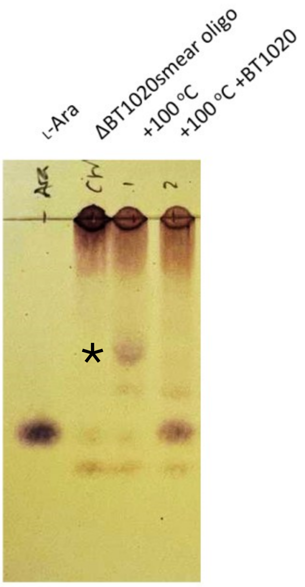


**Supplemental fig. 19: Effect of heat treatment on ΔBT1020smear oligo and subsequent treatment with BT1020 recombinant enzyme.** The latter shows the release or generation of L-Ara from the heat released product*

**A**

| | Gene | Prediction | Score |
|---|---|---|---|
| 1 | BT_0984 | SpI | score=7.27601 margin=4.21811 cleavage=25-26 |
| 2 | **BT_0985** | **SpII** | score=18.11 margin=2.8749 cleavage=19-20 Pos+2=L |
| 3 | BT_0986 | SpI | score=13.1534 margin=10.38197 cleavage=25-26 |
| 4 | BT_0992 | SpI | score=14.9997 margin=15.200613 cleavage=19-20 |
| 5 | BT_0993 | SpI | score=6.4885 margin=6.689413 cleavage=20-21 |
| 6 | BT_0996 | SpI | score=7.61798 margin=6.12371 cleavage=27-28 |
| 7 | BT_0997 | CYT | score=-0.200913 |
| 8 | BT_1001 | SpI | score=5.23856 margin=5.439473 cleavage=15-16 |
| 9 | BT_1002 | SpI | score=5.60203 margin=5.802943 cleavage=19-20 |
| 10 | BT_1003 | SpI | score=20.3956 margin=14.71186 cleavage=20-21 |
| 11 | BT_1010 | SpI | score=9.56058 margin=4.05061 cleavage=23-24 |
| 12 | **BT_1011** | **SpII** | score=8.68617 margin=2.55011 cleavage=19-20 Pos+2=T |
| 13 | BT_1012 | SpI | score=17.3671 margin=17.568013 cleavage=22-23 |
| 14 | BT_1013 | SpI | score=12.1332 margin=12.334113 cleavage=23-24 |
| 15 | BT_1017 | SpI | score=10.6088 margin=10.809713 cleavage=18-19 |
| 16 | BT_1018 | SpI | score=6.41352 margin=6.614433 cleavage=21-22 |
| 17 | BT_1019 | CYT | score=-0.200913 |
| 18 | BT_1020 | SpI | score=11.3618 margin=11.562713 cleavage=22-23 |
| 19 | BT_1021 | CYT | score=-0.200913 |
| 20 | BT_1023 | SpI | score=14.0935 margin=14.294413 cleavage=23-24 |

**B**

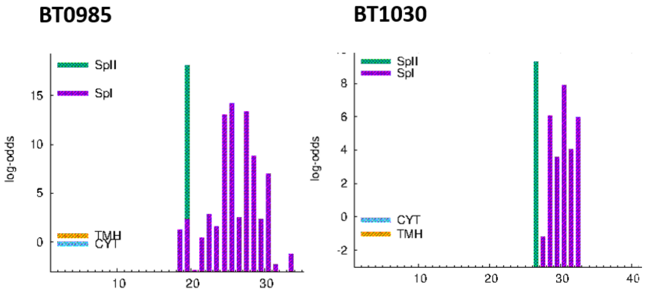


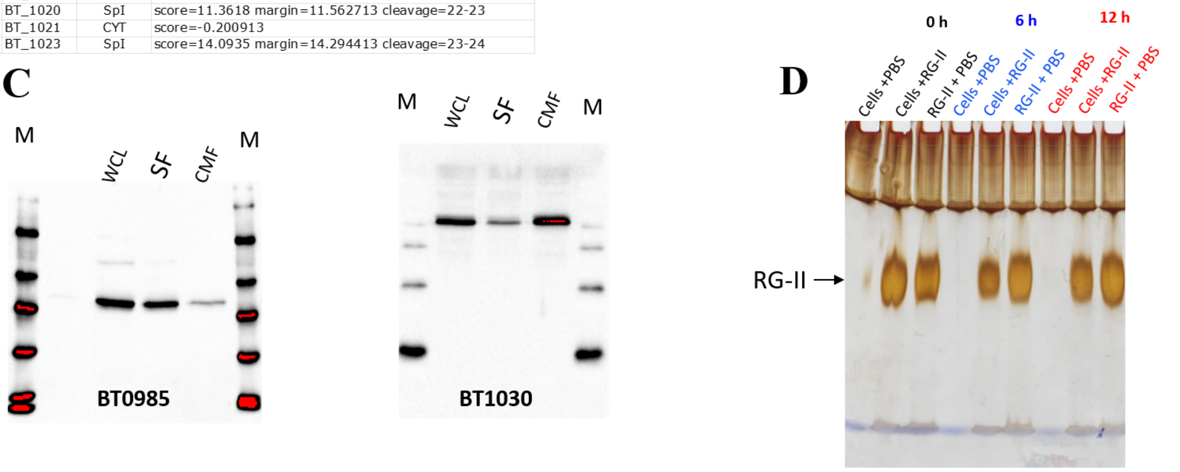


**Supplemental fig. 20: Cellular localisation of RG-II-degrading BT0985 enzyme. A:** Bioinformatic prediction of the cellular localisation BT0985 and other RG-II CAZymes using LipoP tool[3] The protein BT1011, although containing a SpII signal acts on BT1020smear oligo (**Fig. 4G**) secreted from the periplasm in ΔBT1020 and hence is highly likely to be periplasmic. **B:** LipoP prediction graph for BT0985 alongside BT1030 protein previously confirmed to be localised extracellularly on the cell surface[1]. Both show strong SpII signals. **C:** Biochemical detection and localisation of BT0985 and BT1030. *B. theta* Wt cells were sonicated and the resulting whole cell lysate (WCL) ultracentrifuged into soluble fractions (SF) and cell membrane fractions (CMF). Fractions were resolved by SDS-PAGE followed by western blotting using polyclonal antibodies against both proteins. **D:** Aerobic whole cell assays to probe cell-surface endolytic degradation of RG-II. Whole cells of *B. theta* Wt were incubated with apple RG-II from 0 to 12 h followed by acrylamide gel electrophoresis of supernatants as described previously[4]

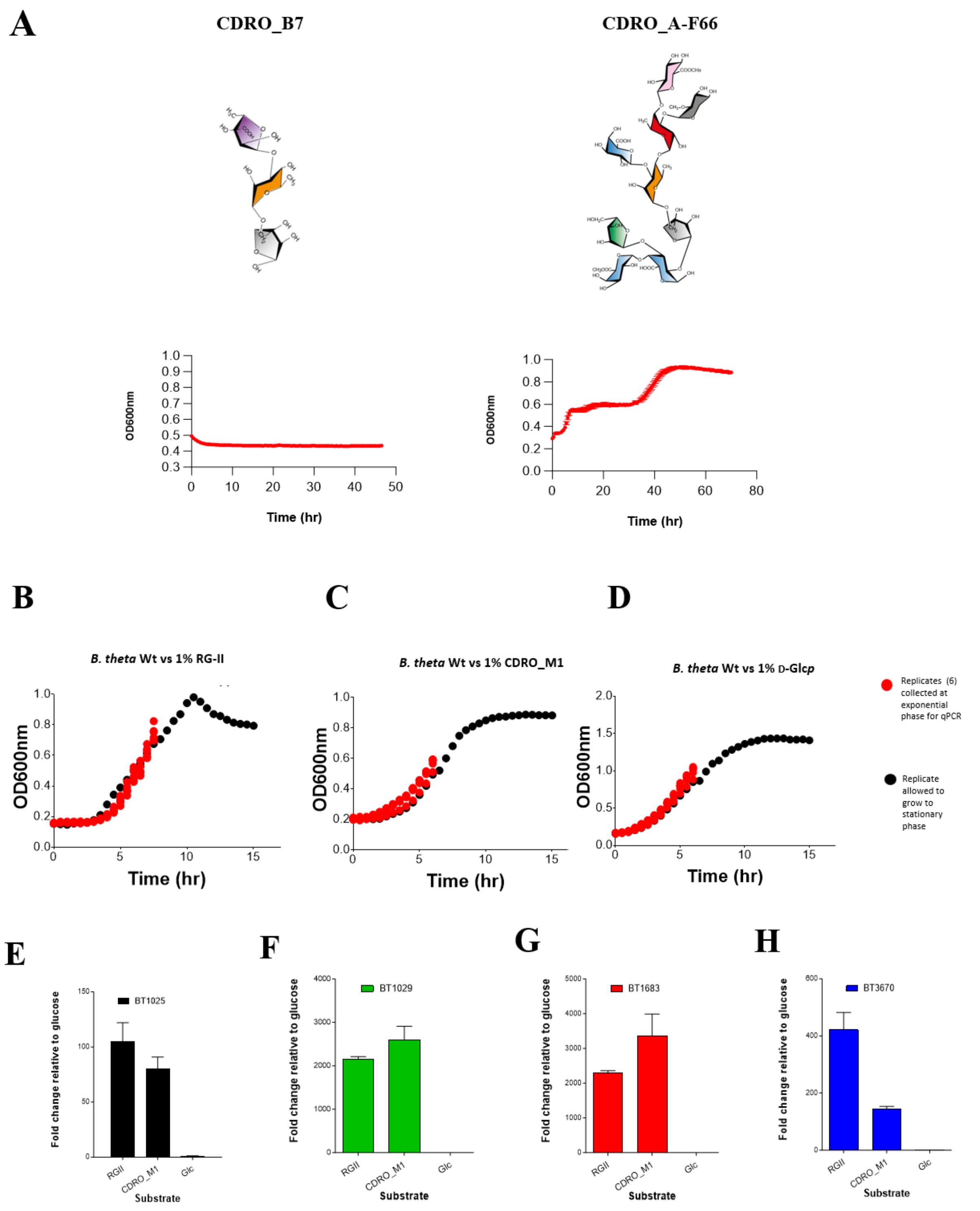


**Supplemental fig. 21: Growth of *B. theta* Wt on various CDROs and RG-II and qPCR quantification of RG-II transporter genes BT1025, BT1029, BT1683 and BT3670**

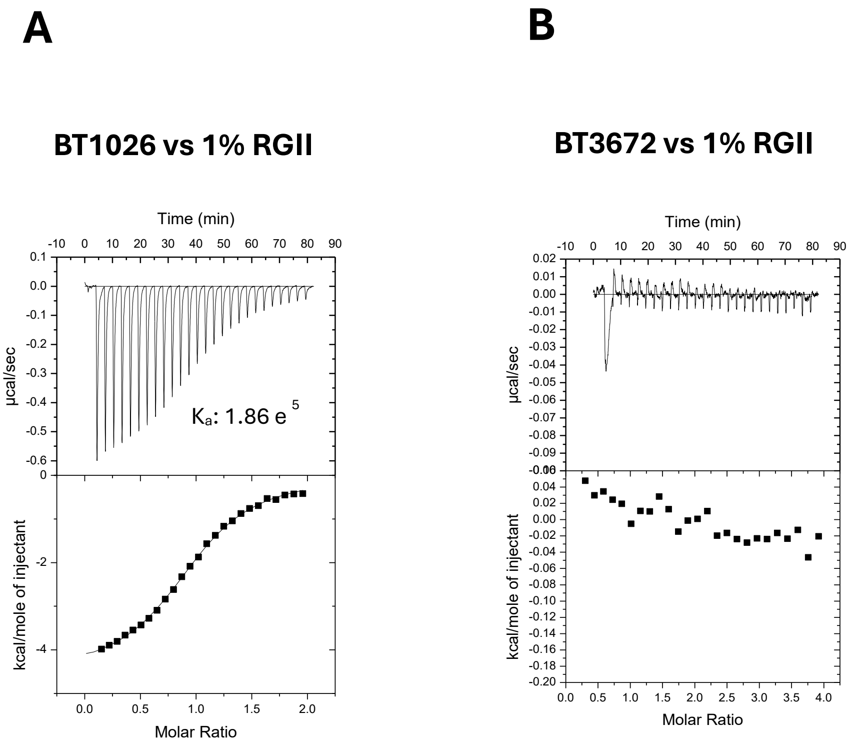


**Supplemental fig. 22: ITC traces showing binding of BT1026 but not BT3672 to RG-II.**

.

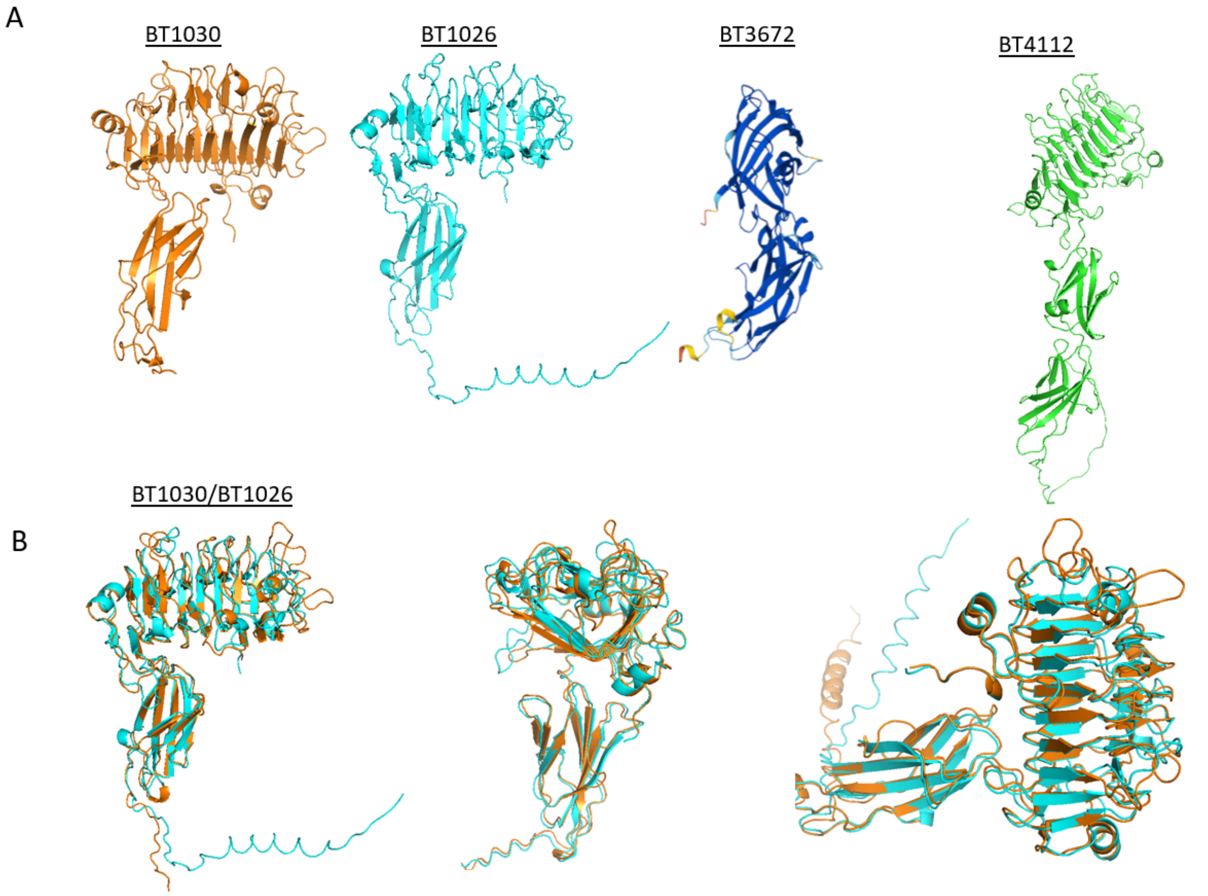


**Supplemental fig. 23: Structural comparison of various predicted surface glycan binding proteins from RG-II PUL and BT4112 from the HG PUL[5,6]. A:** Individual AlphaFold predicted structures are shown. Data shows similarities in structural folds between HG binding proteins in the RG-II PUL and a predicted HG binding protein BT4112. **B:** Alignment of BT1030 and BT1026 structures shows strong structural similarity between structures (~73.2%, rmsd 1.11Å) consistent with their similar binding preferences.

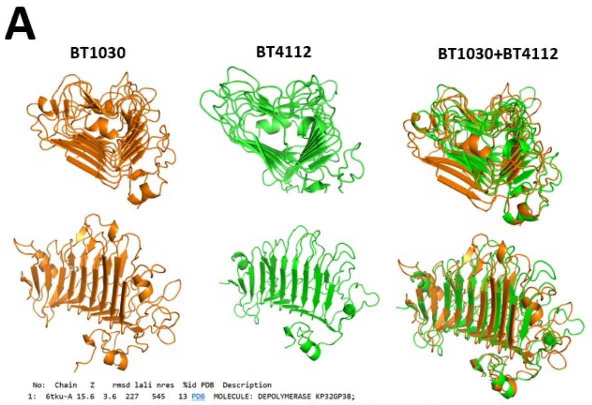


| No: | Chain | Z | rmsd | lali | nres | %id | PDB | Description |
|---|---|---|---|---|---|---|---|---|
| 1: | 6tku-A | 15.6 | 3.6 | 227 | 545 | 13 | PDB | MOLECULE: DEPOLYMERASE KP32GP38; |
| 2: | 6c72-A | 15.4 | 6.2 | 256 | 579 | 11 | PDB | MOLECULE: PARTICLE-ASSOCIATED GLYCOSIDE HYDROLASE; |
| 3: | 7chu-A | 15.4 | 10.8 | 259 | 517 | 10 | PDB | MOLECULE: PUTATIVE PECTIN LYASE; |
| 4: | 4mr0-A | 15.4 | 4.0 | 245 | 456 | 9 | PDB | MOLECULE: PLASMIN AND FIBRONECTIN-BINDING PROTEIN A; |
| 5: | 6e1r-A | 15.3 | 7.1 | 255 | 544 | 9 | PDB | MOLECULE: TAILSPIKE PROTEIN; |
| 6: | 5zru-A | 15.3 | 5.9 | 258 | 576 | 11 | PDB | MOLECULE: ALPHA-1,3-GLUCANASE; |
| 7: | 6kqs-A | 15.2 | 6.9 | 261 | 620 | 8 | PDB | MOLECULE: LACTO-N-BIOSIDASE; |
| 8: | 1bhe-A | 14.9 | 3.4 | 229 | 376 | 10 | PDB | MOLECULE: POLYGALACTURONASE; |
| 9: | 5jsd-A | 14.7 | 10.8 | 245 | 548 | 11 | PDB | MOLECULE: PHIAB6 TAILSPIKE; |
| 10: | 6nw9-C | 14.7 | 16.4 | 251 | 619 | 11 | PDB | MOLECULE: TAILSPIKE PROTEIN; |
| 11: | 5w5p-A | 14.7 | 11.3 | 252 | 610 | 12 | PDB | MOLECULE: TAIL FIBER PROTEIN; |
| 12: | 5gt5-A | 14.6 | 3.8 | 233 | 444 | 10 | PDB | MOLECULE: PECTATE LYASE; |
| 13: | 7eea-C | 14.6 | 5.3 | 262 | 646 | 10 | PDB | MOLECULE: SHORT-TAILED CYANOPHAGE TAILSPIKE RECEPTOR-BINDIN |
| 14: | 7ukj-C | 14.5 | 9.1 | 256 | 622 | 14 | PDB | MOLECULE: GENE 14 PROTEIN; |
| 15: | 9buj-A | 14.4 | 3.6 | 244 | 434 | 10 | PDB | MOLECULE: PECTATE LYASE; |
| 16: | 1pxz-A | 14.4 | 4.1 | 240 | 346 | 8 | PDB | MOLECULE: MAJOR POLLEN ALLERGEN JUN A 1; |
| 17: | 1rmg-A | 14.4 | 4.0 | 246 | 422 | 7 | PDB | MOLECULE: RHAMNOGALACTURONASE A; |
| 18: | 6kfn-A | 14.1 | 3.6 | 221 | 298 | 15 | PDB | MOLECULE: ALGINATE LYASE; |
| 19: | 1air-A | 14.1 | 3.8 | 227 | 352 | 11 | PDB | MOLECULE: PECTATE LYASE C; |
| 20: | 4c2l-A | 14.0 | 3.5 | 222 | 372 | 11 | PDB | MOLECULE: ENDO-XYLOGALACTURONAN HYDROLASE A; |
| 21: | 1idj-A | 13.9 | 3.7 | 224 | 359 | 12 | PDB | MOLECULE: PECTIN LYASE A; |
| 22: | 4nk6-A | 13.9 | 3.7 | 233 | 424 | 9 | PDB | MOLECULE: POLY(BETA-D-MANNURONATE) C5 EPIMERASE; |
| 23: | 5olq-C | 13.9 | 3.9 | 248 | 427 | 11 | PDB | MOLECULE: RHAMNOGALACTURONAN LYASE; |
| 24: | 3suc-A | 13.8 | 3.8 | 234 | 767 | 10 | PDB | MOLECULE: PRENECK APPENDAGE PROTEIN; |
| 25: | 7b8b-B | 13.6 | 3.7 | 223 | 367 | 10 | PDB | MOLECULE: POLYGALACTURONASE ADPG2; |
| 26: | 6tgf-D | 13.4 | 15.2 | 256 | 687 | 8 | PDB | MOLECULE: EXOPOLYSACCHARIDE BIOSYNTHESIS PROTEIN; |
| 27: | 5lw3-A | 13.4 | 3.6 | 229 | 381 | 9 | PDB | MOLECULE: POLY(BETA-D-MANNURONATE) C5 EPIMERASE 6; |
| 28: | 3vst-A | 13.3 | 3.5 | 239 | 638 | 8 | PDB | MOLECULE: XYLOSIDASE; |
| 29: | 5mqp-E | 13.3 | 4.0 | 250 | 602 | 11 | PDB | MOLECULE: GLYCOSIDE HYDROLASE BT_1002; |
| 30: | 5w6h-A | 13.3 | 18.1 | 261 | 697 | 10 | PDB | MOLECULE: TAILSPIKE PROTEIN 4; |
| 31: | 7xy1-A | 13.3 | 6.8 | 243 | 512 | 12 | PDB | MOLECULE: TAIL FIBER PROTEIN; |
| 32: | 7o78-B | 13.1 | 3.6 | 245 | 492 | 9 | PDB | MOLECULE: POLYSACCHARIDE LYASE FROM PSEUDOPEDOBACTER SALTAN |
| 33: | 1hg8-A | 13.1 | 3.8 | 220 | 349 | 12 | PDB | MOLECULE: ENDOPOLYGALACTURONASE; |
| 34: | 7vz3-B | 13.0 | 15.6 | 258 | 754 | 7 | PDB | MOLECULE: DEPOLYMERASE; |
| 35: | 8yk1-A | 13.0 | 3.5 | 244 | 810 | 14 | PDB | MOLECULE: ALPHA-GALACTOSIDASE; |
| 36: | 2uve-A | 13.0 | 3.9 | 241 | 571 | 11 | PDB | MOLECULE: EXOPOLYGALACTURONASE; |
| 37: | 7o17-A | 12.9 | 3.5 | 214 | 403 | 9 | PDB | MOLECULE: PROBABLE ABC TRANSPORTER BINDING PROTEIN NOSD; |
| 38: | 3lmw-A | 12.9 | 3.8 | 234 | 472 | 7 | PDB | MOLECULE: IOTA-CARRAGEENASE, CGIA; |

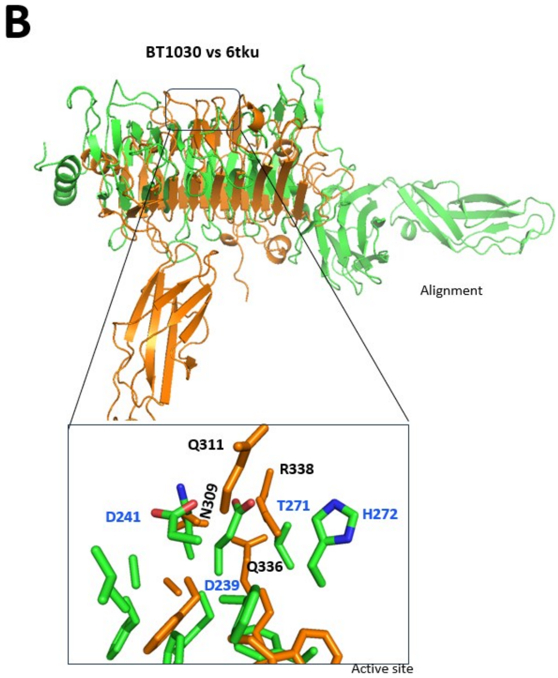


**Supplemental fig. 24: Structural comparison of BT1030 with BT4112 and capsule-specific depolymerase (PDB:6tku) from Klebsiella phage. A:** Alignment of BT1030 and BT4112 structures shows strong structural similarity (~72%, rmsd 2.9Å) **B:** Structurally similar homologues of BT1030 searched on PDBefold. The top hit was identified as a capsule-specific depolymerase (PDB:6tku) from Klebsiella phage. **C:** Alignment of BT1030 and 6tku shows fold conservation however, there was no conservation of active site residues (highlighted in blue).

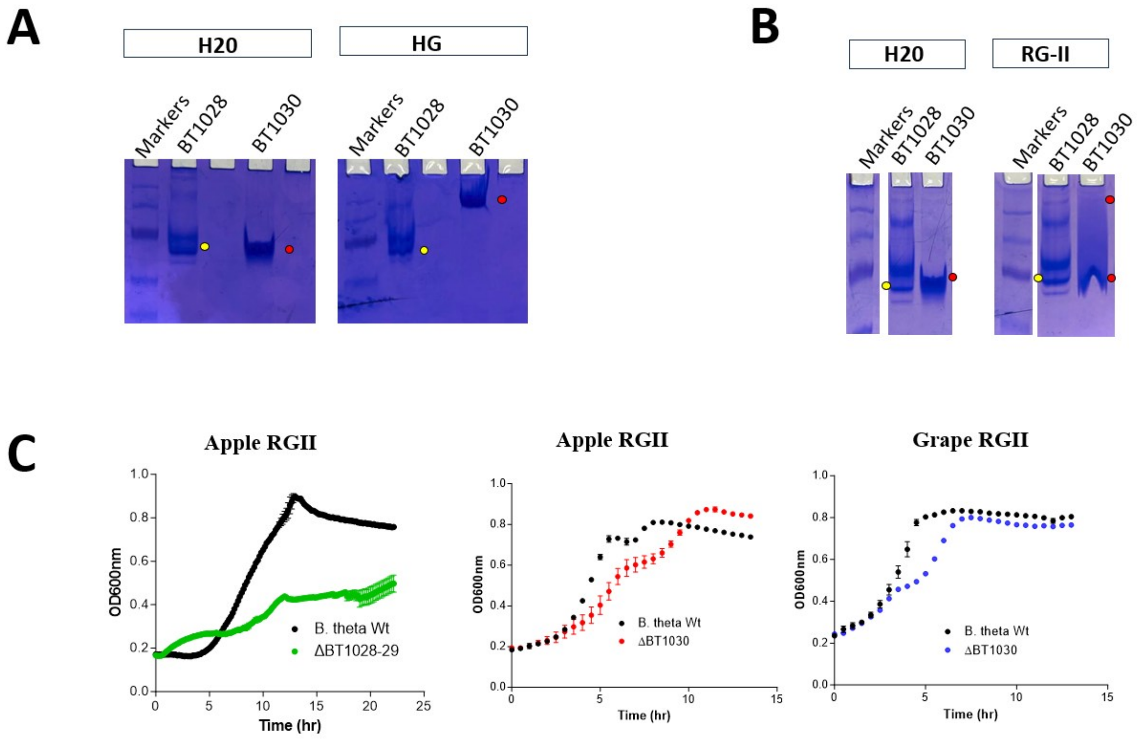


**Supplemental fig. 25: Polysaccharide binding assays and genetic experiments with elements of the RG-II surface acquisition apparatus of *B. theta***. NAGE experiments to probe for BT1028 (*sus*D-like) binding to HG (**A**) and RG-II (**B**) shows no retardation effect compared to control protein BT1030 that binds HG and RG-II. **C:** Growth of knockout mutants of BT1030 and BT1028-29 (double deletion mutant) on apple and wine RG-II.

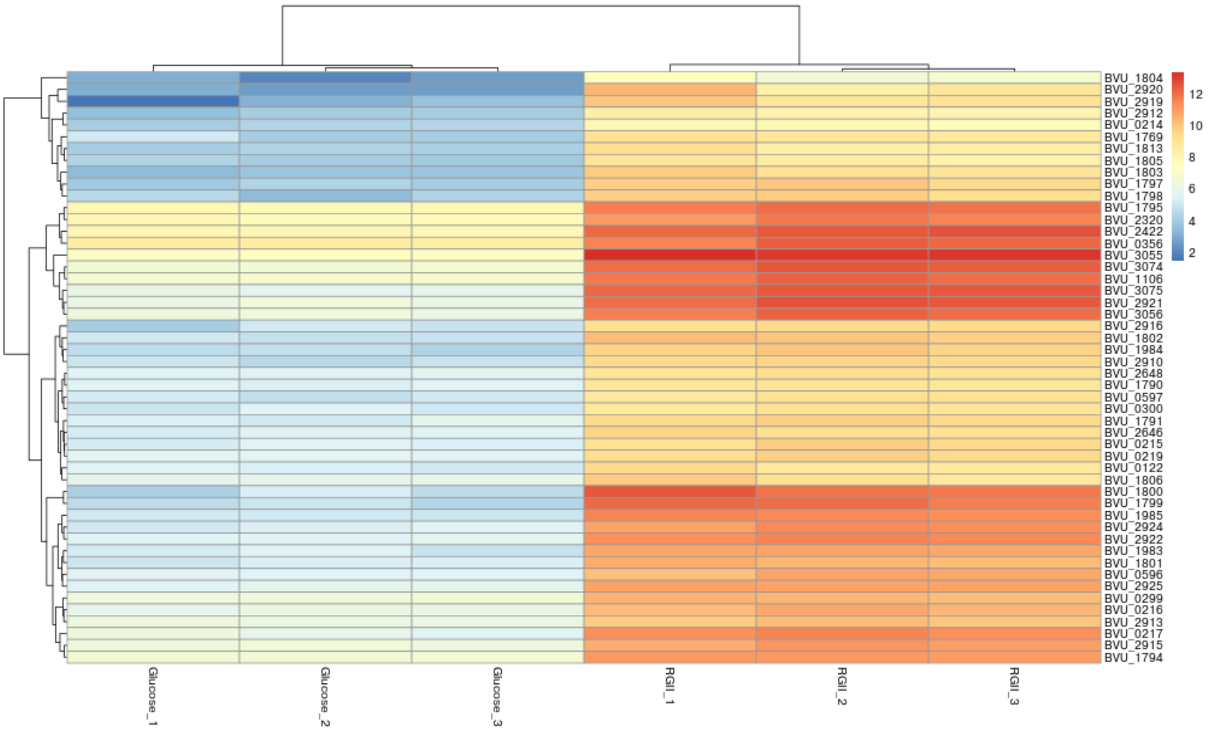


**Supplemental fig. 26: Heat map showing a summary of the most upregulated genes ($log_2$ normalised counts) in *P. vulgatus* when cultured in with apple RG-II as a sole carbon source.**

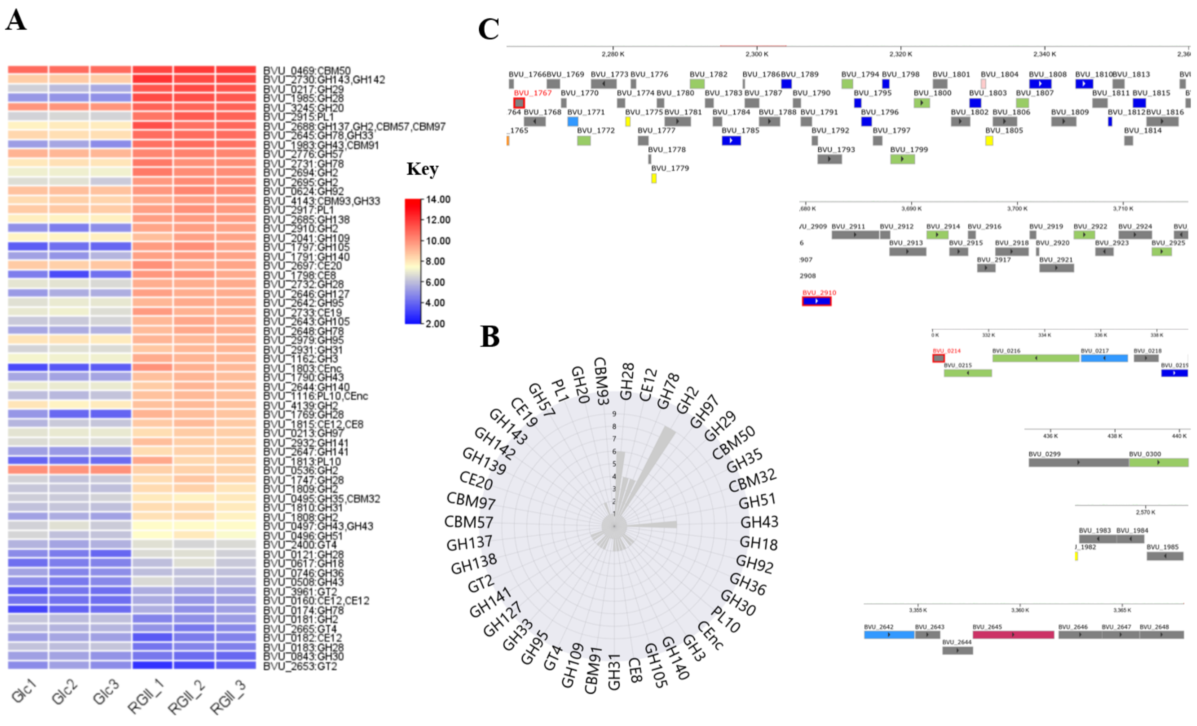


**Supplemental fig. 27: Differentially expressed CAZymes/CAZyme families during growth of *P. vulgatus* on RG-II and their genomic contexts. A:** Log2fold change of various CAZymes on RG-II relative to D-Glc*p* (Glc) from triplicate experiments (1-3). **B:** CAZyme count showing redundancy of various genes/families. **C:** Genomic context of various upregulated *P. vulgatus* RG-II-degrading CAZymes.

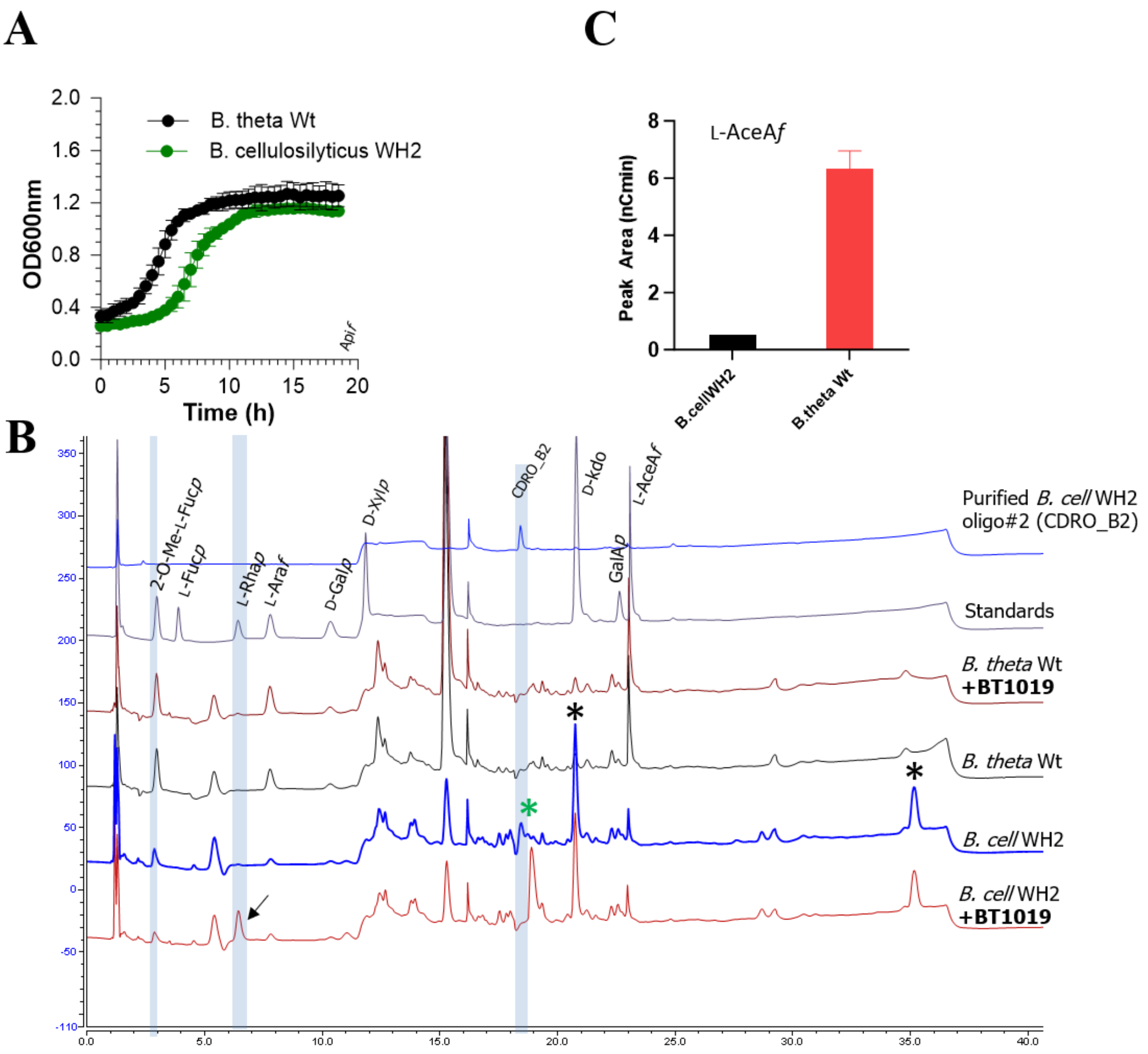


**Supplemental fig. 28: Growth and metabolism of RG-II by *B. cellulosilyticus* WH2. A:** Comparing growth of *B. cell* WH2 and *B. theta* Wt **B:** HPAEC-PAD analyses of culture supernatants following growth of *B. cell* WH2 on RG-II identifies unique sugars (starred peaks) generated by the strain including **CDRO_B2** (green starred peak) which was sensitive to the *B. theta*-derived L-rhamnosidase BT1019. Black arrow points to released L-Rha. CDRO_B2 was also successfully purified by SEC and confirmed by HPAEC-PAD (top chromatogram) **C**: Quantification of L-AceA*f* generated by *B. cell* WH2 and *B. theta* Wt.

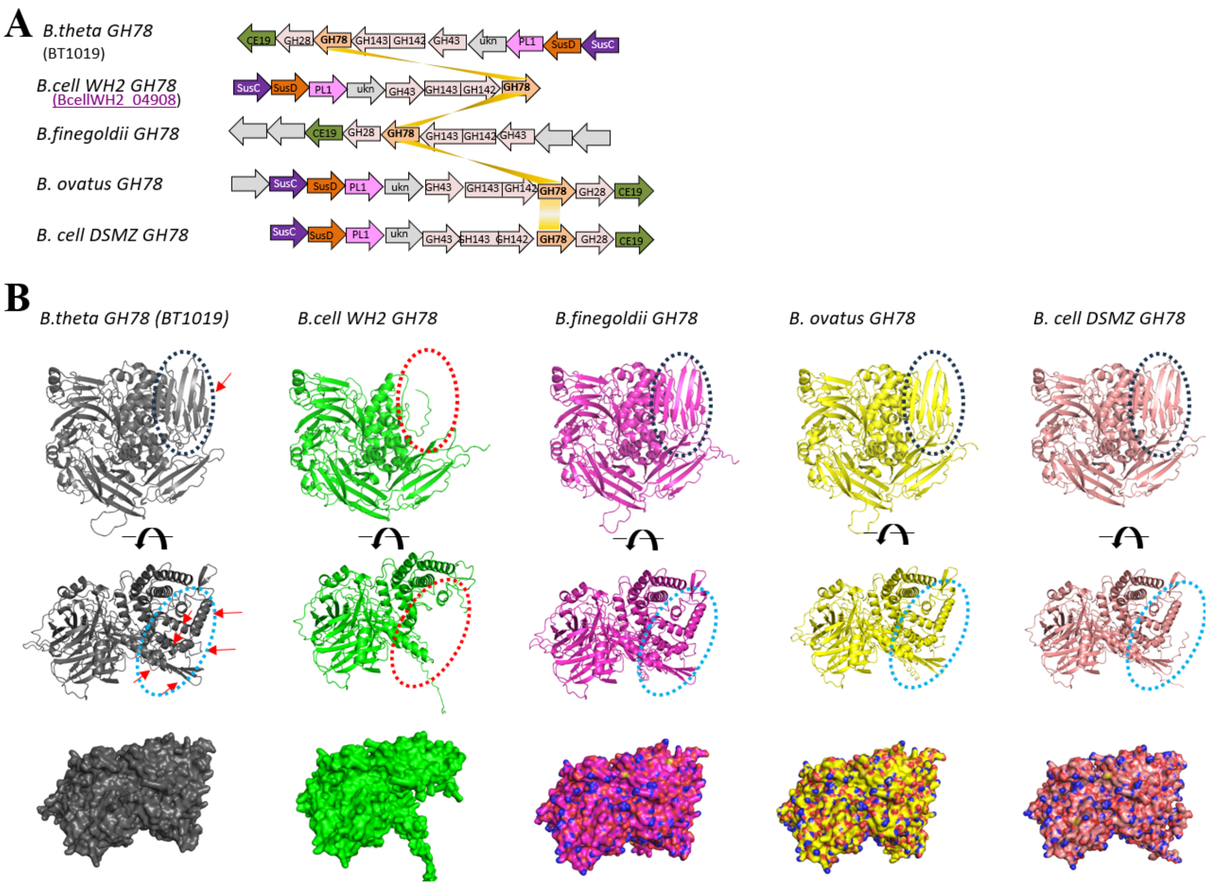


**Supplemental fig. 29: Comparing the genomic context and structure of *B. cell* WH2 GH78 enzyme. A:** Structure of various PULs carrying orthologues of *B. theta* GH78 enzymes including *B. cell* WH2 GH78. **B:** Comparison of AlphaFold structures of *B. theta* GH78 enzymes showing large portions of missing structural folds (red ellipses) in *B. cell* WH2 GH78 compared to the rest

**A**

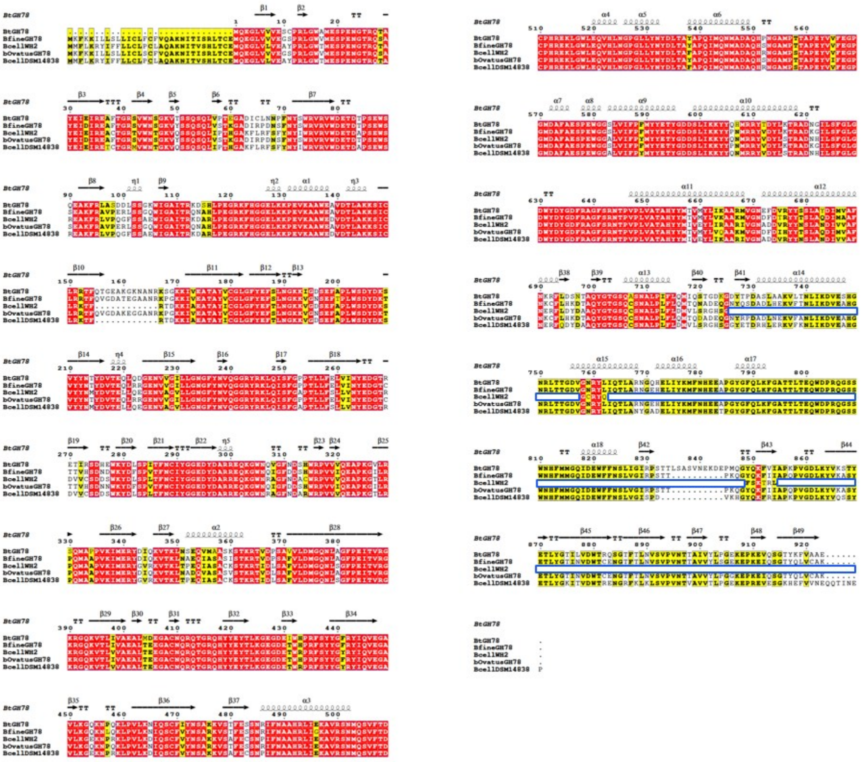

**B**

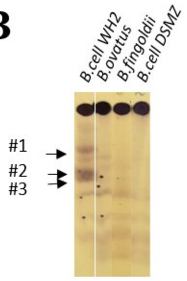


**Supplemental fig. 30:** Alignment of various *B. theta* GH78 orthologues including *B. cell* WH2 GH78. **(A)** and comparison of secretion product profiles post growth on RG-II by TLC **(B)**

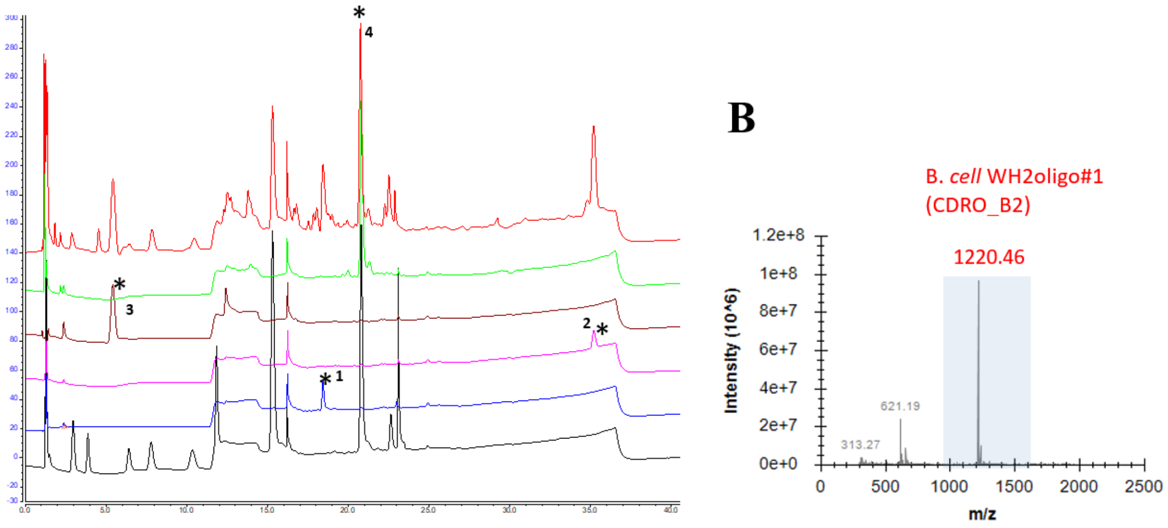


**Supplemental Fig. 31. Purification and analyses of sugars and CDROs generated by *B. cell* WH2.**
**A:** Various sugars including CDRO_B2 were resolved by SEC and analysed by HPAEC-PAD. Peaks for all purified sugars are starred **B**: MS analyses of purified CDRO_B2

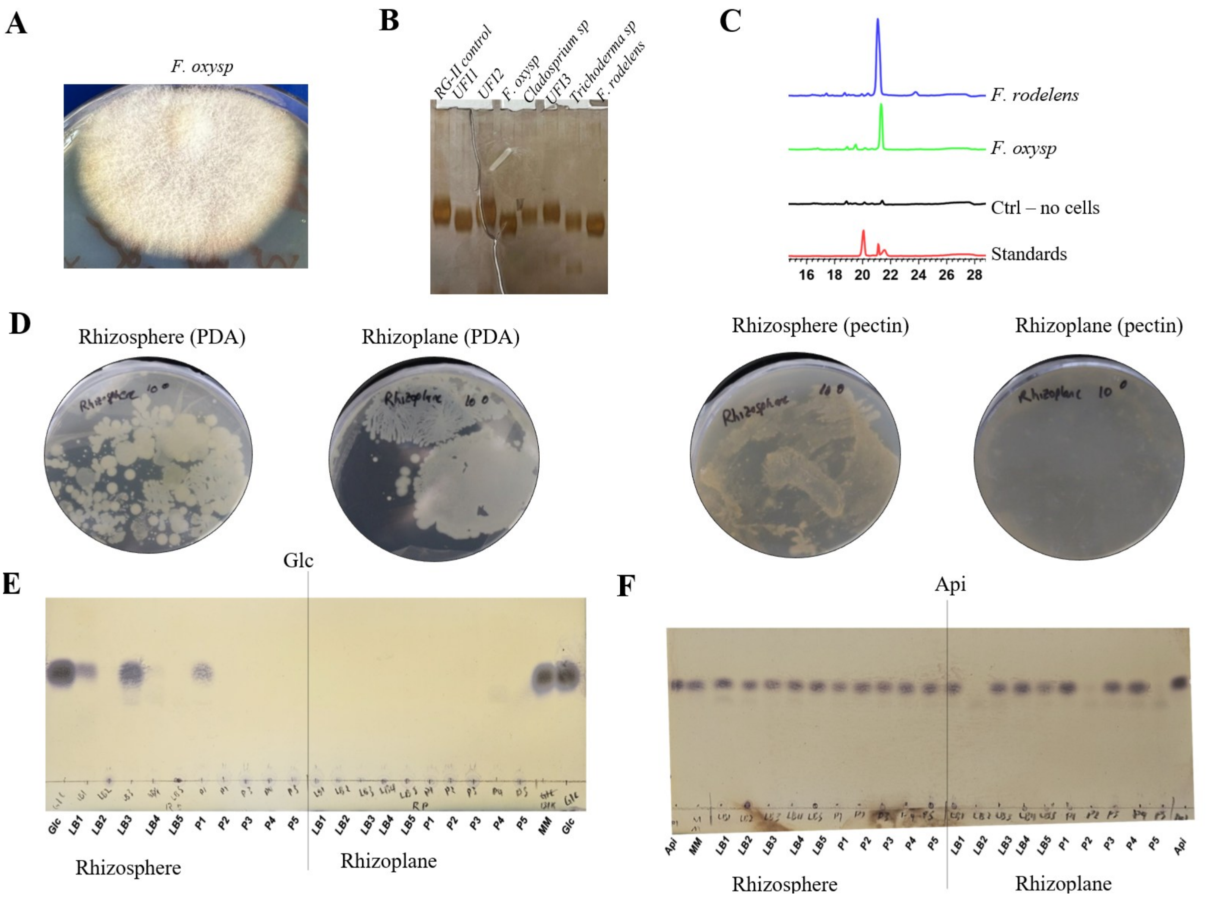


**Supplemental Fig. 32. Degradation of RG-II and utilisation of RG-II sugars by soil and plant associated microbes A:** Growth of a Barley (*Hordeum vulgare*) endophytic isolate of *F. oxysporum* on potatoe dextrose agar (PDA). **B:** Degradation of RG-II by various endophytic fungal root isolates from Barley (UFI denotes unknown fungal isolates) **C:** Detection of D-GalA*p* in culture during growth of *F. oxysporum* and *F. rodelens* on apple RG-II, **D:** Growth of rhizosphere and rhizoplane microbes from the roots of *N. benthamiana* on PDA and agar containing apple pectin **E:** Utilisation of D-Glc*p* (Glc) and D-Api*f* (Api) by diverse microbial strains from selected colonies in D. Colonies were cultured and then incubated in liquid medium containing D-Glc*p* or D-Api*f* overnight. Media were analysed by TLC for disappearance of D-Glc*p* and D-Api*f* as evidence for utilisation of the sugars. Majority of rhizosphere and rhizoplane sugars metabolised D-Glc*p,* however only three rhizoplane strains LB2, P2 and P5 metabolised D-Api*f*

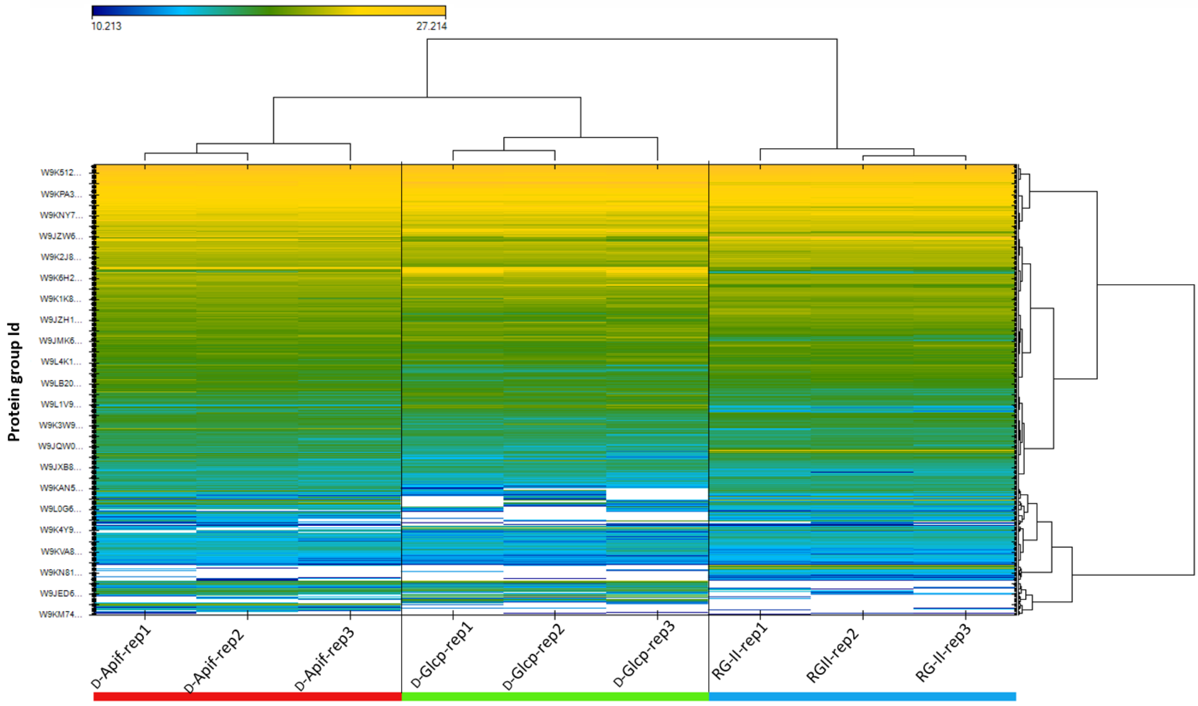


**Supplemental Fig. 33. Heatmap showing global differential gene expression in *F. oxysporum* root isolate (from Barley) during growth on three substrates D-Glc*p*, D-Api*f*, RG-II**

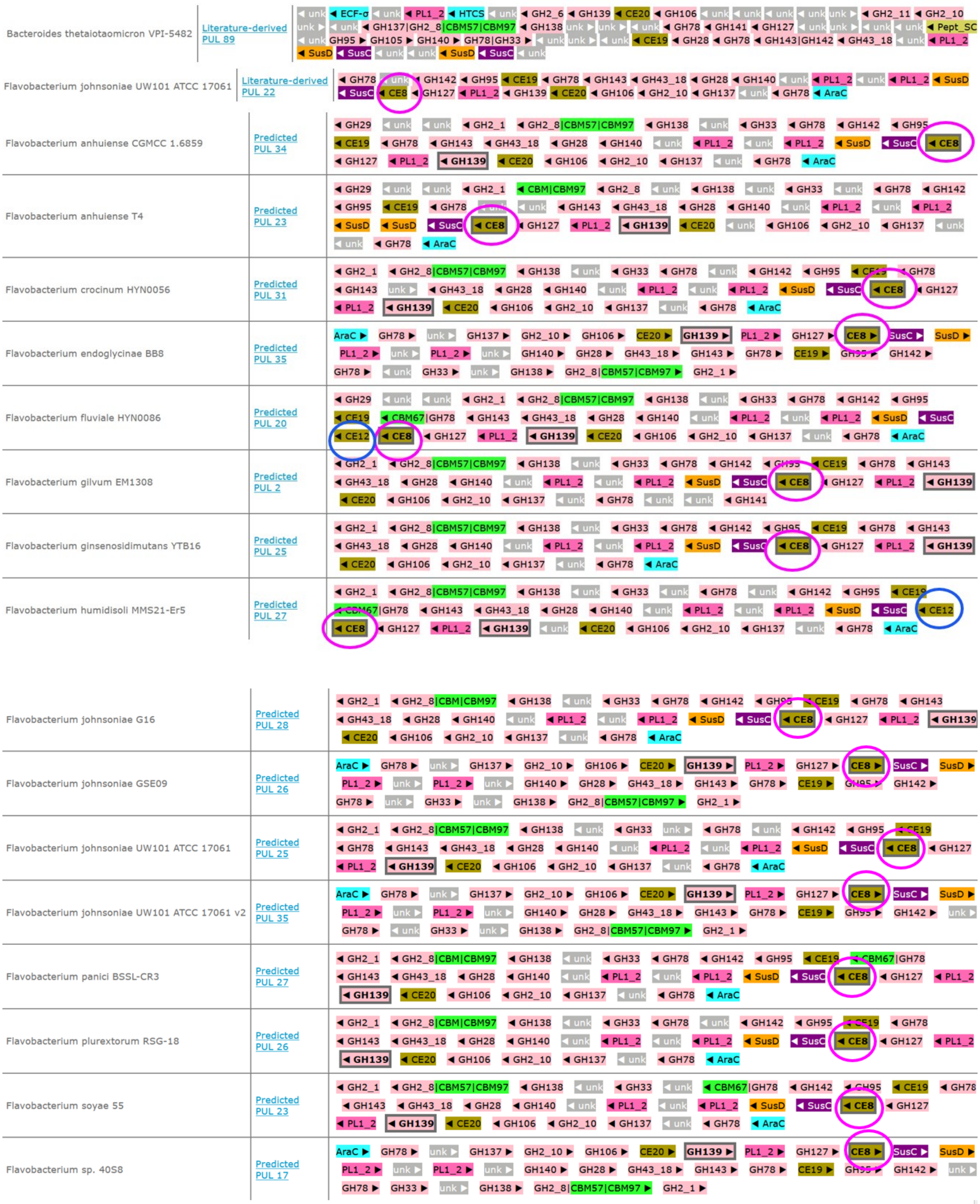

**Supplemental Fig. 34. Flavobacteria PULs (from PULDB database[7]) showing the presence of new features and CAZyme gene familiess e.g. AraC, CE8 and CE12 not normally associated with RG-II metabolism in *B. theta***

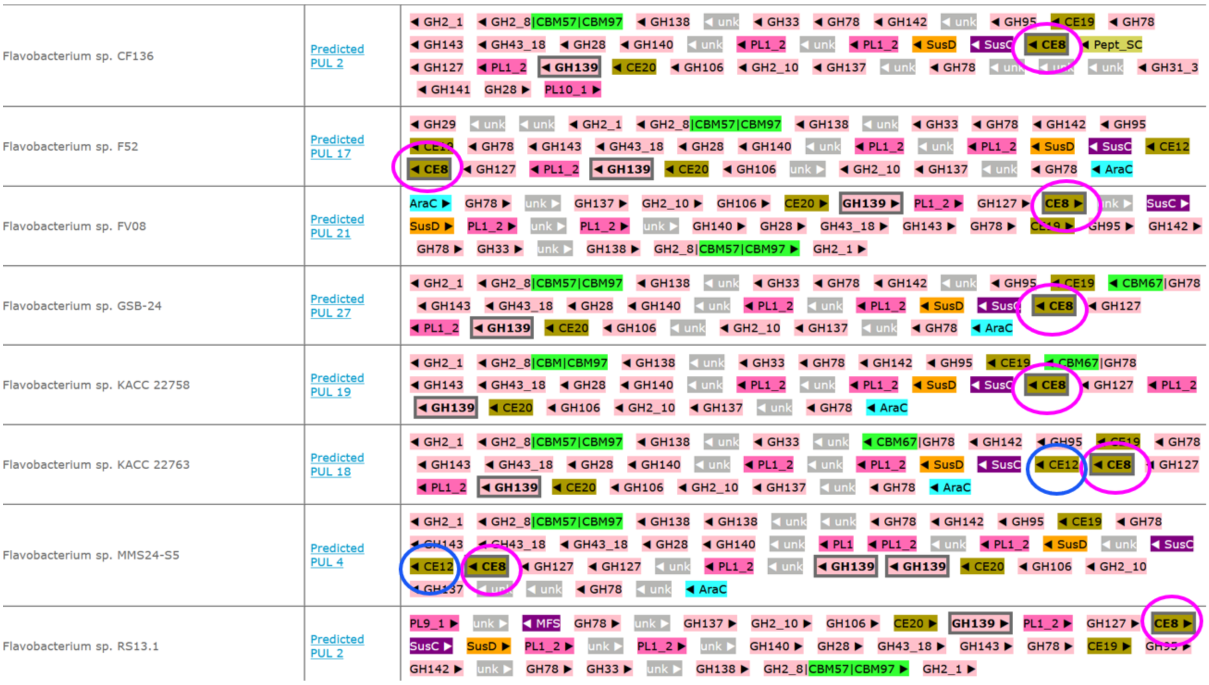

**Supplemental Fig. 34. Flavobacteria PULs (from PULDB database[7]) showing the presence of new features and CAZyme gene familiess e.g. AraC, CE8 and CE12 not normally associated with RG-II metabolism in *B. theta***

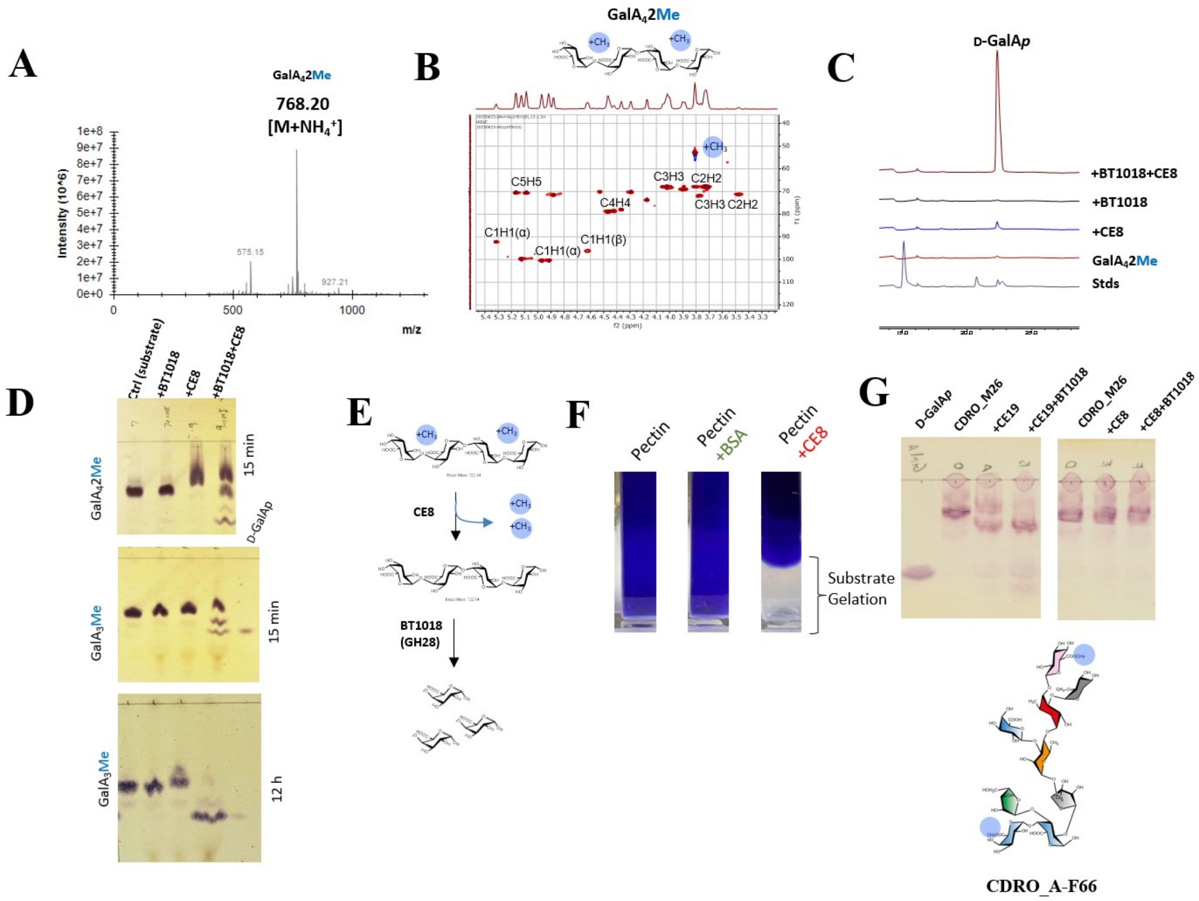


**Supplemental Fig. 35: Determining the biochemical activity Fjoh_4096 (CE8 enzyme-encoding gene) of *F. johnsoniae* UW101. A:** MS analyses of a purified substrate demethylated tetragalacturonate (GalA$_4$2Me). The substrate was prepared by digestion of apple pectin (75% esterification) followed by SEC purification and MS analyses **B**: NMR analyses of GalA$_4$2Me. **C:** HPAEC-PAD analyses showing coordination of Fjoh_4096 (CE8) and *B. theta*-derived polygalacturonase enzyme (BT1018) in the degradation of GalA$_4$2Me. Access to the substrate and release of D-GalA*p* is significantly increased following pretreatment of the substrate with Fjoh_4096 (CE8). D: TLC analyses showing concerted action of CE8 and BT1018 enzyme using GalA$_4$2Me and another methylated sugar (methyl-trigalacturonic acid) as substrates. E: Scheme of events for the concerted enzymatic activities of CE8 and BT1018. **F:** Results of CE8 catalysed pectin gelation using Fjoh_4096 as enzyme. **G:** TLC showing treatment of CDRO_A_F66 oligo with CE8 and CE19. Results show no processing of the sugar by CE8 as compared to the control enzyme CE19 known to degrade the substrate. Structure of the substrate is shown below the TLC plates.
