## Supplemental table 1 for "Towards a comprehensive chemical and genetic tool library for rhamnogalacturonan-II oligosaccharides and exploitation"

**Table 1. List of diverse chemically defined RG-II-derived oligosaccharide structures (CDROs) predicted and/or generated in this study and previous studies.** Underlined and yellow highlighted entries are some examples of CDROs that were conclusively produced or detected in this study. Starred CDROs are those shared structures between side chain A and B. The rest of the structures can potentially be generated based on the new findings in our current study.

**Side chain A/side chain A+backbone CDROs (CDRO_A#)**

**CDRO_A#**

1. L-Galp-α1,2-D-GlcA-β1,4(2-O-Me-D-Xylp-α1,3)-L-Fucp-α1,4-(D-GalAp-β1,3)(D-GalAp-α1,2)-L-Rhap-α1,-3-D-Apif
2. D-GlcA-β1,4(2-O-Me-D-Xylp-α1,3)-L-Fucp-α1,4-(D-GalAp-β1,3)(D-GalAp-α1,2)-L-Rhap-α1,3-D-Apif
3. 2-O-Me-D-Xylp-α1,3-L-Fucp-α1,4-(D-GalAp-β1,3)(D-GalAp-α1,2)-L-Rhap-α1,3-D-Apif
4. 2-O-Me-D-Xylp-α1,3-L-Fucp-α1,4-(D-GalAp-β1,3)-L-Rhap-α1,3-D-Apif
5. 2-O-Me-D-Xylp-α1,3-L-Fucp-α1,4-L-Rhap-α1,3-D-Apif
6. 2-O-Me-D-Xylp-α1,3-L-Fucp
7. L-Rhap-α1,3-D-Apif*
8. L-Galp-α1,2-D-GlcA-β1,4(2-O-Me-D-Xylp-α1,3)-L-Fucp-α1,4-((3-O-Me-)-D-GalAp-β1,3)(D-GalAp-α1,2)-L-Rhap-α1,3-D-Apif
9. D-GlcA-β1,4(2-O-Me-D-Xylp-α1,3)-L-Fucp-α1,4-((3-O-Me-)-D-GalAp-β1,3)(D-GalAp-α1,2)-L-Rhap-α1,3-D-Apif
10. 2-O-Me-D-Xylp-α1,3-L-Fucp-α1,4-((3-O-Me-)-D-GalAp-β1,3)(D-GalAp-α1,2)-L-Rhap-α1,3-D-Apif
11. 2-O-Me-D-Xylp-α1,3-L-Fucp-α1,4-((3-O-Me-)-D-GalAp-β1,3)-L-Rhap-α1,3-D-Apif
12. L-Galp-α1,2-D-GlcA-β1,4(2-O-Me-D-Xylp-α1,3)-L-Fucp-α1,4-((4-O-Me-)-D-GalAp-β1,3)(D-GalAp-α1,2)-L-Rhap-α1,3-D-Apif
13. D-GlcA-β1,4(2-O-Me-D-Xylp-α1,3)-L-Fucp-α1,4-((4-O-Me-)-D-GalAp-β1,3)(D-GalAp-α1,2)-L-Rhap-α1,3-D-Apif
14. 2-O-Me-D-Xylp-α1,3-L-Fucp-α1,4-((4-O-Me-)-D-GalAp-β1,3)(D-GalAp-α1,2)-L-Rhap-α1,3-D-Apif
15. 2-O-Me-D-Xylp-α1,3-L-Fucp-α1,4-((4-O-Me-)-D-GalAp-β1,3)-L-Rhap-α1,3-D-Apif
16. L-Galp-α1,2-D-GlcA-β1,4(2-O-Me-D-Xylp-α1,3)-L-Fucp-α1,4-((3-O-Me-)(4-O-Me-)-D-GalAp-β1,3)(D-GalAp-α1,2)-L-Rhap-α1,3-D-Apif
17. D-GlcA-β1,4(2-O-Me-D-Xylp-α1,3)-L-Fucp-α1,4-((3-O-Me-)(4-O-Me-)-D-GalAp-β1,3)(D-GalAp-α1,2)-L-Rhap-α1,3-D-Apif
18. 2-O-Me-D-Xylp-α1,3-L-Fucp-α1,4-((3-O-Me-)(4-O-Me-)-D-GalAp-β1,3)(D-GalAp-α1,2)-L-Rhap-α1,3-D-Apif
19. 2-O-Me-D-Xylp-α1,3-L-Fucp-α1,4-((3-O-Me-)(4-O-Me-)-D-GalAp-β1,3)-L-Rhap-α1,3-D-Apif
20. L-Galp-α1,2-D-GlcA-β1,4(2-O-Me-D-Xylp-α1,3)-L-Fucp-α1,4-(D-GalAp-β1,3)(D-GalAp-α1,2)-L-Rhap-α1,3-D-Apif-β1,2-D-GalAp
21. D-GlcA-β1,4(2-O-Me-D-Xylp-α1,3)-L-Fucp-α1,4-(D-GalAp-β1,3)(D-GalAp-α1,2)-L-Rhap-α1,3-D-Apif-β1,2-D-GalAp
22. 2-O-Me-D-Xylp-α1,3-L-Fucp-α1,4-(D-GalAp-β1,3)(D-GalAp-α1,2)-L-Rhap-α1,3-D-Apif-β1,2-D-GalAp
23. 2-O-Me-D-Xylp-α1,3-L-Fucp-α1,4-(D-GalAp-β1,3)-L-Rhap-α1,3-D-Apif-β1,2-D-GalAp
24. 2-O-Me-D-Xylp-α1,3-L-Fucp-α1,4-L-Rhap-α1,3-D-Apif-β1,2-D-GalAp
25. 2-O-Me-D-Xylp-α1,3-L-Fucp-β1,2-D-GalAp
26. L-Rhap-α1,3-D-Apif-β1,2-D-GalAp*
27. L-Galp-α1,2-D-GlcA-β1,4(2-O-Me-D-Xylp-α1,3)-L-Fucp-α1,4-((3-O-Me-)-D-GalAp-β1,3)(D-GalAp-α1,2)-L-Rhap-α1,3-D-Apif-β1,2-D-GalAp
28. D-GlcA-β1,4(2-O-Me-D-Xylp-α1,3)-L-Fucp-α1,4-((3-O-Me-)-D-GalAp-β1,3)(D-GalAp-α1,2)-L-Rhap-α1,3-D-Apif-β1,2-D-GalAp
29. 2-O-Me-D-Xylp-α1,3-L-Fucp-α1,4-((3-O-Me-)-D-GalAp-β1,3)(D-GalAp-α1,2)-L-Rhap-α1,3-D-Apif-β1,2-D-GalAp
30. 2-O-Me-D-Xylp-α1,3-L-Fucp-α1,4-((3-O-Me-)-D-GalAp-β1,3)-L-Rhap-α1,3-D-Apif-β1,2-D-GalAp
31. L-Galp-α1,2-D-GlcA-β1,4(2-O-Me-D-Xylp-α1,3)-L-Fucp-α1,4-((4-O-Me-)-D-GalAp-β1,3)(D-GalAp-α1,2)-L-Rhap-α1,3-D-Apif-β1,2-D-GalAp
32. D-GlcA-β1,4(2-O-Me-D-Xylp-α1,3)-L-Fucp-α1,4-((4-O-Me-)-D-GalAp-β1,3)(D-GalAp-α1,2)-L-Rhap-α1,3-D-Apif-β1,2-D-GalAp
33. 2-O-Me-D-Xylp-α1,3-L-Fucp-α1,4-((4-O-Me-)-D-GalAp-β1,3)(D-GalAp-α1,2)-L-Rhap-α1,3-D-Apif-β1,2-D-GalAp
34. 2-O-Me-D-Xylp-α1,3-L-Fucp-α1,4-((4-O-Me-)-D-GalAp-β1,3)-L-Rhap-α1,3-D-Apif-β1,2-D-GalAp
35. L-Galp-α1,2-D-GlcA-β1,4(2-O-Me-D-Xylp-α1,3)-L-Fucp-α1,4-((3-O-Me-)(4-O-Me-)-D-GalAp-β1,3)(D-GalAp-α1,2)-L-Rhap-α1,3-D-Apif-β1,2-D-GalAp
36. D-GlcA-β1,4(2-O-Me-D-Xylp-α1,3)-L-Fucp-α1,4-((3-O-Me-)(4-O-Me-)-D-GalAp-β1,3)(D-GalAp-α1,2)-L-Rhap-α1,3-D-Apif-β1,2-D-GalAp
37. 2-O-Me-D-Xylp-α1,3-L-Fucp-α1,4-((3-O-Me-)(4-O-Me-)-D-GalAp-β1,3)(D-GalAp-α1,2)-L-Rhap-α1,3-D-Apif-β1,2-D-GalAp
38. 2-O-Me-D-Xylp-α1,3-L-Fucp-α1,4-((3-O-Me-)(4-O-Me-)-D-GalAp-β1,3)-L-Rhap-α1,3-D-Apif-β1,2-D-GalAp

**Side chain A/side chain A+side chain F CDROs (CDRO_A-F#)**

**CDRO_A-F#**

39. L-Galp-α1,2-D-GlcA-β1,4(2-O-Me-D-Xylp-α1,3)-L-Fucp-α1,4-(D-GalAp-β1,3)(D-GalAp-α1,2)-L-Rhap-α1,3-D-Apif-β1,2-(L-Araf1,3-)-D-GalAp
40. D-GlcA-β1,4(2-O-Me-D-Xylp-α1,3)-L-Fucp-α1,4-(D-GalAp-β1,3)(D-GalAp-α1,2)-L-Rhap-α1,3-D-Apif-β1,2-(L-Araf1,3-)-D-GalAp
41. 2-O-Me-D-Xylp-α1,3-L-Fucp-α1,4-(D-GalAp-β1,3)(D-GalAp-α1,2)-L-Rhap-α1,3-D-Apif-β1,2-(L-Araf1,3-)-D-GalAp
42. 2-O-Me-D-Xylp-α1,3-L-Fucp-α1,4-(D-GalAp-β1,3)-L-Rhap-α1,3-D-Apif-β1,2-(L-Araf1,3-)-D-GalAp
43. 2-O-Me-D-Xylp-α1,3-L-Fucp-α1,4-L-Rhap-α1,3-D-Apif-β1,2-(L-Araf1,3-)-D-GalAp
44. 2-O-Me-D-Xylp-α1,3-L-Fucp-β1,2-(L-Araf1,3-)-D-GalAp
45. L-Rhap-α1,3-D-Apif-β1,2-(L-Araf1,3-)-D-GalAp*
46. L-Galp-α1,2-D-GlcA-β1,4(2-O-Me-D-Xylp-α1,3)-L-Fucp-α1,4-((3-O-Me-)-D-GalAp-β1,3)(D-GalAp-α1,2)-L-Rhap-α1,3-D-Apif-β1,2-(L-Araf1,3-)-D-GalAp
47. D-GlcA-β1,4(2-O-Me-D-Xylp-α1,3)-L-Fucp-α1,4-((3-O-Me-)-D-GalAp-β1,3)(D-GalAp-α1,2)-L-Rhap-α1,3-D-Apif-β1,2-(L-Araf1,3-)-D-GalAp
48. 2-O-Me-D-Xylp-α1,3-L-Fucp-α1,4-((3-O-Me-)-D-GalAp-β1,3)(D-GalAp-α1,2)-L-Rhap-α1,3-D-Apif-β1,2-(L-Araf1,3-)-D-GalAp
49. 2-O-Me-D-Xylp-α1,3-L-Fucp-α1,4-((3-O-Me-)-D-GalAp-β1,3)-L-Rhap-α1,3-D-Apif-β1,2-(L-Araf1,3-)-D-GalAp
50. L-Galp-α1,2-D-GlcA-β1,4(2-O-Me-D-Xylp-α1,3)-L-Fucp-α1,4-((4-O-Me-)-D-GalAp-β1,3)(D-GalAp-α1,2)-L-Rhap-α1,3-D-Apif-β1,2-(L-Araf1,3-)-D-GalAp
51. D-GlcA-β1,4(2-O-Me-D-Xylp-α1,3)-L-Fucp-α1,4-((4-O-Me-)-D-GalAp-β1,3)(D-GalAp-α1,2)-L-Rhap-α1,3-D-Apif-β1,2-(L-Araf1,3-)-D-GalAp
52. 2-O-Me-D-Xylp-α1,3-L-Fucp-α1,4-((4-O-Me-)-D-GalAp-β1,3)(D-GalAp-α1,2)-L-Rhap-α1,3-D-Apif-β1,2-(L-Araf1,3-)-D-GalAp
53. 2-O-Me-D-Xylp-α1,3-L-Fucp-α1,4-((4-O-Me-)-D-GalAp-β1,3)-L-Rhap-α1,3-D-Apif-β1,2-(L-Araf1,3-)-D-GalAp
54. L-Galp-α1,2-D-GlcA-β1,4(2-O-Me-D-Xylp-α1,3)-L-Fucp-α1,4-((3-O-Me-)(4-O-Me-)-D-GalAp-β1,3)(D-GalAp-α1,2)-L-Rhap-α1,3-D-Apif-β1,2-(L-Araf1,3-)-D-GalAp
55. D-GlcA-β1,4(2-O-Me-D-Xylp-α1,3)-L-Fucp-α1,4-((3-O-Me-)(4-O-Me-)-D-GalAp-β1,3)(D-GalAp-α1,2)-L-Rhap-α1,3-D-Apif-β1,2-(L-Araf1,3-)-D-GalAp
56. 2-O-Me-D-Xylp-α1,3-L-Fucp-α1,4-((3-O-Me-)(4-O-Me-)-D-GalAp-β1,3)(D-GalAp-α1,2)-L-Rhap-α1,3-D-Apif-β1,2-(L-Araf1,3-)-D-GalAp
57. 2-O-Me-D-Xylp-α1,3-L-Fucp-α1,4-((3-O-Me-)(4-O-Me-)-D-GalAp-β1,3)-L-Rhap-α1,3-D-Apif-β1,2-(L-Araf1,3-)-D-GalAp
58. L-Galp-α1,2-D-GlcA-β1,4(2-O-Me-D-Xylp-α1,3)-L-Fucp-α1,4-(D-GalAp-β1,3)(D-GalAp-α1,2)-L-Rhap-α1,3-D-Apif-β1,2-(6-O-Me-D-GalAp-α1,4-)(L-Araf1,3-)-D-GalAp
59. D-GlcA-β1,4(2-O-Me-D-Xylp-α1,3)-L-Fucp-α1,4-(D-GalAp-β1,3)(D-GalAp-α1,2)-L-Rhap-α1,3-D-Apif-β1,2-(6-O-Me-D-GalAp-α1,4-)(L-Araf1,3-)-D-GalAp
60. 2-O-Me-D-Xylp-α1,3-L-Fucp-α1,4-(D-GalAp-β1,3)(D-GalAp-α1,2)-L-Rhap-α1,3-D-Apif-β1,2-(6-O-Me-D-GalAp-α1,4-)(L-Araf1,3-)-D-GalAp
61. 2-O-Me-D-Xylp-α1,3-L-Fucp-α1,4-(D-GalAp-β1,3)-L-Rhap-α1,3-D-Apif-β1,2-(6-O-Me-D-GalAp-α1,4-)(L-Araf1,3-)-D-GalAp
62. 2-O-Me-D-Xylp-α1,3-L-Fucp-α1,4-L-Rhap-α1,3-D-Apif-β1,2-(6-O-Me-D-GalAp-α1,4-)(L-Araf1,3-)-D-GalAp
63. 2-O-Me-D-Xylp-α1,3-L-Fucp-β1,2-(6-O-Me-D-GalAp-α1,4-)(L-Araf1,3-)-D-GalAp
64. L-Rhap-α1,3-D-Apif-β1,2-(6-O-Me-D-GalAp-α1,4-)(L-Araf1,3-)-D-GalAp
65. L-Galp-α1,2-(6-O-Me-)-D-GlcA-β1,4(2-O-Me-D-Xylp-α1,3)-L-Fucp-α1,4-(D-GalAp-β1,3)(D-GalAp-α1,2)-L-Rhap-α1,3-D-Apif-β1,2-(6-O-Me-D-GalAp-α1,4-)(L-Araf1,3-)-D-GalAp
66. 6-O-Me-D-GlcA-β1,4(2-O-Me-D-Xylp-α1,3)-L-Fucp-α1,4-(D-GalAp-β1,3)(D-GalAp-α1,2)-L-Rhap-α1,3-D-Apif-β1,2-(6-O-Me-D-GalAp-α1,4-) (L-Araf1,3-)-D-GalAp
67. L-Galp-α1,2-D-GlcA-β1,4(2-O-Me-D-Xylp-α1,3)-L-Fucp-α1,4-((3-O-Me-)-D-GalAp-β1,3)(D-GalAp-α1,2)-L-Rhap-α1,3-D-Apif-β1,2-(6-O-Me-D-GalAp-α1,4-) (L-Araf1,3-)-D-GalAp
68. D-GlcA-β1,4(2-O-Me-D-Xylp-α1,3)-L-Fucp-α1,4-((3-O-Me-)-D-GalAp-β1,3)(D-GalAp-α1,2)-L-Rhap-α1,3-D-Apif-β1,2-(6-O-Me-D-GalAp-α1,4-) (L-Araf1,3-)-D-GalAp
69. 2-O-Me-D-Xylp-α1,3-L-Fucp-α1,4-((3-O-Me-)-D-GalAp-β1,3)(D-GalAp-α1,2)-L-Rhap-α1,3-D-Apif-β1,2-(6-O-Me-D-GalAp-α1,4-)(L-Araf1,3-)-D-GalAp
70. 2-O-Me-D-Xylp-α1,3-L-Fucp-α1,4-((3-O-Me-)-D-GalAp-β1,3)-L-Rhap-α1,3-D-Apif-β1,2-(6-O-Me-D-GalAp-α1,4-)(L-Araf1,3-)-D-GalAp
71. L-Galp-α1,2-D-GlcA-β1,4(2-O-Me-D-Xylp-α1,3)-L-Fucp-α1,4-((4-O-Me-)-D-GalAp-β1,3)(D-GalAp-α1,2)-L-Rhap-α1,3-D-Apif-β1,2-(6-O-Me-D-GalAp-α1,4-) (L-Araf1,3-)-D-GalAp
72. D-GlcA-β1,4(2-O-Me-D-Xylp-α1,3)-L-Fucp-α1,4-((4-O-Me-)-D-GalAp-β1,3)(D-GalAp-α1,2)-L-Rhap-α1,3-D-Apif-β1,2-(6-O-Me-D-GalAp-α1,4-) (L-Araf1,3-)-D-GalAp
73. 2-O-Me-D-Xylp-α1,3-L-Fucp-α1,4-((4-O-Me-)-D-GalAp-β1,3)(D-GalAp-α1,2)-L-Rhap-α1,3-D-Apif-β1,2-(6-O-Me-D-GalAp-α1,4-) (L-Araf1,3-)-D-GalAp
74. 2-O-Me-D-Xylp-α1,3-L-Fucp-α1,4-((4-O-Me-)-D-GalAp-β1,3)-L-Rhap-α1,3-D-Apif-β1,2-(6-O-Me-D-GalAp-α1,4-) (L-Araf1,3-)-D-GalAp
75. L-Galp-α1,2-D-GlcA-β1,4(2-O-Me-D-Xylp-α1,3)-L-Fucp-α1,4-((3-O-Me-)(4-O-Me-)-D-GalAp-β1,3)(D-GalAp-α1,2)-L-Rhap-α1,3-D-Apif-β1,2-(6-O-Me-D-GalAp-α1,4-)(L-Araf1,3-)-D-GalAp
76. D-GlcA-β1,4(2-O-Me-D-Xylp-α1,3)-L-Fucp-α1,4-((3-O-Me-)(4-O-Me-)-D-GalAp-β1,3)(D-GalAp-α1,2)-L-Rhap-α1,3-D-Apif-β1,2-(6-O-Me-D-GalAp-α1,4-)(L-Araf1,3-)-D-GalAp
77. 2-O-Me-D-Xylp-α1,3-L-Fucp-α1,4-((3-O-Me-)(4-O-Me-)-D-GalAp-β1,3)(D-GalAp-α1,2)-L-Rhap-α1,3-D-Apif-β1,2-(6-O-Me-D-GalAp-α1,4-) (L-Araf1,3-)-D-GalAp
78. 2-O-Me-D-Xylp-α1,3-L-Fucp-α1,4-((3-O-Me-)(4-O-Me-)-D-GalAp-β1,3)-L-Rhap-α1,3-D-Apif-β1,2-(6-O-Me-D-GalAp-α1,4-) (L-Araf1,3-)-D-GalAp
79. L-Galp-α1,2-(6-O-Me-)-D-GlcA-β1,4(2-O-Me-D-Xylp-α1,3)-L-Fucp-α1,4-((3-O-Me-)-D-GalAp-β1,3)(D-GalAp-α1,2)-L-Rhap-α1,3-D-Apif-β1,2-(6-O-Me-D-GalAp-α1,4-) (L-Araf1,3-)-D-GalAp
80. 6-O-Me-D-GlcA-β1,4(2-O-Me-D-Xylp-α1,3)-L-Fucp-α1,4-((3-O-Me-)-D-GalAp-β1,3)(D-GalAp-α1,2)-L-Rhap-α1,3-D-Apif-β1,2-(6-O-Me-D-GalAp-α1,4-) (L-Araf1,3-)-D-GalAp
81. L-Galp-α1,2-(6-O-Me-)-D-GlcA-β1,4(2-O-Me-D-Xylp-α1,3)-L-Fucp-α1,4-((4-O-Me-)-D-GalAp-β1,3)(D-GalAp-α1,2)-L-Rhap-α1,3-D-Apif-β1,2-(6-O-Me-D-GalAp-α1,4-) (L-Araf1,3-)-D-GalAp
82. 6-O-Me-D-GlcA-β1,4(2-O-Me-D-Xylp-α1,3)-L-Fucp-α1,4-((4-O-Me-)-D-GalAp-β1,3)(D-GalAp-α1,2)-L-Rhap-α1,3-D-Apif-β1,2-(6-O-Me-D-GalAp-α1,4-) (L-Araf1,3-)-D-GalAp
83. L-Galp-α1,2-(6-O-Me-)-D-GlcA-β1,4(2-O-Me-D-Xylp-α1,3)-L-Fucp-α1,4-((3-O-Me-)(4-O-Me-)-D-GalAp-β1,3)(D-GalAp-α1,2)-L-Rhap-α1,3-D-Apif-β1,2-(6-O-Me-D-GalAp-α1,4-) (L-Araf1,3-)-D-GalAp
84. 6-O-Me-D-GlcA-β1,4(2-O-Me-D-Xylp-α1,3)-L-Fucp-α1,4-((3-O-Me-)(4-O-Me-)-D-GalAp-β1,3)(D-GalAp-α1,2)-L-Rhap-α1,3-D-Apif-β1,2-(6-O-Me-D-GalAp-α1,4-) (L-Araf1,3-)-D-GalAp

**Side chain B/side chain B-backbone CDROs (CDRO_B#)**

1. L-Araf-β1,2-L-Rhap-α1,2-(L-Rhap-α-1,3-)-L-Arap-α1,4-(2-O-Me-L-Fucp-α1,2-)-D-Galp-β1,3-L-AceA-α1,3-L-Rhap-α1,3-D-Apif
2. L-Rhap-α1,2-(L-Rhap-α-1,3-)-L-Arap-α1,4-(2-O-Me-L-Fucp-α1,2-)-D-Galp-β1,3-L-AceA-α1,3-L-Rhap-α1,3-D-Apif
3. L-Rhap-α1,2-L-Arap-α1,4-(2-O-Me-L-Fucp-α1,2-)-D-Galp-β1,3-L-AceA-α1,3-L-Rhap-α1,3-D-Apif
4. L-Arap-α1,4-(2-O-Me-L-Fucp-α1,2-)-D-Galp-β1,3-L-AceA-α1,3-L-Rhap-α1,3-D-Apif
5. 2-O-Me-L-Fucp-α1,2-D-Galp-β1,3-L-AceA-α1,3-L-Rhap-α1,3-D-Apif
6. D-Galp-β1,3-L-AceA-α1,3-L-Rhap-α1,3-D-Apif
7. L-AceA-α1,3-L-Rhap-α1,3-D-Apif
8. L-Rhap-α1,3-D-Apif*
9. L-Araf-β1,2-L-Rhap-α1,2-(L-Rhap-α-1,3-)-L-Arap-α1,4-(2-O-Me-[3-OAc]-L-Fucp-α1,2-)-D-Galp-β1,3-L-AceA-α1,3-L-Rhap-α1,3-D-Apif
10. L-Rhap-α1,2-(L-Rhap-α-1,3-)-L-Arap-α1,4-(2-O-Me-[3-OAc]-L-Fucp-α1,2-)-D-Galp-β1,3-L-AceA-α1,3-L-Rhap-α1,3-D-Apif
11. L-Rhap-α1,2-L-Arap-α1,4-(2-O-Me-[3-OAc]-L-Fucp-α1,2-)-D-Galp-β1,3-L-AceA-α1,3-L-Rhap-α1,3-D-Apif
12. L-Arap-α1,4-(2-O-Me-[3-OAc]-L-Fucp-α1,2-)-D-Galp-β1,3-L-AceA-α1,3-L-Rhap-α1,3-D-Apif
13. 2-O-Me-[3-OAc]-L-Fucp-α1,2-D-Galp-β1,3-L-AceA-α1,3-L-Rhap-α1,3-D-Apif
14. L-Araf-β1,2-L-Rhap-α1,2-(L-Rhap-α-1,3-)-L-Arap-α1,4-(2-O-Me-L-Fucp-α1,2-)-D-Galp-β1,3-(4-OAc)-L-AceA-α1,3-L-Rhap-α1,3-D-Apif
15. L-Rhap-α1,2-(L-Rhap-α-1,3-)-L-Arap-α1,4-(2-O-Me-L-Fucp-α1,2-)-D-Galp-β1,3- (4-OAc)-L-AceA-α1,3-L-Rhap-α1,3-D-Apif
16. L-Rhap-α1,2-L-Arap-α1,4-(2-O-Me-L-Fucp-α1,2-)-D-Galp-β1,3- (4-OAc)-L-AceA-α1,3-L-Rhap-α1,3-D-Apif
17. L-Arap-α1,4-(2-O-Me-L-Fucp-α1,2-)-D-Galp-β1,3-(4-OAc)-L-AceA-α1,3-L-Rhap-α1,3-D-Apif

18. 2-O-Me-L-Fucp-α1,2-D-Galp-β1,3-(4-OAc)-L-AceA-α1,3-L-Rhap-α1,3-D-Apif
19. D-Galp-β1,3-(4-OAc)-L-AceA-α1,3-L-Rhap-α1,3-D-Apif
20. 4-OAc-L-AceA-α1,3-L-Rhap-α1,3-D-Apif

---

21. L-Araf-β1,2-L-Rhap-α1,2-(L-Rhap-α-1,3-)-L-Arap-α1,4-(2-O-Me-[3-OAc]-L-Fucp-α1,2-)-D-Galp-β1,3-(4-OAc)-L-AceA-α1,3-L-Rhap-α1,3-D-Apif
22. L-Rhap-α1,2-(L-Rhap-α-1,3-)-L-Arap-α1,4-(2-O-Me-[3-OAc]- L-Fucp-α1,2-)-D-Galp-β1,3- (4-OAc)-L-AceA-α1,3-L-Rhap-α1,3-D-Apif
23. L-Rhap-α1,2-L-Arap-α1,4-(2-O-Me-[3-OAc]- L-Fucp-α1,2-)-D-Galp-β1,3- (4-OAc)-L-AceA-α1,3-L-Rhap-α1,3-D-Apif
24. L-Arap-α1,4-(2-O-Me-[3-OAc]- L-Fucp-α1,2-)-D-Galp-β1,3-(4-OAc)-L-AceA-α1,3-L-Rhap-α1,3-D-Apif
25. 2-O-Me-[3-OAc]- L-Fucp-α1,2-D-Galp-β1,3-(4-OAc)-L-AceA-α1,3-L-Rhap-α1,3-D-Apif

---

26. L-Araf-β1,2-L-Rhap-α1,2-(L-Rhap-α-1,3-)-L-Arap-α1,4-(2-O-Me-L-Fucp-α1,2-)-D-Galp-β1,3-L-AceA-α1,3-L-Rhap-α1,3-D-Apif-β1,2-D-GalAp
27. L-Rhap-α1,2-(L-Rhap-α-1,3-)-L-Arap-α1,4-(2-O-Me-L-Fucp-α1,2-)-D-Galp-β1,3-L-AceA-α1,3-L-Rhap-α1,3-D-Apif-β1,2-D-GalAp
28. L-Rhap-α1,2-L-Arap-α1,4-(2-O-Me-L-Fucp-α1,2-)-D-Galp-β1,3-L-AceA-α1,3-L-Rhap-α1,3-D-Apif-β1,2-D-GalAp
29. L-Arap-α1,4-(2-O-Me-L-Fucp-α1,2-)-D-Galp-β1,3-L-AceA-α1,3-L-Rhap-α1,3-D-Apif-β1,2-D-GalAp
30. 2-O-Me-L-Fucp-α1,2-D-Galp-β1,3-L-AceA-α1,3-L-Rhap-α1,3-D-Apif-β1,2-D-GalAp*
31. D-Galp-β1,3-L-AceA-α1,3-L-Rhap-α1,3-D-Apif-β1,2-D-GalAp
32. L-AceA-α1,3-L-Rhap-α1,3-D-Apif-β1,2-D-GalAp
33. L-Rhap-α1,3-D-Apif-β1,2-D-GalAp*
34. L-Araf-β1,2-L-Rhap-α1,2-(L-Rhap-α-1,3-)-L-Arap-α1,4-(2-O-Me-[3-OAc]-L-Fucp-α1,2-)-D-Galp-β1,3-L-AceA-α1,3-L-Rhap-α1,3-D-Apif-β1,2-D-GalAp
35. L-Rhap-α1,2-(L-Rhap-α-1,3-)-L-Arap-α1,4-(2-O-Me-[3-OAc]-L-Fucp-α1,2-)-D-Galp-β1,3-L-AceA-α1,3-L-Rhap-α1,3-D-Apif-β1,2-D-GalAp
36. L-Rhap-α1,2-L-Arap-α1,4-(2-O-Me-[3-OAc]-L-Fucp-α1,2-)-D-Galp-β1,3-L-AceA-α1,3-L-Rhap-α1,3-D-Apif-β1,2-D-GalAp
37. L-Arap-α1,4-(2-O-Me-[3-OAc]-L-Fucp-α1,2-)-D-Galp-β1,3-L-AceA-α1,3-L-Rhap-α1,3-D-Apif-β1,2-D-GalAp
38. 2-O-Me-[3-OAc]-L-Fucp-α1,2-D-Galp-β1,3-L-AceA-α1,3-L-Rhap-α1,3-D-Apif-β1,2-D-GalAp
39. L-Araf-β1,2-L-Rhap-α1,2-(L-Rhap-α-1,3-)-L-Arap-α1,4-(2-O-Me-L-Fucp-α1,2-)-D-Galp-β1,3-(4-OAc)-L-AceA-α1,3-L-Rhap-α1,3-D-Apif-β1,2-D-GalAp
40. L-Rhap-α1,2-(L-Rhap-α-1,3-)-L-Arap-α1,4-(2-O-Me-L-Fucp-α1,2-)-D-Galp-β1,3- (4-OAc)-L-AceA-α1,3-L-Rhap-α1,3-D-Apif-β1,2-D-GalAp
41. L-Rhap-α1,2-L-Arap-α1,4-(2-O-Me-L-Fucp-α1,2-)-D-Galp-β1,3- (4-OAc)-L-AceA-α1,3-L-Rhap-α1,3-D-Apif-β1,2-D-GalAp
42. L-Arap-α1,4-(2-O-Me-L-Fucp-α1,2-)-D-Galp-β1,3-(4-OAc)-L-AceA-α1,3-L-Rhap-α1,3-D-Apif-β1,2-D-GalAp
43. 2-O-Me-L-Fucp-α1,2-D-Galp-β1,3-(4-OAc)-L-AceA-α1,3-L-Rhap-α1,3-D-Apif-β1,2-D-GalAp
44. D-Galp-β1,3-(4-OAc)-L-AceA-α1,3-L-Rhap-α1,3-D-Apif-β1,2-D-GalAp
45. 4-OAc-L-AceA-α1,3-L-Rhap-α1,3-D-Apif-β1,2-D-GalAp
46. L-Araf-β1,2-L-Rhap-α1,2-(L-Rhap-α-1,3-)-L-Arap-α1,4-(2-O-Me-[3-OAc]-L-Fucp-α1,2-)-D-Galp-β1,3-(4-OAc)-L-AceA-α1,3-L-Rhap-α1,3-D-Apif-β1,2-D-GalAp
47. L-Rhap-α1,2-(L-Rhap-α-1,3-)-L-Arap-α1,4-(2-O-Me-[3-OAc]- L-Fucp-α1,2-)-D-Galp-β1,3- (4-OAc)-L-AceA-α1,3-L-Rhap-α1,3-D-Apif-β1,2-D-GalAp
48. L-Rhap-α1,2-L-Arap-α1,4-(2-O-Me-[3-OAc]- L-Fucp-α1,2-)-D-Galp-β1,3- (4-OAc)-L-AceA-α1,3-L-Rhap-α1,3-D-Apif-β1,2-D-GalAp
49. L-Arap-α1,4-(2-O-Me-[3-OAc]- L-Fucp-α1,2-)-D-Galp-β1,3-(4-OAc)-L-AceA-α1,3-L-Rhap-α1,3-D-Apif-β1,2-D-GalAp
50. 2-O-Me-[3-OAc]- L-Fucp-α1,2-D-Galp-β1,3-(4-OAc)-L-AceA-α1,3-L-Rhap-α1,3-D-Apif-β1,2-D-GalAp
51. L-Araf-β1,2-L-Rhap-α1,2-(L-Rhap-α-1,3-)-L-Arap-α1,4-(2-O-Me-L-Fucp-α1,2-)-D-Galp-β1,3-L-AceA-α1,3-L-Rhap-α1,3-D-Apif-β1,2-[D-GalAp]$_2$
52. L-Rhap-α1,2-(L-Rhap-α-1,3-)-L-Arap-α1,4-(2-O-Me-L-Fucp-α1,2-)-D-Galp-β1,3-L-AceA-α1,3-L-Rhap-α1,3-D-Apif-β1,2-[D-GalAp]$_2$
53. L-Rhap-α1,2-L-Arap-α1,4-(2-O-Me-L-Fucp-α1,2-)-D-Galp-β1,3-L-AceA-α1,3-L-Rhap-α1,3-D-Apif-β1,2-[D-GalAp]$_2$
54. L-Arap-α1,4-(2-O-Me-L-Fucp-α1,2-)-D-Galp-β1,3-L-AceA-α1,3-L-Rhap-α1,3-D-Apif-β1,2-[D-GalAp]$_2$
55. 2-O-Me-L-Fucp-α1,2-D-Galp-β1,3-L-AceA-α1,3-L-Rhap-α1,3-D-Apif-β1,2-D-[D-GalAp]$_2$
56. D-Galp-β1,3-L-AceA-α1,3-L-Rhap-α1,3-D-Apif-β1,2-[D-GalAp]$_2$
57. L-AceA-α1,3-L-Rhap-α1,3-D-Apif-β1,2-[D-GalAp]$_2$
58. L-Rhap-α1,3-D-Apif-β1,2-D-[D-GalAp]$_2$
59. L-Araf-β1,2-L-Rhap-α1,2-(L-Rhap-α-1,3-)-L-Arap-α1,4-(2-O-Me-[3-OAc]-L-Fucp-α1,2-)-D-Galp-β1,3-L-AceA-α1,3-L-Rhap-α1,3-D-Apif-β1,2-[D-GalAp]$_2$
60. L-Rhap-α1,2-(L-Rhap-α-1,3-)-L-Arap-α1,4-(2-O-Me-[3-OAc]-L-Fucp-α1,2-)-D-Galp-β1,3-L-AceA-α1,3-L-Rhap-α1,3-D-Apif-β1,2-[D-GalAp]$_2$
61. L-Rhap-α1,2-L-Arap-α1,4-(2-O-Me-[3-OAc]-L-Fucp-α1,2-)-D-Galp-β1,3-L-AceA-α1,3-L-Rhap-α1,3-D-Apif-β1,2-[D-GalAp]$_2$
62. L-Arap-α1,4-(2-O-Me-[3-OAc]-L-Fucp-α1,2-)-D-Galp-β1,3-L-AceA-α1,3-L-Rhap-α1,3-D-Apif-β1,2-[D-GalAp]$_2$
63. 2-O-Me-[3-OAc]-L-Fucp-α1,2-D-Galp-β1,3-L-AceA-α1,3-L-Rhap-α1,3-D-Apif-β1,2-[D-GalAp]$_2$
64. L-Araf-β1,2-L-Rhap-α1,2-(L-Rhap-α-1,3-)-L-Arap-α1,4-(2-O-Me-L-Fucp-α1,2-)-D-Galp-β1,3-(4-OAc)-L-AceA-α1,3-L-Rhap-α1,3-D-Apif-β1,2-[D-GalAp]$_2$
65. L-Rhap-α1,2-(L-Rhap-α-1,3-)-L-Arap-α1,4-(2-O-Me-L-Fucp-α1,2-)-D-Galp-β1,3- (4-OAc)-L-AceA-α1,3-L-Rhap-α1,3-D-Apif-β1,2-[D-GalAp]$_2$
66. L-Rhap-α1,2-L-Arap-α1,4-(2-O-Me-L-Fucp-α1,2-)-D-Galp-β1,3- (4-OAc)-L-AceA-α1,3-L-Rhap-α1,3-D-Apif-β1,2-[D-GalAp]$_2$
67. L-Arap-α1,4-(2-O-Me-L-Fucp-α1,2-)-D-Galp-β1,3-(4-OAc)-L-AceA-α1,3-L-Rhap-α1,3-D-Apif-β1,2-[D-GalAp]$_2$
68. 2-O-Me-L-Fucp-α1,2-D-Galp-β1,3-(4-OAc)-L-AceA-α1,3-L-Rhap-α1,3-D-Apif-β1,2-[D-GalAp]$_2$
69. D-Galp-β1,3-(4-OAc)-L-AceA-α1,3-L-Rhap-α1,3-D-Apif-β1,2-[D-GalAp]$_2$
70. 4-OAc-L-AceA-α1,3-L-Rhap-α1,3-D-Apif-β1,2-[D-GalAp]$_2$
71. L-Araf-β1,2-L-Rhap-α1,2-(L-Rhap-α-1,3-)-L-Arap-α1,4-(2-O-Me-[3-OAc]-L-Fucp-α1,2-)-D-Galp-β1,3-(4-OAc)-L-AceA-α1,3-L-Rhap-α1,3-D-Apif-β1,2-[D-GalAp]$_2$
72. L-Rhap-α1,2-(L-Rhap-α-1,3-)-L-Arap-α1,4-(2-O-Me-[3-OAc]- L-Fucp-α1,2-)-D-Galp-β1,3- (4-OAc)-L-AceA-α1,3-L-Rhap-α1,3-D-Apif-β1,2-[D-GalAp]$_2$
73. L-Rhap-α1,2-L-Arap-α1,4-(2-O-Me-[3-OAc]- L-Fucp-α1,2-)-D-Galp-β1,3- (4-OAc)-L-AceA-α1,3-L-Rhap-α1,3-D-Apif-β1,2-[D-GalAp]$_2$
74. L-Arap-α1,4-(2-O-Me-[3-OAc]- L-Fucp-α1,2-)-D-Galp-β1,3-(4-OAc)-L-AceA-α1,3-L-Rhap-α1,3-D-Apif-β1,2-[D-GalAp]$_2$
75. 2-O-Me-[3-OAc]- L-Fucp-α1,2-D-Galp-β1,3-(4-OAc)-L-AceA-α1,3-L-Rhap-α1,3-D-Apif-β1,2-[D-GalAp]$_2$
76. L-Araf-β1,2-L-Rhap-α1,2-(L-Rhap-α-1,3-)-L-Arap-α1,4-(2-O-Me-L-Fucp-α1,2-)-D-Galp-β1,3-L-AceA-α1,3-L-Rhap-α1,3-D-Apif-β1,2-[D-GalAp]$_3$
77. L-Rhap-α1,2-(L-Rhap-α-1,3-)-L-Arap-α1,4-(2-O-Me-L-Fucp-α1,2-)-D-Galp-β1,3-L-AceA-α1,3-L-Rhap-α1,3-D-Apif-β1,2-[D-GalAp]$_3$
78. L-Rhap-α1,2-L-Arap-α1,4-(2-O-Me-L-Fucp-α1,2-)-D-Galp-β1,3-L-AceA-α1,3-L-Rhap-α1,3-D-Apif-β1,2-[D-GalAp]$_3$
79. L-Arap-α1,4-(2-O-Me-L-Fucp-α1,2-)-D-Galp-β1,3-L-AceA-α1,3-L-Rhap-α1,3-D-Apif-β1,2-[D-GalAp]$_3$
80. 2-O-Me-L-Fucp-α1,2-D-Galp-β1,3-L-AceA-α1,3-L-Rhap-α1,3-D-Apif-β1,2-D-[D-GalAp]$_3$
81. D-Galp-β1,3-L-AceA-α1,3-L-Rhap-α1,3-D-Apif-β1,2-[D-GalAp]$_3$
82. L-AceA-α1,3-L-Rhap-α1,3-D-Apif-β1,2-[D-GalAp]$_3$
83. L-Rhap-α1,3-D-Apif-β1,2-D-[D-GalAp]$_3$
84. L-Araf-β1,2-L-Rhap-α1,2-(L-Rhap-α-1,3-)-L-Arap-α1,4-(2-O-Me-[3-OAc]-L-Fucp-α1,2-)-D-Galp-β1,3-L-AceA-α1,3-L-Rhap-α1,3-D-Apif-β1,2-[D-GalAp]$_3$
85. L-Rhap-α1,2-(L-Rhap-α-1,3-)-L-Arap-α1,4-(2-O-Me-[3-OAc]-L-Fucp-α1,2-)-D-Galp-β1,3-L-AceA-α1,3-L-Rhap-α1,3-D-Apif-β1,2-[D-GalAp]$_3$
86. L-Rhap-α1,2-L-Arap-α1,4-(2-O-Me-[3-OAc]-L-Fucp-α1,2-)-D-Galp-β1,3-L-AceA-α1,3-L-Rhap-α1,3-D-Apif-β1,2-[D-GalAp]$_3$
87. L-Arap-α1,4-(2-O-Me-[3-OAc]-L-Fucp-α1,2-)-D-Galp-β1,3-L-AceA-α1,3-L-Rhap-α1,3-D-Apif-β1,2-[D-GalAp]$_3$
88. 2-O-Me-[3-OAc]-L-Fucp-α1,2-D-Galp-β1,3-L-AceA-α1,3-L-Rhap-α1,3-D-Apif-β1,2-[D-GalAp]$_3$
89. L-Araf-β1,2-L-Rhap-α1,2-(L-Rhap-α-1,3-)-L-Arap-α1,4-(2-O-Me-L-Fucp-α1,2-)-D-Galp-β1,3-(4-OAc)-L-AceA-α1,3-L-Rhap-α1,3-D-Apif-β1,2-[D-GalAp]$_3$
90. L-Rhap-α1,2-(L-Rhap-α-1,3-)-L-Arap-α1,4-(2-O-Me-L-Fucp-α1,2-)-D-Galp-β1,3- (4-OAc)-L-AceA-α1,3-L-Rhap-α1,3-D-Apif-β1,2-[D-GalAp]$_3$
91. L-Rhap-α1,2-L-Arap-α1,4-(2-O-Me-L-Fucp-α1,2-)-D-Galp-β1,3- (4-OAc)-L-AceA-α1,3-L-Rhap-α1,3-D-Apif-β1,2-[D-GalAp]$_3$
92. L-Arap-α1,4-(2-O-Me-L-Fucp-α1,2-)-D-Galp-β1,3-(4-OAc)-L-AceA-α1,3-L-Rhap-α1,3-D-Apif-β1,2-[D-GalAp]$_3$
93. 2-O-Me-L-Fucp-α1,2-D-Galp-β1,3-(4-OAc)-L-AceA-α1,3-L-Rhap-α1,3-D-Apif-β1,2-[D-GalAp]$_3$
94. D-Galp-β1,3-(4-OAc)-L-AceA-α1,3-L-Rhap-α1,3-D-Apif-β1,2-[D-GalAp]$_3$
95. 4-OAc-L-AceA-α1,3-L-Rhap-α1,3-D-Apif-β1,2-[D-GalAp]$_3$
96. L-Araf-β1,2-L-Rhap-α1,2-(L-Rhap-α-1,3-)-L-Arap-α1,4-(2-O-Me-[3-OAc]-L-Fucp-α1,2-)-D-Galp-β1,3-(4-OAc)-L-AceA-α1,3-L-Rhap-α1,3-D-Apif-β1,2-[D-GalAp]$_3$
97. L-Rhap-α1,2-(L-Rhap-α-1,3-)-L-Arap-α1,4-(2-O-Me-[3-OAc]- L-Fucp-α1,2-)-D-Galp-β1,3- (4-OAc)-L-AceA-α1,3-L-Rhap-α1,3-D-Apif-β1,2-[D-GalAp]$_3$
98. L-Rhap-α1,2-L-Arap-α1,4-(2-O-Me-[3-OAc]- L-Fucp-α1,2-)-D-Galp-β1,3- (4-OAc)-L-AceA-α1,3-L-Rhap-α1,3-D-Apif-β1,2-[D-GalAp]$_3$
99. L-Arap-α1,4-(2-O-Me-[3-OAc]- L-Fucp-α1,2-)-D-Galp-β1,3-(4-OAc)-L-AceA-α1,3-L-Rhap-α1,3-D-Apif-β1,2-[D-GalAp]$_3$
100. 2-O-Me-[3-OAc]- L-Fucp-α1,2-D-Galp-β1,3-(4-OAc)-L-AceA-α1,3-L-Rhap-α1,3-D-Apif-β1,2-[D-GalAp]$_3$
101. L-Araf-β1,2-L-Rhap-α1,2-(L-Rhap-α-1,3-)-L-Arap-α1,4-(2-O-Me-L-Fucp-α1,2-)-D-Galp-β1,3-L-AceA-α1,3-L-Rhap-α1,3-D-Apif-β1,2-[D-GalAp]$_4$
102. L-Rhap-α1,2-(L-Rhap-α-1,3-)-L-Arap-α1,4-(2-O-Me-L-Fucp-α1,2-)-D-Galp-β1,3-L-AceA-α1,3-L-Rhap-α1,3-D-Apif-β1,2-[D-GalAp]$_4$
103. L-Rhap-α1,2-L-Arap-α1,4-(2-O-Me-L-Fucp-α1,2-)-D-Galp-β1,3-L-AceA-α1,3-L-Rhap-α1,3-D-Apif-β1,2-[D-GalAp]$_4$
104. L-Arap-α1,4-(2-O-Me-L-Fucp-α1,2-)-D-Galp-β1,3-L-AceA-α1,3-L-Rhap-α1,3-D-Apif-β1,2-[D-GalAp]$_4$
105. 2-O-Me-L-Fucp-α1,2-D-Galp-β1,3-L-AceA-α1,3-L-Rhap-α1,3-D-Apif-β1,2-D-[D-GalAp]$_4$
106. D-Galp-β1,3-L-AceA-α1,3-L-Rhap-α1,3-D-Apif-β1,2-[D-GalAp]$_4$
107. L-AceA-α1,3-L-Rhap-α1,3-D-Apif-β1,2-[D-GalAp]$_4$
108. L-Rhap-α1,3-D-Apif-β1,2-D-[D-GalAp]$_4$
109. L-Araf-β1,2-L-Rhap-α1,2-(L-Rhap-α-1,3-)-L-Arap-α1,4-(2-O-Me-[3-OAc]-L-Fucp-α1,2-)-D-Galp-β1,3-L-AceA-α1,3-L-Rhap-α1,3-D-Apif-β1,2-[D-GalAp]$_4$
110. L-Rhap-α1,2-(L-Rhap-α-1,3-)-L-Arap-α1,4-(2-O-Me-[3-OAc]-L-Fucp-α1,2-)-D-Galp-β1,3-L-AceA-α1,3-L-Rhap-α1,3-D-Apif-β1,2-[D-GalAp]$_4$
111. L-Rhap-α1,2-L-Arap-α1,4-(2-O-Me-[3-OAc]-L-Fucp-α1,2-)-D-Galp-β1,3-L-AceA-α1,3-L-Rhap-α1,3-D-Apif-β1,2-[D-GalAp]$_4$
112. L-Arap-α1,4-(2-O-Me-[3-OAc]-L-Fucp-α1,2-)-D-Galp-β1,3-L-AceA-α1,3-L-Rhap-α1,3-D-Apif-β1,2-[D-GalAp]$_4$
113. 2-O-Me-[3-OAc]-L-Fucp-α1,2-D-Galp-β1,3-L-AceA-α1,3-L-Rhap-α1,3-D-Apif-β1,2-[D-GalAp]$_4$
114. L-Araf-β1,2-L-Rhap-α1,2-(L-Rhap-α-1,3-)-L-Arap-α1,4-(2-O-Me-L-Fucp-α1,2-)-D-Galp-β1,3-(4-OAc)-L-AceA-α1,3-L-Rhap-α1,3-D-Apif-β1,2-[D-GalAp]$_4$
115. L-Rhap-α1,2-(L-Rhap-α-1,3-)-L-Arap-α1,4-(2-O-Me-L-Fucp-α1,2-)-D-Galp-β1,3- (4-OAc)-L-AceA-α1,3-L-Rhap-α1,3-D-Apif-β1,2-[D-GalAp]$_4$
116. L-Rhap-α1,2-L-Arap-α1,4-(2-O-Me-L-Fucp-α1,2-)-D-Galp-β1,3- (4-OAc)-L-AceA-α1,3-L-Rhap-α1,3-D-Apif-β1,2-[D-GalAp]$_4$
117. L-Arap-α1,4-(2-O-Me-L-Fucp-α1,2-)-D-Galp-β1,3-(4-OAc)-L-AceA-α1,3-L-Rhap-α1,3-D-Apif-β1,2-[D-GalAp]$_4$
118. 2-O-Me-L-Fucp-α1,2-D-Galp-β1,3-(4-OAc)-L-AceA-α1,3-L-Rhap-α1,3-D-Apif-β1,2-[D-GalAp]$_4$
119. D-Galp-β1,3-(4-OAc)-L-AceA-α1,3-L-Rhap-α1,3-D-Apif-β1,2-[D-GalAp]$_4$
120. 4-OAc-L-AceA-α1,3-L-Rhap-α1,3-D-Apif-β1,2-[D-GalAp]$_4$
121. L-Araf-β1,2-L-Rhap-α1,2-(L-Rhap-α-1,3-)-L-Arap-α1,4-(2-O-Me-[3-OAc]-L-Fucp-α1,2-)-D-Galp-β1,3-(4-OAc)-L-AceA-α1,3-L-Rhap-α1,3-D-Apif-β1,2-[D-GalAp]$_4$
122. L-Rhap-α1,2-(L-Rhap-α-1,3-)-L-Arap-α1,4-(2-O-Me-[3-OAc]- L-Fucp-α1,2-)-D-Galp-β1,3- (4-OAc)-L-AceA-α1,3-L-Rhap-α1,3-D-Apif-β1,2-[D-GalAp]$_4$
123. L-Rhap-α1,2-L-Arap-α1,4-(2-O-Me-[3-OAc]- L-Fucp-α1,2-)-D-Galp-β1,3- (4-OAc)-L-AceA-α1,3-L-Rhap-α1,3-D-Apif-β1,2-[D-GalAp]$_4$
124. L-Arap-α1,4-(2-O-Me-[3-OAc]- L-Fucp-α1,2-)-D-Galp-β1,3-(4-OAc)-L-AceA-α1,3-L-Rhap-α1,3-D-Apif-β1,2-[D-GalAp]$_4$
125. 2-O-Me-[3-OAc]- L-Fucp-α1,2-D-Galp-β1,3-(4-OAc)-L-AceA-α1,3-L-Rhap-α1,3-D-Apif-β1,2-[D-GalAp]$_4$

**D/B-backbone CDROs (CDRO-D/B#)**

**1.** L-Araf-β1,2-L-Rhap-α1,2-(L-Rhap-α-1,3-)-L-Arap-α1,4-(2-O-Me-L-Fucp-α1,2-)-D-Galp-β1,3-L-AceA-α1,3-L-Rhap-α1,3-D-Apif-β1,2-(L-Araf-β1,5-D-Dha-β1,3-)[UA/D-GalAp]n
**2.** L-Rhap-α1,2-(L-Rhap-α-1,3-)-L-Arap-α1,4-(2-O-Me-L-Fucp-α1,2-)-D-Galp-β1,3-L-AceA-α1,3-L-Rhap-α1,3-D-Apif-β1,2-(L-Araf-β1,5-D-Dha-β1,3-)[UA/D-GalAp]n
**3.** L-Rhap-α1,2-L-Arap-α1,4-(2-O-Me-L-Fucp-α1,2-)-D-Galp-β1,3-L-AceA-α1,3-L-Rhap-α1,3-D-Apif-β1,2-(L-Araf-β1,5-D-Dha-β1,3-)[UA/D-GalAp]n
**4.** L-Arap-α1,4-(2-O-Me-L-Fucp-α1,2-)-D-Galp-β1,3-L-AceA-α1,3-L-Rhap-α1,3-D-Apif-β1,2-(L-Araf-β1,5-D-Dha-β1,3-)[UA/D-GalAp]n
**5.** 2-O-Me-L-Fucp-α1,2-D-Galp-β1,3-L-AceA-α1,3-L-Rhap-α1,3-D-Apif-β1,2-D-(L-Araf-β1,5-D-Dha-β1,3-)[UA/D-GalAp]n
**6.** D-Galp-β1,3-L-AceA-α1,3-L-Rhap-α1,3-D-Apif-β1,2-(L-Araf-β1,5-D-Dha-β1,3-)[UA/D-GalAp]n
**7.** L-AceA-α1,3-L-Rhap-α1,3-D-Apif-β1,2-(L-Araf-β1,5-D-Dha-β1,3-)[UA/D-GalAp]n
**8.** L-Rhap-α1,3-D-Apif-β1,2-D-(L-Araf-β1,5-D-Dha-β1,3-)[UA/D-GalAp]n
**9.** L-Araf-β1,2-L-Rhap-α1,2-(L-Rhap-α-1,3-)-L-Arap-α1,4-(2-O-Me-[3-OAc]-L-Fucp-α1,2-)-D-Galp-β1,3-L-AceA-α1,3-L-Rhap-α1,3-D-Apif-β1,2-(L-Araf-β1,5-D-Dha-β1,3-)[UA/D-GalAp]n
**10.** L-Rhap-α1,2-(L-Rhap-α-1,3-)-L-Arap-α1,4-(2-O-Me-[3-OAc]-L-Fucp-α1,2-)-D-Galp-β1,3-L-AceA-α1,3-L-Rhap-α1,3-D-Apif-β1,2-(L-Araf-β1,5-D-Dha-β1,3-)[UA/D-GalAp]n
**11.** L-Rhap-α1,2-L-Arap-α1,4-(2-O-Me-[3-OAc]-L-Fucp-α1,2-)-D-Galp-β1,3-L-AceA-α1,3-L-Rhap-α1,3-D-Apif-β1,2-(L-Araf-β1,5-D-Dha-β1,3-)[UA/D-GalAp]n
**12.** L-Arap-α1,4-(2-O-Me-[3-OAc]-L-Fucp-α1,2-)-D-Galp-β1,3-L-AceA-α1,3-L-Rhap-α1,3-D-Apif-β1,2-(L-Araf-β1,5-D-Dha-β1,3-)[UA/D-GalAp]n
**13.** 2-O-Me-[3-OAc]-L-Fucp-α1,2-D-Galp-β1,3-L-AceA-α1,3-L-Rhap-α1,3-D-Apif-β1,2-(L-Araf-β1,5-D-Dha-β1,3-)[UA/D-GalAp]n
**14.** L-Araf-β1,2-L-Rhap-α1,2-(L-Rhap-α-1,3-)-L-Arap-α1,4-(2-O-Me-L-Fucp-α1,2-)-D-Galp-β1,3-(4-OAc)-L-AceA-α1,3-L-Rhap-α1,3-D-Apif-β1,2-(L-Araf-β1,5-D-Dha-β1,3-)[ UA/D-GalAp]n
**15.** L-Rhap-α1,2-(L-Rhap-α-1,3-)-L-Arap-α1,4-(2-O-Me-L-Fucp-α1,2-)-D-Galp-β1,3- (4-OAc)-L-AceA-α1,3-L-Rhap-α1,3-D-Apif-β1,2-(L-Araf-β1,5-D-Dha-β1,3-)[ UA/D-GalAp]n
**16.** L-Rhap-α1,2-L-Arap-α1,4-(2-O-Me-L-Fucp-α1,2-)-D-Galp-β1,3- (4-OAc)-L-AceA-α1,3-L-Rhap-α1,3-D-Apif-β1,2-(L-Araf-β1,5-D-Dha-β1,3-)[ UA/D-GalAp]n
**17.** L-Arap-α1,4-(2-O-Me-L-Fucp-α1,2-)-D-Galp-β1,3-(4-OAc)-L-AceA-α1,3-L-Rhap-α1,3-D-Apif-β1,2-(L-Araf-β1,5-D-Dha-β1,3-)[ UA/D-GalAp]n
**18.** 2-O-Me-L-Fucp-α1,2-D-Galp-β1,3-(4-OAc)-L-AceA-α1,3-L-Rhap-α1,3-D-Apif-β1,2-(L-Araf-β1,5-D-Dha-β1,3-)[ UA/D-GalAp]n
**19.** D-Galp-β1,3-(4-OAc)-L-AceA-α1,3-L-Rhap-α1,3-D-Apif-β1,2-(L-Araf-β1,5-D-Dha-β1,3-)[ UA/D-GalAp]n
**20.** 4-OAc-L-AceA-α1,3-L-Rhap-α1,3-D-Apif-β1,2-(L-Araf-β1,5-D-Dha-β1,3-)[ UA/D-GalAp]n
**21.** L-Araf-β1,2-L-Rhap-α1,2-(L-Rhap-α-1,3-)-L-Arap-α1,4-(2-O-Me-[3-OAc]-L-Fucp-α1,2-)-D-Galp-β1,3-(4-OAc)-L-AceA-α1,3-L-Rhap-α1,3-D-Apif-β1,2-(L-Araf-β1,5-D-Dha-β1,3-)[ UA/D-GalAp]n
**22.** L-Rhap-α1,2-(L-Rhap-α-1,3-)-L-Arap-α1,4-(2-O-Me-[3-OAc]- L-Fucp-α1,2-)-D-Galp-β1,3- (4-OAc)-L-AceA-α1,3-L-Rhap-α1,3-D-Apif-β1,2-(L-Araf-β1,5-D-Dha-β1,3-)[ UA/D-GalAp]n
**23.** L-Rhap-α1,2-L-Arap-α1,4-(2-O-Me-[3-OAc]- L-Fucp-α1,2-)-D-Galp-β1,3- (4-OAc)-L-AceA-α1,3-L-Rhap-α1,3-D-Apif-β1,2-(L-Araf-β1,5-D-Dha-β1,3-)[ UA/D-GalAp]n
**24.** L-Arap-α1,4-(2-O-Me-[3-OAc]- L-Fucp-α1,2-)-D-Galp-β1,3-(4-OAc)-L-AceA-α1,3-L-Rhap-α1,3-D-Apif-β1,2-(L-Araf-β1,5-D-Dha-β1,3-)[ UA/D-GalAp]n
**25.** 2-O-Me-[3-OAc]- L-Fucp-α1,2-D-Galp-β1,3-(4-OAc)-L-AceA-α1,3-L-Rhap-α1,3-D-Apif-β1,2-(L-Araf-β1,5-D-Dha-β1,3-)[ UA/D-GalAp]n

**Miscellaneous CDROs (CDRO_A_M#)**

1. L-Galp-α1,2-D-GlcA-β1,4(2-O-Me-D-Xylp-α1,3)-L-Fucp-α1,4-(D-GalAp-β1,3)(D-GalAp-α1,2)-L-Rhap-α1,3-D-Apif-β1,2-(L-Araf1,2-)-[D-GalAp]n
2. L-Galp-α1,2-D-GlcA-β1,4(2-O-Me-D-Xylp-α1,3)-L-Fucp-α1,4-(D-GalAp-β1,3)-L-Rhap-α1,3-D-Apif
3. **D-GlcA-β1,4(2-O-Me-D-Xylp-α1,3)-L-Fucp-α1,4-(D-GalAp-β1,3)-L-Rhap-α1,3-D-Apif**
4. L-Galp-α1,2-D-GlcA-β1,4(2-O-Me-D-Xylp-α1,3)-L-Fucp-α1,4-((3-O-Me-)-D-GalAp-β1,3)-L-Rhap-α1,3-D-Apif
5. D-GlcA-β1,4(2-O-Me-D-Xylp-α1,3)-L-Fucp-α1,4-((3-O-Me-)-D-GalAp-β1,3)-L-Rhap-α1,3-D-Apif
6. L-Galp-α1,2-D-GlcA-β1,4(2-O-Me-D-Xylp-α1,3)-L-Fucp-α1,4-((4-O-Me-)-D-GalAp-β1,3)-L-Rhap-α1,3-D-Apif
7. D-GlcA-β1,4(2-O-Me-D-Xylp-α1,3)-L-Fucp-α1,4-((4-O-Me-)-D-GalAp-β1,3)-L-Rhap-α1,3-D-Apif
8. L-Galp-α1,2-D-GlcA-β1,4(2-O-Me-D-Xylp-α1,3)-L-Fucp-α1,4-((3-O-Me-)(4-O-Me-)-D-GalAp-β1,3)-L-Rhap-α1,3-D-Apif
9. D-GlcA-β1,4(2-O-Me-D-Xylp-α1,3)-L-Fucp-α1,4-((3-O-Me-)(4-O-Me-)-D-GalAp-β1,3)-L-Rhap-α1,3-D-Apif
10. L-Galp-α1,2-D-GlcA-β1,4(2-O-Me-D-Xylp-α1,3)-L-Fucp-α1,4-(D-GalAp-β1,3)-L-Rhap-α1,3-D-Apif-β1,2-D-GalAp
11. D-GlcA-β1,4(2-O-Me-D-Xylp-α1,3)-L-Fucp-α1,4-(D-GalAp-β1,3)-L-Rhap-α1,3-D-Apif-β1,2-D-GalAp
12. L-Galp-α1,2-D-GlcA-β1,4(2-O-Me-D-Xylp-α1,3)-L-Fucp-α1,4-((3-O-Me-)-D-GalAp-β1,3)-L-Rhap-α1,3-D-Apif-β1,2-D-GalAp
13. D-GlcA-β1,4(2-O-Me-D-Xylp-α1,3)-L-Fucp-α1,4-((3-O-Me-)-D-GalAp-β1,3)-L-Rhap-α1,3-D-Apif-β1,2-D-GalAp
14. L-Galp-α1,2-D-GlcA-β1,4(2-O-Me-D-Xylp-α1,3)-L-Fucp-α1,4-((4-O-Me-)-D-GalAp-β1,3)-L-Rhap-α1,3-D-Apif-β1,2-D-GalAp
15. D-GlcA-β1,4(2-O-Me-D-Xylp-α1,3)-L-Fucp-α1,4-((4-O-Me-)-D-GalAp-β1,3)-L-Rhap-α1,3-D-Apif-β1,2-D-GalAp
16. L-Galp-α1,2-D-GlcA-β1,4(2-O-Me-D-Xylp-α1,3)-L-Fucp-α1,4-((3-O-Me-)(4-O-Me-)-D-GalAp-β1,3)-L-Rhap-α1,3-D-Apif-β1,2-D-GalAp
17. D-GlcA-β1,4(2-O-Me-D-Xylp-α1,3)-L-Fucp-α1,4-((3-O-Me-)(4-O-Me-)-D-GalAp-β1,3)-L-Rhap-α1,3-D-Apif-β1,2-D-GalAp
18. L-Galp-α1,2-D-GlcA-β1,4(2-O-Me-D-Xylp-α1,3)-L-Fucp-α1,4-(D-GalAp-β1,3)-L-Rhap-α1,3-D-Apif-β1,2-(L-Araf1,3-)-D-GalAp
19. D-GlcA-β1,4(2-O-Me-D-Xylp-α1,3)-L-Fucp-α1,4-(D-GalAp-β1,3)-L-Rhap-α1,3-D-Apif-β1,2-(L-Araf1,3-)-D-GalAp
20. L-Galp-α1,2-D-GlcA-β1,4(2-O-Me-D-Xylp-α1,3)-L-Fucp-α1,4-((3-O-Me-)-D-GalAp-β1,3)-L-Rhap-α1,3-D-Apif-β1,2-(L-Araf1,3-)-D-GalAp
21. D-GlcA-β1,4(2-O-Me-D-Xylp-α1,3)-L-Fucp-α1,4-((3-O-Me-)-D-GalAp-β1,3)-L-Rhap-α1,3-D-Apif-β1,2-(L-Araf1,3-)-D-GalAp
22. L-Galp-α1,2-D-GlcA-β1,4(2-O-Me-D-Xylp-α1,3)-L-Fucp-α1,4-((4-O-Me-)-D-GalAp-β1,3)-L-Rhap-α1,3-D-Apif-β1,2-(L-Araf1,3-)-D-GalAp
23. D-GlcA-β1,4(2-O-Me-D-Xylp-α1,3)-L-Fucp-α1,4-((4-O-Me-)-D-GalAp-β1,3)-L-Rhap-α1,3-D-Apif-β1,2-(L-Araf1,3-)-D-GalAp
24. L-Galp-α1,2-D-GlcA-β1,4(2-O-Me-D-Xylp-α1,3)-L-Fucp-α1,4-((3-O-Me-)(4-O-Me-)-D-GalAp-β1,3)-L-Rhap-α1,3-D-Apif-β1,2-(L-Araf1,3-)-D-GalAp
25. D-GlcA-β1,4(2-O-Me-D-Xylp-α1,3)-L-Fucp-α1,4-((3-O-Me-)(4-O-Me-)-D-GalAp-β1,3)-L-Rhap-α1,3-D-Apif-β1,2-(L-Araf1,3-)-D-GalAp
26. 6-O-Me-D-GlcA-β1,4(2-O-Me-D-Xylp-α1,3)-L-Fucp-α1,4-(D-GalAp-β1,3)-L-Rhap-α1,3-D-Apif
27. L-Rhap-α1,3-D-Apif-β1,2-(6-O-Me-D-GalAp-α1,4-)(L-Araf1,3-)-D-GalAp-α1,2-D-GalAp
28. 6-O-Me-D-GlcA-β1,4(2-O-Me-D-Xylp-α1,3)-L-Fucp-α1,4-(D-GalAp-β1,3)(D-GalAp-α1,2)-L-Rhap-α1,-3-D-Apif
29. 6-O-Me-D-GlcA-β1,4(2-O-Me-D-Xylp-α1,3)-L-Fucp-α1,4-(D-GalAp-β1,3)-L-Rhap-α1,-3- D-Apif-β1,2-D-GalAp
30. 6-O-Me-D-GlcA-β1,4(2-O-Me-D-Xylp-α1,3)-L-Fucp-α1,4-(D-GalAp-β1,3)-L-Rhap-α1,3-D-Apif-β1,2- (L-Araf1,3-)-D-GalAp
