## Supplemental table 2 for "Towards a comprehensive chemical and genetic tool library for rhamnogalacturonan-II oligosaccharides and exploitation"

**Supplemental table 2: CDROs and corresponding genes in the *B. theta* RG-II PUL that can be targeted through genetic engineering to produce them.** Data is based on screens of *B. theta* transposon library and newly generated knockout (KO) mutant strains. * denotes abundant CDROs generated for the specific strain, ✓: yes (available or detected), ⊗ : (no) not available or not detected, n/a: not applicable, TLC = thin layer chromatography, HPAEC = high performance anion exchange chromatography, MS = Mass spectrometry, Tbc= to be confirmed. Percentage yield is the recovered amount of CDRO expressed as a percentage of the original starting material (typically 100 mg of starting RG-II substrate).

| Tranposon mutant ID | Gene | Strains available | | CDRO detection method | | | CDROs generated | CDRO yield % total RG-II | Study |
|---|---|---|---|---|---|---|---|---|---|
| | | Tn | KO | TLC | HPAEC | MS | | | |
| P5-C6 | **BT0980** | ✓ | ⊗ | ⊗ | ⊗ | ⊗ | not detected | n/a | This study |
| P1-B8 | **BT0981** | ✓ | ✓ | ⊗ | ⊗ | ⊗ | not detected | n/a | This study |
| n/a | **BT0983** | ⊗ | ⊗ | n/a | n/a | n/a | n/a | n/a | This study |
| P21-F8 | **BT0984** | ✓ | ✓ | ✓ | ✓ | ✓ | CDRO_B5*, CDRO_B21/31*, CDRO_B13/18, CDRO_B10/B15 | >5% | This study |
| n/a | **BT0985** | ⊗ | ✓ | ✓ | ✓ | ✓ | CDRO_B4*, CDRO_B12, CDRO_B17 | >5% | This study |
| P21-E11 | **BT0986** | ✓ | ✓ | ✓ | ✓ | ✓ | CDRO_B3*, CDRO_B11*, CDRO_B23 | >5% | This study and Ndeh et al., 2017 |
| n/a | **BT0992** | ⊗ | ✓ | ✓ | ✓ | ✓ | CDRO_B5* | >5% | Ndeh et al., 2017 |
| P11-D8 | **BT0993** | ✓ | ⊗ | ✓ | ✓ | ✓ | CDRO_B6*, CDRO_B19 | >5% | This study |
| P13-B7 | **BT0994** | ✓ | ⊗ | ⊗ | ⊗ | ⊗ | not detected | n/a | This study |
| P19-B9 | **BT0996** | ✓ | ✓ | ✓ | ✓ | ✓ | CDRO_A2*, CDRO_A_M3*, CDRO_A27*, CDRO_A20*, CDRO_B1, B9/14 | >5% | This study |
| P29-C10 | **BT0997** | ✓ | ✓ | ✓ | ✓ | ✓ | CDRO_A3*, CDRO_A10, CDRO_A14 | >5% | This study and Ndeh et al., 2017 |
| P4-G10 | **BT0998** | ✓ | ⊗ | ⊗ | ⊗ | ⊗ | not detected | n/a | This study |
| P6-G8 | **BT1000** | ✓ | ⊗ | ⊗ | ⊗ | ⊗ | not detected | n/a | This study |
| P16-A7 | **BT1001** | ✓ | ✓ | ✓ | ✓ | ✓ | CDRO_A7*/ CDRO_B8* | >5% | This study and Ndeh et al., 2017 |
| P22-G5 | **BT1002** | ✓ | ✓ | ✓ | ✓ | ✓ | CDRO_A5* | >5% | This study and Ndeh et al., 2017 |
| P32-C09 | **BT1003** | ✓ | ✓ | ✓ | ✓ | ✓ | CDROB7* | >5% | This study and Ndeh et al., 2017 |
| P32-F08 | **BT1010** | ✓ | ✓ | ✓ | ✓ | ✓ | CDRO_A1* CDRO_A20*, CDRO_M2, CDRO_A8/CDRO_A12 | >5% | This study |
| P2-E7 | **BT1011** | ✓ | ⊗ | ✓ | ✓ | ✓ | tbc | >5% | This study |
| n/a | **BT1012** | ⊗ | ✓ | ✓ | ✓ | ✓ | CDRO_B33/A26*, CDRO_B58, CDRO_B83, CDRO_B108 | >5% | This study |
| P5-B6 | **BT1013** | ✓ | ✓ | ✓ | ✓ | nt | tbc | >5% | This study |
| n/a | **BT1017** | ⊗ | ✓ | ✓ | ✓ | ✓ | CDRO_A-F64, CDRO_A-F66, CDRO_M26-M30 | >5% | This study and Ndeh et al., 2017 |
| n/a | **BT1018** | ⊗ | ⊗ | n/a | n/a | n/a | n/a | n/a | n/a |
| P34-F05 | **BT1019** | ✓ | ✓ | ✓ | ✓ | ✓ | CDRO_B1*, CDRO_B2* | >5% | This study |
| P22-E1 | **BT1020** | ✓ | ✓ | ✓ | ✓ | ✓ | CDRO-D/B7*, others tbc | >5% | This study and Ndeh et al., 2017 |
| n/a | **BT1021** | ✓ | ✓ | ✓ | ✓ | ✓ | CDRO_AF45 | >5% | Ndeh et al., 2017 |
| n/a | **BT1022** | ⊗ | ⊗ | n/a | n/a | n/a | n/a | n/a | n/a |
| P13-H8 | **BT1023** | ✓ | ✓ | ⊗ | ⊗ | ⊗ | not detected | n/a | Ndeh et al., 2017 |
| P23-F7 | **BT1024** | ✓ | ⊗ | ⊗ | ⊗ | ⊗ | not detected | n/a | This study |
| P26-C7 | **BT1025** | ✓ | ⊗ | ⊗ | ⊗ | ⊗ | not detected | n/a | This study |
| P17-E6 | **BT1026** | ✓ | ⊗ | ⊗ | ⊗ | ⊗ | not detected | n/a | This study |
| P29-E11 | **BT1028** | ✓ | ⊗ | ⊗ | ⊗ | ⊗ | not detected | n/a | This study |
| P7-E2 | **BT1029** | ✓ | ⊗ | ⊗ | ⊗ | ⊗ | not detected | n/a | This study |
| P29-F3 | **BT1030** | ✓ | ✓ | ⊗ | ⊗ | ⊗ | not detected | n/a | This study |
| P32-A06 | **BT1031** | ✓ | ⊗ | ⊗ | ⊗ | ⊗ | not detected | n/a | This study |
| P22-E9 | **BT3662** | ✓ | ⊗ | ⊗ | ⊗ | ⊗ | not detected | n/a | This study |
| P22-C1 | **BT3663** | ✓ | ⊗ | ⊗ | ⊗ | ⊗ | not detected | n/a | This study |
| P29-G12 | **BT3664** | ✓ | ⊗ | ⊗ | ⊗ | ⊗ | not detected | n/a | This study |
| P27-E9 | **BT3665** | ✓ | ✓ | ⊗ | ⊗ | ⊗ | not detected | n/a | This study |
| P27-E4 | **BT3670** | ✓ | ⊗ | ⊗ | ⊗ | ⊗ | not detected | n/a | This study |
| P3-D3 | **BT3671** | ✓ | ⊗ | ⊗ | ⊗ | ⊗ | not detected | n/a | This study |
| P8-C6 | **BT3672** | ✓ | ⊗ | ⊗ | ⊗ | ⊗ | not detected | n/a | This study |
| P11-B5 | **BT1683** | ✓ | ⊗ | ⊗ | ⊗ | ⊗ | not detected | n/a | This study |
| n/a | **BT0984/BT1012** | ⊗ | ✓ | ✓ | ✓ | ✓ | CDRO_B30 *, CDRO_B13, CDRO_B18, CDRO_B25, CDRO_B4, B10/B15 | >5% | This study |
| n/a | **BT1021/BT1010** | ⊗ | ✓ | ✓ | ✓ | ✓ | CDRO_AF39, CDRO_M10 | >5% | This study and Ndeh et al., 2017 |
| n/a | ***none (B. theta Wt)*** | n/a | n/a | ✓ | ✓ | nt | CDRO_A6*, CDROB7* | >5% | This study and Ndeh et al., 2017 |
